# Whole genome assembly of the Little Millet (*Panicum sumatrense*) genome: A climate resilient crop species

**DOI:** 10.1101/2024.02.26.582036

**Authors:** Rasmita Rani Das, Ashish Kumar, Dushyant Singh Baghel, Ajay K Mahato, Nrisingha Dey, Pulok K Mukherjee, Seema Pradhan

## Abstract

Little millet (*Panicum sumatrense*) belongs to one of the largest genera, *Panicum* comprising of 500 species that are distributed especially in tropical and sub-tropical Asia as well as Africa. This is a climate resilient crop that has the ability to adapt to adverse growing conditions, especially drought. Added to this is the better nutritional profile of little millet, which has higher iron and fibre contents as compared with rice. These characteristics make it an important crop species that would be pivotal for ensuring food security. Generating genomic resources for the species is significant and will have implications in crop improvement. Therefore, in the present study we report the *de novo* whole genome sequence of *P. sumatrense*. Long read sequencing and Hi-C based scaffolding resulted in a total of 279 scaffolds with N50 of 7.8Mb. The genome was annotated to predict protein coding genes and orthologous groups were analysed after comparison with various plant genomes. This high-quality genome assembly can be a valuable resource as future reference for genomic studies in this crop and related crop species.

## Background

*Panicum sumatrense* (Little Millet) is a small seeded cereal crop mostly grown in the semi-arid tropical and sub-tropical regions of Asia and Africa. The genus *Panicum* L., comprising of about 500 species are distributed worldwide in tropical and subtropical regions, and is one of the largest genera of the family Poaceae **[1]**. *P. sumatrense* is a tetraploid (2n = 4x = 36) **[2]** crop species and has some additional benefits due to their unique ability to grow in unfavorable ecological conditions like various abiotic stresses including adverse environments such as soil with high salinity and high temperatures **[3]**. Along with its ability to grow under unfavorable conditions, the species is highly nutritious with higher content of nutrients such as Iron and Fiber as compared to other staple crops like Rice and Wheat **[4,5]**.

This nutri-cereal is a good source of food and feed across India and sub-Saharan Africa. The crop is consumed as a staple food in drought prone Indian states like Rajasthan, parts of Odisha, and is popularly known as “Indian Millet” **[6]**. Around 5,000 years ago, little millet was domesticated in India **[7]** and is grown mainly in certain Asian countries like India, Myanmar, Nepal and Sri Lanka **[8]**. The Indian little millet has a short crop cycle and can efficiently grow in both dry and waterlogged conditions with adverse environments such as soil with high salinity and temperatures. Its consistency in providing better yield in drought prone marginal lands enhances its importance as a crop for regional food stability and nutritional security. This versatile crop with better nutritional contents and health benefits with similar compositions as other cereal crops can be a worthy addition to the diet. Despite its nutritional, economical values, the crop is limited with genomic resources. In an attempt to generate some genetic database of little millet, our previous study tried to explore the transcriptome profile with the help of next-generation sequencing **[6]**. Although the chloroplast genome of little millet has earlier been reported **[9]**, the whole genome remains to be sequenced yet.

We have previously reported the de novo transcriptome assembly of *P. sumatrense* cv. OLM 20 **[6]**, and now have selected the same variety for whole genome sequencing and assembly with the help of third generation sequencing technology. With advent of high-throughput sequencing technology, the third-generation sequencing platforms have become new tools of genome research. These contemporary sequencing platforms circumvent some limitations posed by the second-generation technology such as reads being too short and may match with many different regions of the genome and are not unique to any specific region of the sequence. The significantly longer reads from platforms like the PacBio sequel II, have made genome assemblies comparatively error free and especially effective for handling polyploid genomes **[10-11]**. In this study, we present the first report of the whole genome assembly of *P. sumatrense*.

## Methods

### Sampling and Sequencing

The seeds from a single plant of *P. sumatrense* OLM 20 were obtained from CPR, Berhampur, India (19° 18’ 53.8632’’ N, 84° 47’ 38.7240’’ E). They were grown in soil (local sandy loam soil) in controlled conditions (light/dark:16hr/8hr; 55% RH, 28°C ±2 temperature). Young leaves were collected from the plants at 5 leaf stage (Supplementary figure S1), surface sterilized with 70% ethanol and stored at -80°C after freezing in liquid N_2_. High molecular weight genomic DNA was isolated gDNA was isolated with QIAGEN 100G kit by following manufacturers protocol. DNA purity, size and concentration was recorded by using NanoDrop 2000 (Thermo Scientific), FEMTO Pulse (Agilent) and Qubit 3 fluorometer (Invitrogen). Good quality, high molecular weight genomic DNA was used for library preparation and sequencing. High molecular weight genomic DNA was sheared mechanically on Megaruptor 3 (Diagenode) to generate 30 Kb fragments. These fragments were used to prepare the library using SMRTBell Express template Preparation Kit2.0 (Pacific Biosciences) following manufacturer’s protocol. The libraries were purified using AMPure PB beads (PACBIO) and size selected using BluePippin (Sage science). The quality filtered libraries were sequenced on PacBio Sequel II platform to generate a total of 317 GB of unique molecular yield data. Then the high molecular weight DNA was purified and used for library preparation for generating 1.01 GB of assembly HiC data using Proximity Ligation technology workflow (Phase Genomics)

### *De novo* genome assembly

Initially, the PacBio reads were screened for generation of Hifi reads using CCS. A primary genome assembly was generated using hifiasm with the option to integrate Hi-C reads (v0.16.1 r375) **[12]**. HiFi reads were aligned to the assembled genome using minimap2 (v2.24 r1122) **[13]** to create a coverage histogram. This coverage histogram was used to purge haplotigs from the assembly using purge_haplotigs (v1.1.2) **[14]** and the contigs were collapsed based on this purging. HiC reads were aligned to this contig level assembly using BWA (v 0.7.17 r1188) **[15]**. Each pair was aligned separately to the assembly. The alignments were processed by removing chimeric alignments and duplicates followed by a MAPQ filter of 30 to retain only the best alignments. Scaffolding of the contigs was done using the SALSA pipeline (v2.3) which performs large scaffold construction using graph construction and link scoring function **[16]**. Misjoin, detection and correction was performed repeatedly until the best final scaffolds were achieved. Contig and scaffold level assemblies were assessed for completeness using BUSCO (v5.3.1) **[17]** and assembly metrics were calculated using QUAST (v5.2.0) **[18]**. The final assembly (after scaffolding with Hi-C data) consisted of scaffolds with an average length of 2.03Mb (Table 1). The N50 value for assembled contigs/scaffolds was calculated to be 7.85 Mb, which is better than those reported for other millets species (Table 2).

**Table 1:**
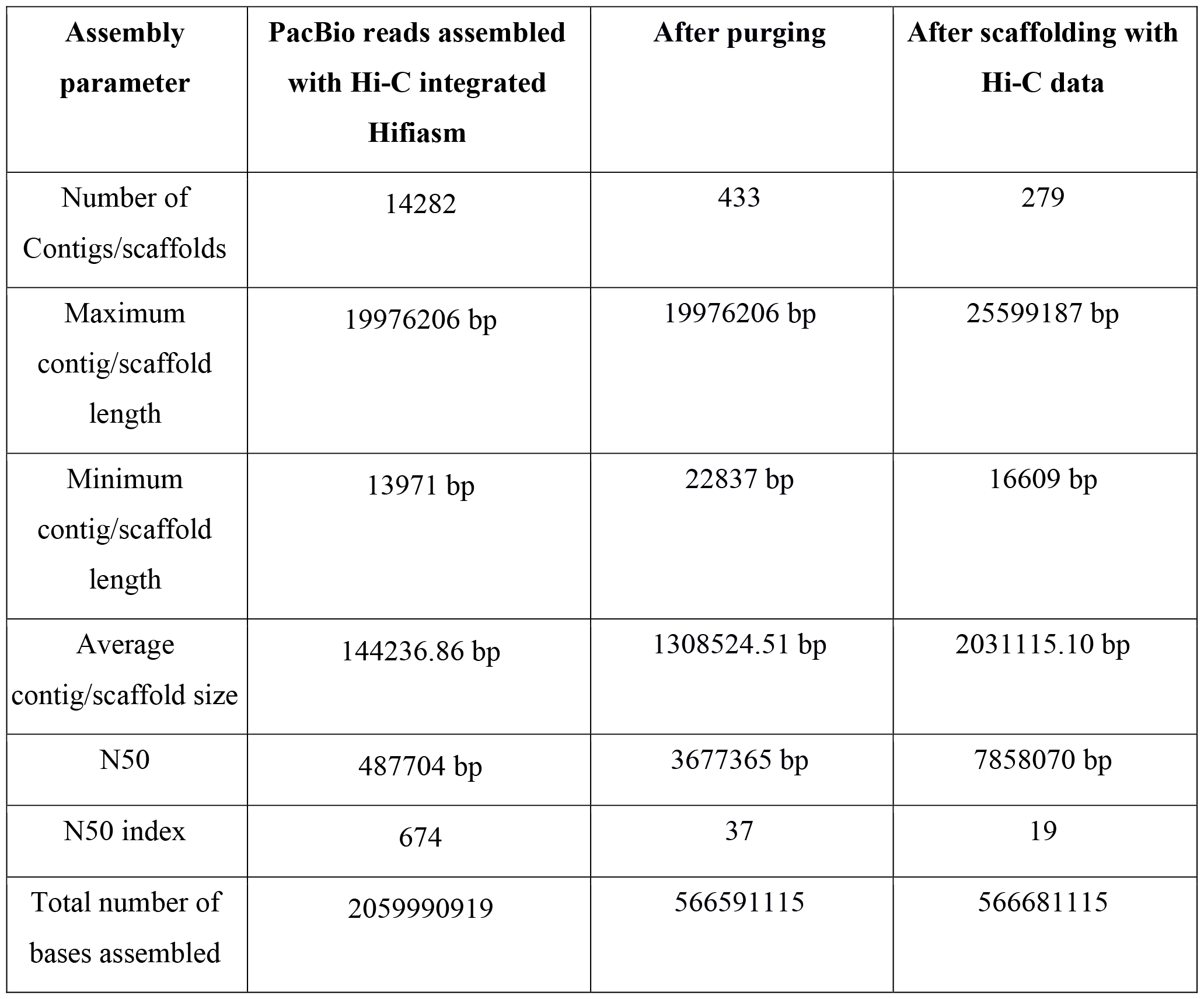
Assembly statistics after primary assembly with Hifiasm and improved assembly after Hi-C sequencing.

**Table 2:**
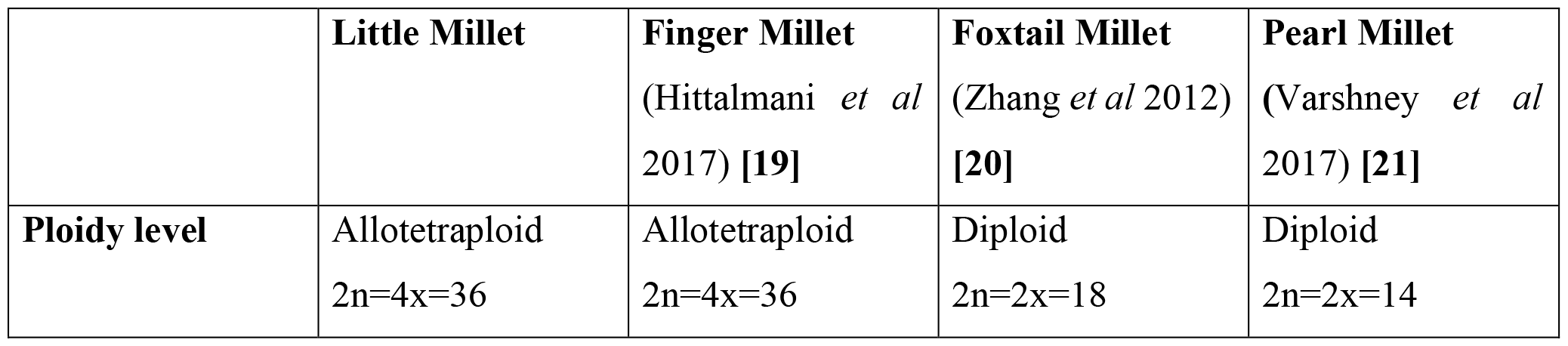
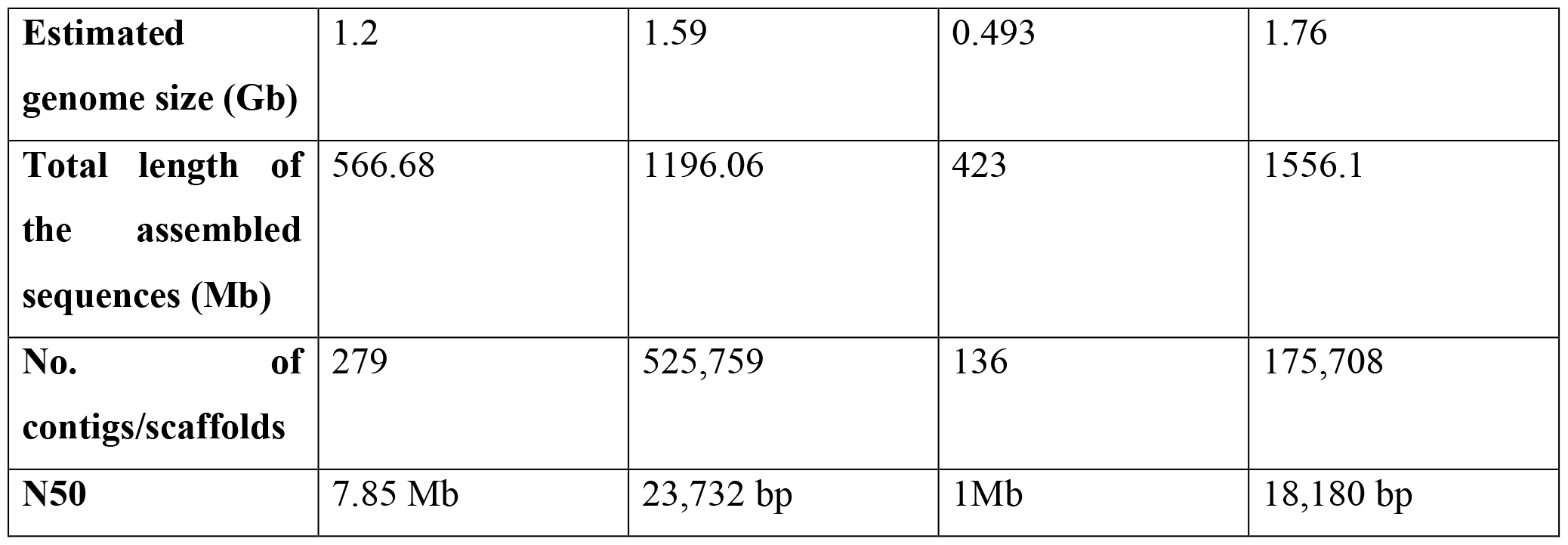
Comparison with other millet genomes.

|  | Little Millet | Finger Millet<br>(Hittalmani *et al* 2017) [19] | Foxtail Millet<br>(Zhang *et al* 2012) [20] | Pearl Millet<br>(Varshney *et al* 2017) [21] |
| --- | --- | --- | --- | --- |
| **Ploidy level** | Allotetraploid<br>2n=4x=36 | Allotetraploid<br>2n=4x=36 | Diploid<br>2n=2x=18 | Diploid<br>2n=2x=14 |
| **Estimated genome size (Gb)** | 1.2 | 1.59 | 0.493 | 1.76 |
| **Total length of the assembled sequences (Mb)** | 566.68 | 1196.06 | 423 | 1556.1 |
| **No. of contigs/scaffolds** | 279 | 525,759 | 136 | 175,708 |
| **N50** | 7.85 Mb | 23,732 bp | 1Mb | 18,180 bp |

### Genome quality assessment

The final assembly generated after scaffolding with HiC was assessed for quality using BUSCO (v5.3.1) **[17]**. We found a total of 4690 complete BUSCOs out of a total of 4896 BUSCOs from the poales_odb10 database to be represented in this assembly of little millet genome (Figure 1). We also mapped the reads from RNA-Seq of little millet onto the assembled genome and found that, on an average, more than 90% of the reads aligned to the genome (Figure 2). The LTRs identified through LTR_harvest **[19]** were analysed with LTR_retriever [https://github.com/oushujun/LTR_retriever] to obtain the LAI (LTR Assembly Index). The LAI for the whole genome assembly of *P. sumatrense* was found to be 18.26 (Supplementary data). The data from all the quality assessment methods/tools suggests that this assembly of the little millet genome is of a superior quality.

**Figure 1.**
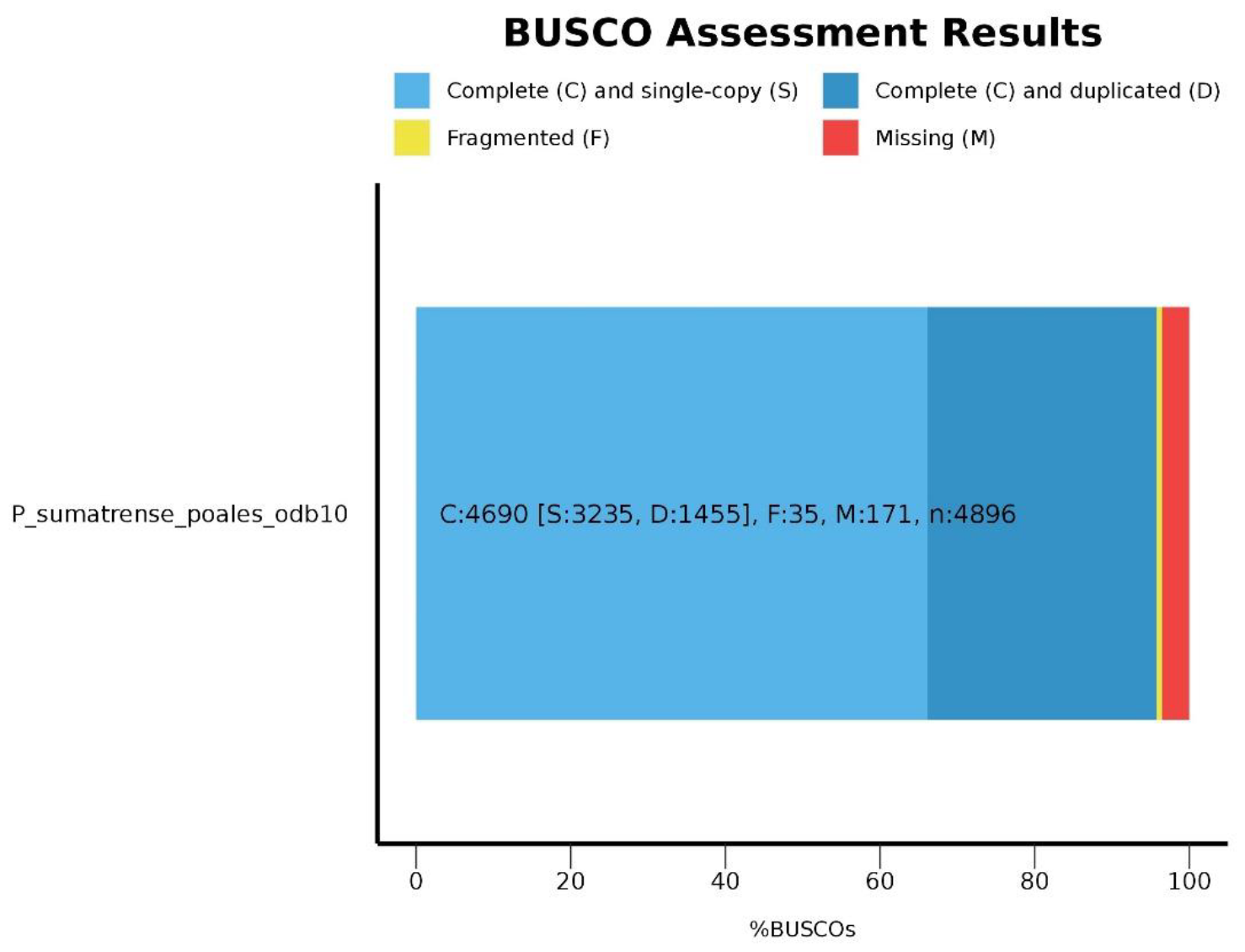
BUSCO assessment results for little millet genome assembly using Poales_odb10 database (C:95.8%[S:66.1%, D:29.7%],F:0.7%,M:3.5%,n:4896)

**Figure 2.**
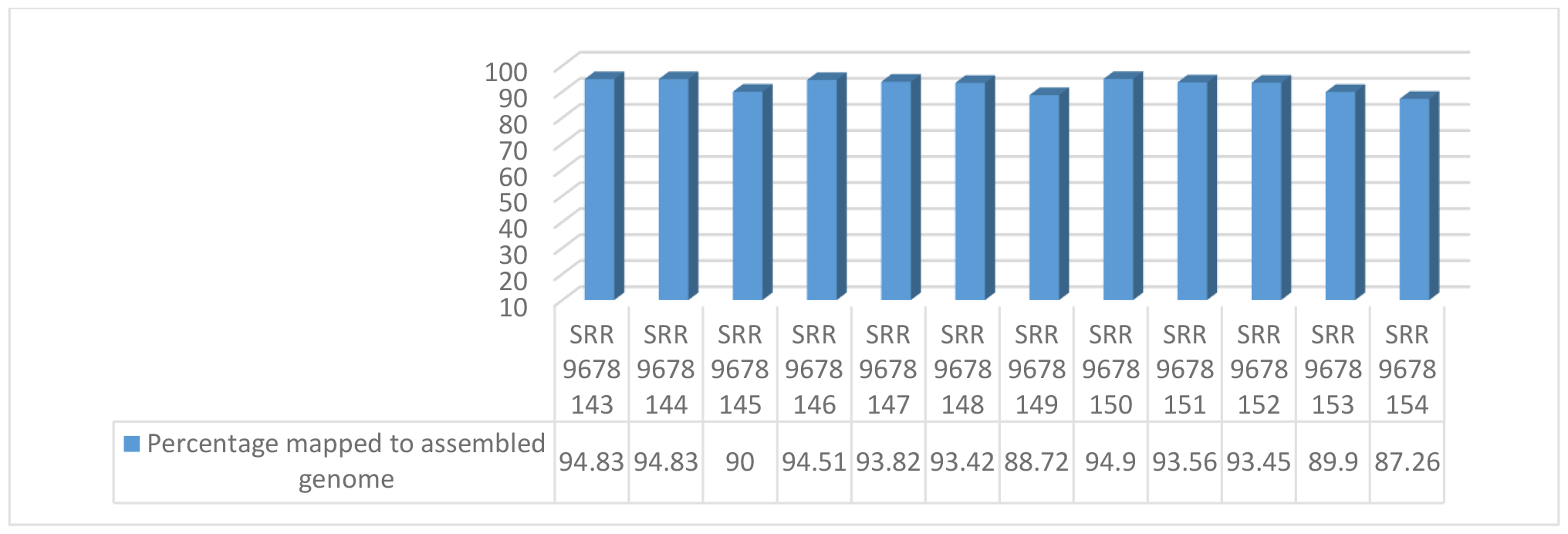
Percentage of reads from RNA-Seq data of Little millet in NCBI mapped onto the assembled genome.

### Repeat elements and Genome annotation

RepeatModeler was also used to generate a customised repeat database for little millet. The resulting customized repeat database consisting of a total of 8950 sequences was used to screen the assembled genome using RepeatMasker program (v4.1.1, http://www.repeatmasker.org/) that screens DNA sequences for retroelements, interspersed repeats, transposons and low complexity DNA sequences (Table 3). Most of the repeat elements were “retroelements” (30.05%) and LTR elements were found to be the most abundant type of retroelements (29.05%). The GC content was found to be 45.86% and about 54% bases were masked. The protein sequences reported in *Panicum hallii* (https://plants.ensembl.org/Panicum_hallii/Info/Index) were used to predict the protein coding sequences in *P. sumatrense* using BRAKER (https://github.com/Gaius-Augustus/BRAKER; **[20]**). This led to the prediction of 49,690 protein coding sequences in the genome of little millet, of which 43,961 sequences (88.4%) could be annotated with the Uniprot SwissProt database.

**Table 3:** Statistics of genome repeat masking.

|  |  |  |  |
| --- | --- | --- | --- |
| Sequences: 279 |  |  |  |
| Total length: 566681115 bp |  |  |  |
| GC level: 45.86% |  |  |  |
| Bases masked: 308857442 bp (54.50%) |  |  |  |
|  | **Number of elements** | **Length occupied** | **Percentage of sequence** |
| **Retroelements** | **85473** | **170312443 bp** | **30.05%** |
| SINEs | 0 | 0 bp | 0.00% |
| Penelope | 0 | 0 bp | 0.00% |
| LINEs | 11734 | 5670933 bp | 1.00% |
| CRE/SLACS | 0 | 0 bp | 0.00% |
| L2/CR1/Rex | 0 | 0 bp | 0.00% |
| R1/LOA/Jockey | 0 | 0 bp | 0.00% |
| R2/R4/NeSL | 0 | 0 bp | 0.00% |
| RTE/Bov-B | 757 | 259125 bp | 0.05% |
| L1/CIN4 | 10977 | 5411808 bp | 0.96% |
| LTR elements | 73739 | 164641510 bp | 29.05% |
| BEL/Pao | 0 | 0 bp | 0.00% |
| Ty1/Copia | 16449 | 28221702 bp | 4.98% |
| Gypsy/DIRS1 | 57222 | 136364159 bp | 24.06% |
| Retroviral | 0 | 0 bp | 0.00% |
| **DNA transposons** | **16273** | **10827300 bp** | **1.91%** |
| hobo-Activator | 2712 | 1540593 bp | 0.27% |
| Tc1-IS630-Pogo | 302 | 211507 bp | 0.04% |
| En-Spm | 0 | 0 bp | 0.00% |
| MuDR-IS905 | 0 | 0 bp | 0.00% |
| PiggyBac | 0 | 0 bp | 0.00% |
| Tourist/Harbinger | 1734 | 704105 bp | 0.12% |
| Other | 0 | 0 bp | 0.00% |
| **Rolling-circles** | **1600** | **903838 bp** | **0.16%** |
| **Unclassified** | **293564** | **121897832 bp** | **21.51%** |
| **Total interspersed repeats** | **--** | **303037575 bp** | **53.48%** |
| **Small RNA** | **0** | **0 bp** | **0.00%** |
| **Satellites** | **0** | **0 bp** | **0.00%** |
| **Simple repeats** | **93833** | **4403677 bp** | **0.78%** |
| **Low complexity** | **10280** | **512352 bp** | **0.09%** |

### Gene expression under drought and salinity stress

Analysing the orthologous groups present in plants that may be closely or distantly related provide insights into the probable evolution of a plant. The protein sequences predicted from the whole genome of *Panicum sumatrense* were compared with those from 8 other plants; *Oryza sativa (indica), Brachypodium distachyon, Eleusine coracana, Panicum hallii, Panicum miliaceum, Panicum virgatum, Pennisetum glaucum*, and *Setaria italica* using OrthoFinder **[21]** to identify the orthologous groups shared by and specific to these plants. The analysis showed that a total of 442873 genes were assigned to 55,120 orthogroups (Supplementary data). Of these, 1834 orthogroups were specific to *P. sumatrense* (Supplementary data). The genes present in these little millet specific orthologous groups were grouped into various GO terms to get an idea about what distinguishes little millet from its relatives in the poaceae family. It was observed that processes like hydrolase (carbon-nitrogen bonds) activity, glucosidase activity and oxidoreductase activity under “GO Slim-Molecular function” category and processes related to metabolism of phosphate containing compounds and organonitrogen compounds under the “GO Slim-Biological processes” category were enriched in these orthologous groups.

Further, a gene expression atlas was generated after mapping the RNA-seq reads from a previous study **[6]** onto the complete set of protein coding genes predicted from the whole genome of little millet to identify differentially expressed genes in response to drought and salinity stresses. Briefly, the clean reads from each sample were mapped onto the genes using Bowtie2 **[22]** and it was observed that genes encoding LEA proteins (g4028.t1), Dehydrin (g12235.t1), seed maturation protein (g14378.t1), F-box proteins (g43661.t1, g6953.t1), E3 ubiquitin ligases (g11981.t2), pyruvate dehydrogenase (g39238.t1) and transcription factors encoding MYB (g31522.t1) and MADS (g29003.t1) TFs were upregulated in leaf tissue of little millet under conditions of drought and salinity stress (Figure 3A, Supplementary data). A similar analysis in root tissue led to an interesting observation where most of the genes were found to be downregulated in response to drought and salinity stress as compared with the control sample. This is in stark contrast to the results obtained in case of leaf tissue and showed a marked downregulation of genes encoding proteins like Early nodulin (g38606.t1), L-lactate dehydrogenase (g3894.t1), Stearoyl-[acyl-carrier-protein]9-desaturase (g6357.t1), Alcohol dehydrogenase (g20376.t1), heat shock protein (g4349.t1). On the other hand, genes coding for proteins like F-box/kelch-repeat protein (g1251.t1), WRKY transcription factor (g42302.t2), Asparagine synthetase (g25249.t2) and UDP-glycosyltransferase (g19604.t1) were upregulated in root affected by salinity stress while that coding for a probable glycosyltransferase (g35695.t2) was upregulated in root in response to drought stress (Figure 3B, Supplementary data).

**Figure 3A.**
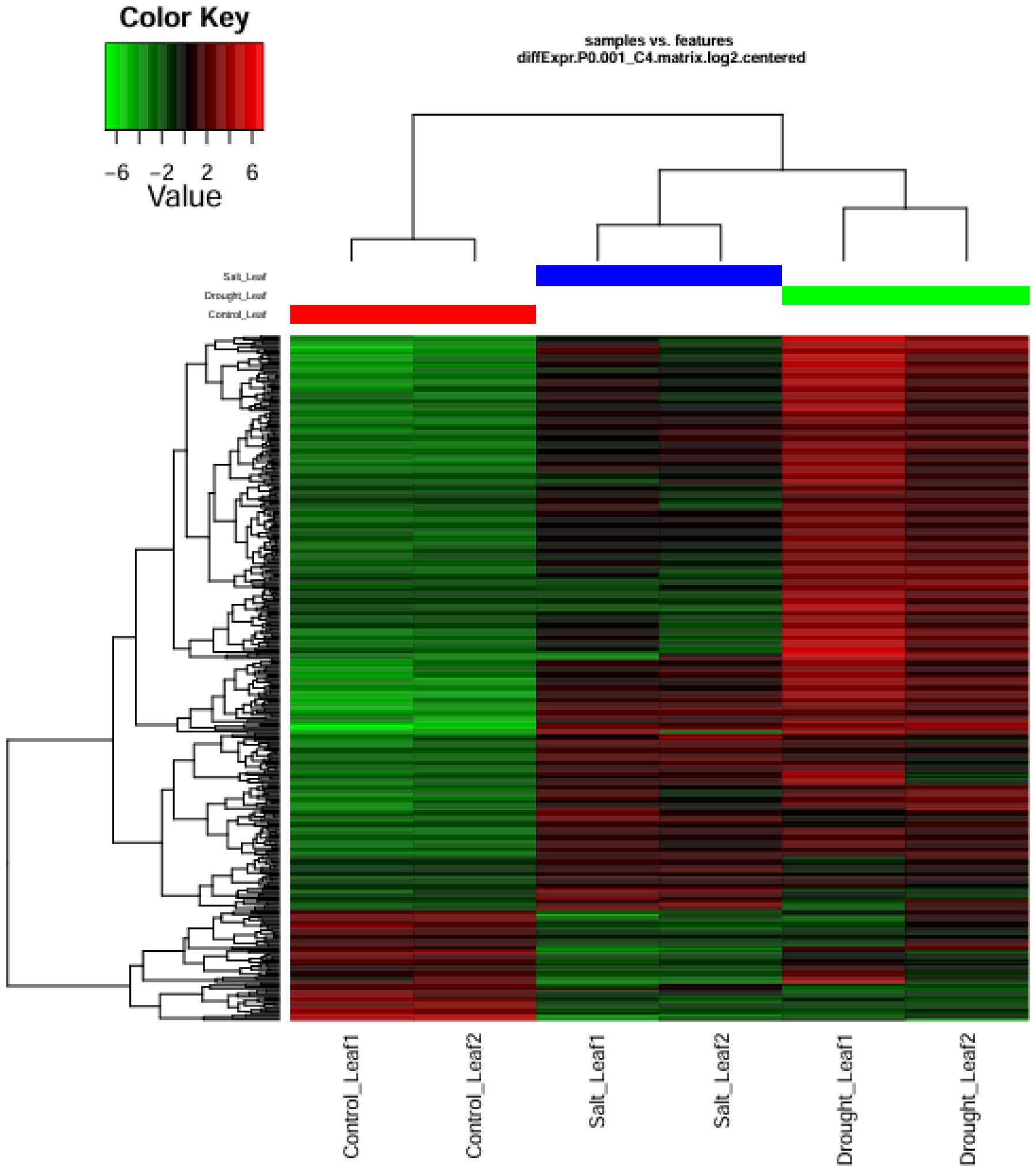
Gene expression profile for *P. sumatrense* leaf tissue in response to drought and salinity stress.

**Figure 3B.**
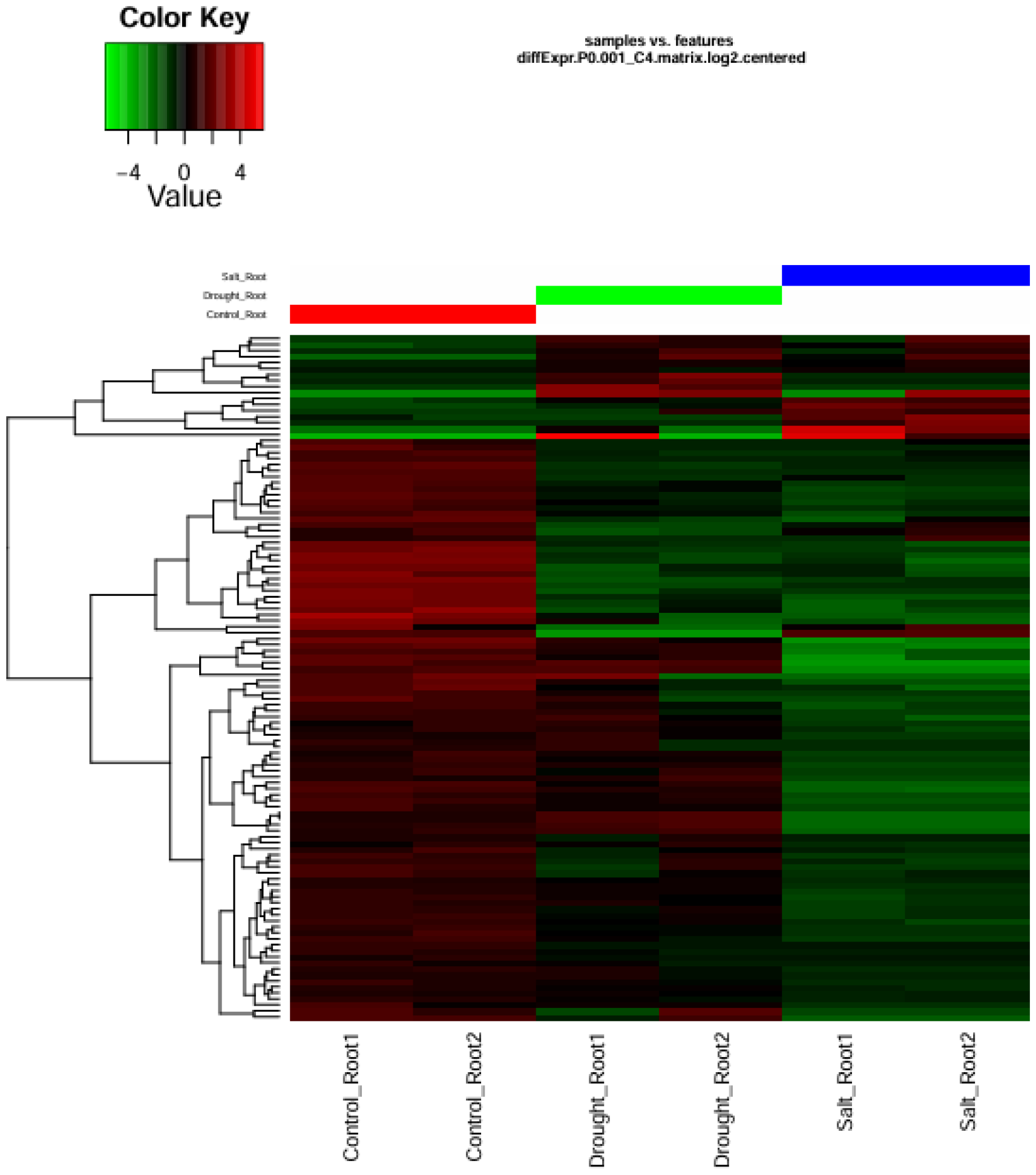
Gene expression profile for *P. sumatrense* root tissue in response to drought and salinity stress.

### SSRs in the little millet genome

Simple sequence repeats (SSRs) are one of the most valuable molecular markers with varied applications, especially in determining genetic diversity and plant breeding. So, MISA software (https://webblast.ipk-gatersleben.de/misa/) was used to mine SSRs in the little millet genome with the following parameters: Unit size / minimum number of repeats: (1/10) (2/6) (3/4) (4/4) (5/3) (6/3); Maximal number of bases interrupting 2 SSRs in a compound microsatellite:100. A total of 79,830 SSRs were identified in 270 sequences (Table 4). Majority of these were mono-nucleotide repeats (42.5%) and tri-nucleotide repeats (34.4%).

**Table 4:** SSRs identified in the genome sequence of *P. sumatrense*.

| Results of Microsatellite search | Value | Unit size | Number of SSRs |
| --- | --- | --- | --- |
| Total number of sequences examined | 279 | 1 | 33942 |
| Total size of examined sequences (bp) | 566681115 | 2 | 5656 |
| Total number of identified SSRs | 79830 | 3 | 27481 |
| Number of SSR containing sequences | 270 | 4 | 2425 |
| Number of sequences containing more than 1 SSR | 265 | 5 | 7217 |
| Number of SSRs present in compound formation | 4690 | 6 | 3109 |

## Discussion

Millets have gained significant popularity, especially in the last few decades, owing to their superior nutritive value and climate resilience, making them a viable food source for future food security. These are also a potential resource for identification of molecular markers and genes that could further the process of crop improvement. However, these endeavours are limited due to a lack of genomic resources in millets. While there are reports on genomic resources of other minor millet species like finger millet **[23]**, foxtail millet **[24]**, pearl millet **[25]**, such studies on little millet are very limited. The large and polyploid genome size of this plant poses certain difficulties in whole genome assembly. This study is an attempt to assemble the complex polyploid genome of *P. sumatrense* which generated a high-quality whole genome assembly for this tetraploid plant species. The good quality of the little millet assembly was evident after comparison with other millet counterparts and reiterated by more than 95% of BUSCOs being represented in this genome assembly and more than 90% of short reads from RNA-Seq being mapped onto the assembled genome **[6]**. The 49,690 predicted genes coding for viable protein products in the little millet genome could serve as a source of genic molecular markers, which in turn would be valuable for marker assisted selections.

*P. sumatrense* is a marginally cultivated cereal crop that can withstand conditions of drought and salinity **[26]**. Thus, studying the pattern of gene expression in this plant could help identify genes that may impart tolerance to these devastating stresses that lead to heavy losses in susceptible crops like rice. Similar analysis has been reported earlier for *P. sumatrense* in a study of the transcriptome of the plant in response to drought and salinity stress **[6]**. However, gene models that arise out of whole genome assembly, provide a more holistic dataset for generating gene expression atlas. The comparison of these protein sequences encoded in the little millet genome with those from other plants provided us with insights into the evolutionary process that could have contributed to the ability of this plant to withstand significant levels of abiotic stress. The observations pointed to the involvement of processes regulating oxidation-reduction and nitrogen and phosphate compound metabolism in the plant’s response to abiotic stresses. The gene expression data also indicated the role of well characterised genes such as LEA, F-box, dehydrin etc. in combating drought and salinity stress. The activity of these genes could be a critical factor contributing to plant growth in difficult terrain and provide a source of nutrition to marginalised population. This study will be a substantial contribution to the field of plant genomics and will serve as useful genomic resources which could contribute to future crop breeding and enhancement. However, this is a first draft of the whole genome assembly of *Panicum sumatrense* and we are curating it to generate an improved reference-grade genome assembly for this valuable crop plant.

## Additional files

**See Supplementary data**

## Abbreviations

Bp: base pairs
Mb: mega-base pairs
Gb: giga-base pairs
PacBio: Pacific Biosciences
BUSCO: Benchmarking Universal Single-Copy Orthologs
BWA: Burrows-Wheeler Aligner
LTR: long terminal repeat
NCBI: National Center for Biotechnology Information
BLAST: Basic Local Alignment Search Tool
SSR: Simple Sequence Repeat

## Authors’ contributions

R.R.D collected the plant materials; R.R.D, S.P, D.S.B, A.K. designed the project; S.P., D.S.B, A.K and A.K.M worked on sequencing and data analysing; R.R.D, P.K.M, N.D and S.P wrote the manuscript; All the authors revised and approved the final version of the manuscript.

## Acknowledgements

All authors acknowledge the contributions of Late Dr. Ajay Parida, who initiated the study and inspired everyone for the execution of the study. RRD is thankful to Department of Science and Technology (DST-INSPIRE), India for her fellowship. The study was carried out with core funds provided by Department of Biotechnology, Ministry of Science and Technology, Government of India.

