## Supplementary data for "Whole genome assembly of the Little Millet (*Panicum sumatrense*) genome: A climate resilient crop species"

**Supplementary table1. LAI value for the whole genome assembly of little millet**

| Chr | From | To | Intact | Total | raw_LAI | LAI |
| --- | --- | --- | --- | --- | --- | --- |
| whole_genome | 1 | 566591115 | 0.0752 | 0.4304 | 17.47 | 18.26 |

**Supplementary table 2. Statistics of species-specific orthologous groups**

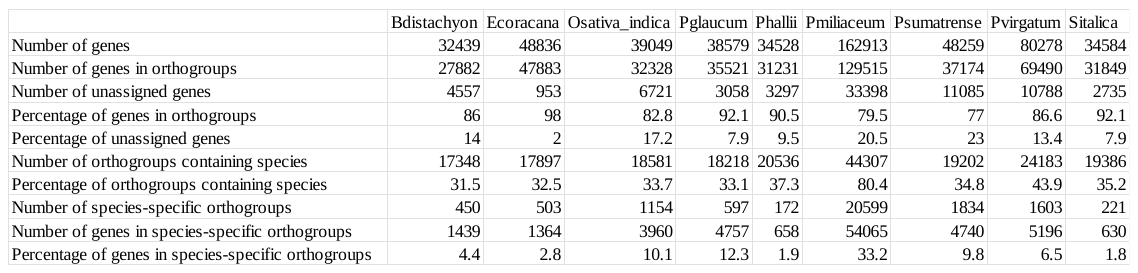

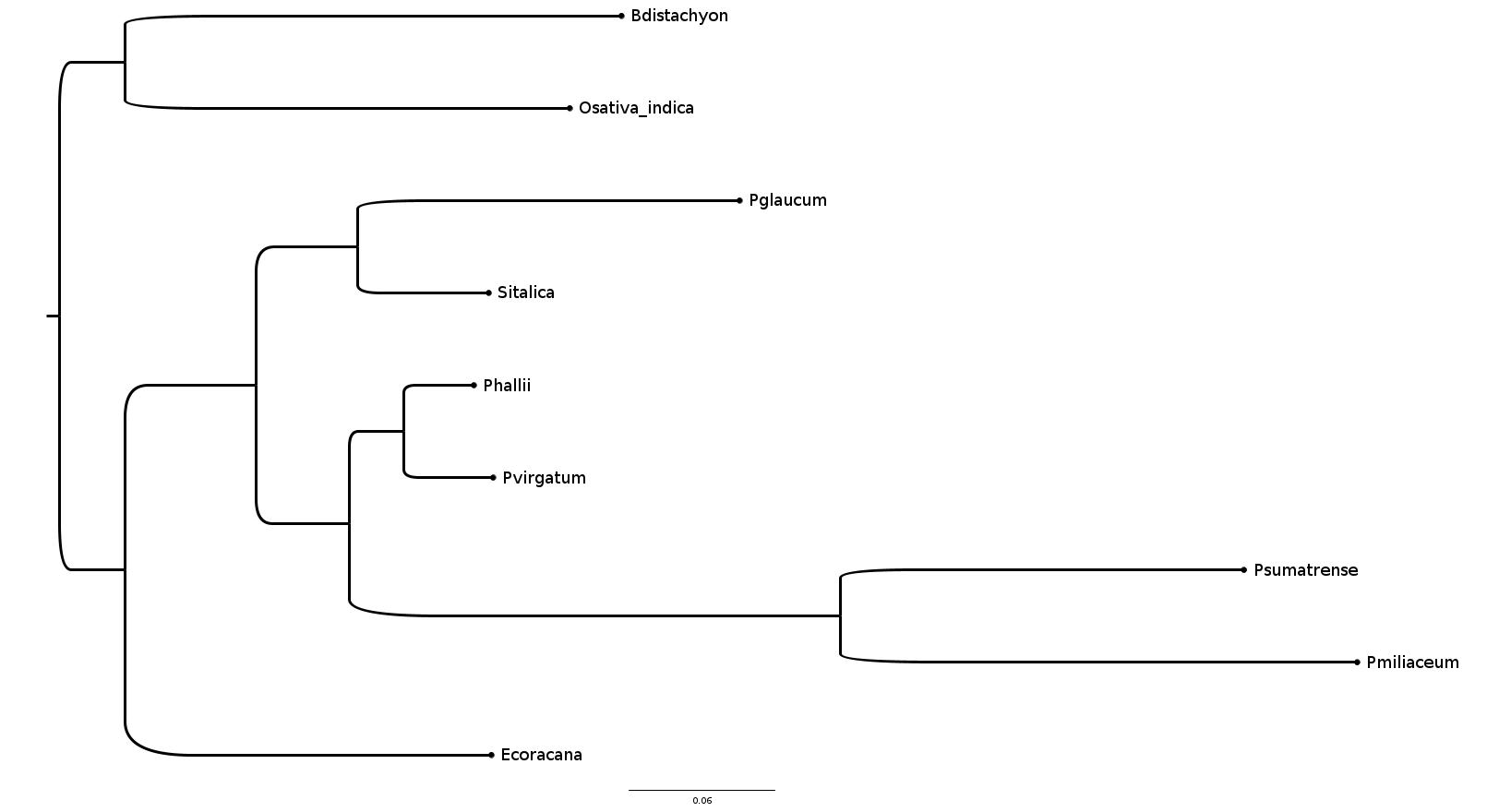

**Supplementary figure S1. Phylogenetic relationship between different millets and rice based on the inferences from single copy orthologs.**

**Supplementary table 3. *P. sumatrense* specific genes grouped into GO terms.**

| **PANTHER GO-Slim Molecular Function** | **Number of genes of *P.sumatrense* represented** | **raw P-value** | **FDR** |
| --- | --- | --- | --- |
| hydrolase activity, acting on carbon-nitrogen (but not peptide) bonds (GO:0016810) | 4 | 1.50E-05 | 1.20E-03 |
| glucosidase activity (GO:0015926) | 3 | 2.34E-04 | 1.12E-02 |
| hydrolase activity, hydrolyzing O-glycosyl compounds (GO:0004553) | 5 | 3.32E-05 | 2.28E-03 |
| hydrolase activity, acting on glycosyl bonds (GO:0016798) | 5 | 6.34E-05 | 3.80E-03 |
| ATP-dependent activity (GO:0140657) | 6 | 6.90E-05 | 3.67E-03 |
| oxidoreductase activity (GO:0016491) | 12 | 9.43E-07 | 9.04E-05 |
| hydrolase activity (GO:0016787) | 17 | 4.13E-09 | 4.95E-07 |
| catalytic activity (GO:0003824) | 44 | 6.62E-18 | 1.06E-15 |
| catalytic activity, acting on a protein (GO:0140096) | 11 | 3.48E-04 | 1.51E-02 |
| molecular_function (GO:0003674) | 56 | 4.80E-18 | 1.15E-15 |
| Unclassified (UNCLASSIFIED) | 40 | 4.80E-18 | 2.30E-15 |
| **PANTHER GO-Slim Biological Process** |  |  |  |
| phosphate-containing compound metabolic process (GO:0006796) | 9 | 1.64E-04 | 3.27E-02 |
| phosphorus metabolic process (GO:0006793) | 9 | 1.76E-04 | 3.08E-02 |
| biological regulation (GO:0065007) | 15 | 8.76E-06 | 3.06E-03 |
| regulation of cellular process (GO:0050794) | 11 | 2.33E-04 | 3.25E-02 |
| regulation of biological process (GO:0050789) | 12 | 2.21E-04 | 3.44E-02 |
| organonitrogen compound metabolic process (GO:1901564) | 14 | 3.17E-04 | 4.02E-02 |
| cellular metabolic process (GO:0044237) | 17 | 1.55E-04 | 4.34E-02 |
| cellular process (GO:0009987) | 28 | 1.05E-06 | 4.89E-04 |
| biological_process (GO:0008150) | 42 | 1.53E-09 | 2.14E-06 |
| metabolic process (GO:0008152) | 22 | 1.56E-04 | 3.62E-02 |
| organic substance metabolic process (GO:0071704) | 20 | 4.17E-04 | 4.85E-02 |
| Unclassified (UNCLASSIFIED) | 54 | 1.53E-09 | 1.07E-06 |

**Supplementary table 4. *In silico* expression matrix for *P.sumatrense* leaf tissue in response to drought and salinity stress**

| Gene ID | Control_Leaf1 | Control_Leaf2 | Drought_Leaf1 | Drought_Leaf2 | Salt_Leaf1 | Salt_Leaf2 |
| --- | --- | --- | --- | --- | --- | --- |
| g10222.t1 | 1.839 | 3.092 | 101.421 | 32.78 | 46.383 | 24.594 |
| g8153.t1 | 0.254 | 0.307 | 1.509 | 3.352 | 3.523 | 7.456 |
| g42231.t1 | 37.911 | 15.819 | 0 | 4.906 | 0 | 0 |
| g31054.t1 | 10.537 | 29.443 | 0 | 0 | 0 | 0 |
| g8224.t1 | 6.349 | 4.279 | 277.689 | 68.018 | 28.513 | 23.746 |
| g16545.t1 | 6.79 | 8.55 | 606.125 | 71.482 | 67.267 | 51.584 |
| g43402.t1 | 8.365 | 10.933 | 288.842 | 93.433 | 52.107 | 46.303 |
| g12304.t1 | 0.323 | 0 | 18.597 | 30.301 | 13.747 | 7.526 |
| g25494.t1 | 2.798 | 0 | 481.638 | 129.809 | 0 | 33.847 |
| g16058.t1 | 5.606 | 5.116 | 521.835 | 205.171 | 29.474 | 60.916 |
| g37968.t1 | 0 | 0.248 | 10.606 | 7.202 | 15.595 | 6.019 |
| g33874.t1 | 0.078 | 0 | 1.452 | 1.707 | 1.807 | 1.777 |
| g5830.t1 | 3.874 | 8.482 | 1163.817 | 345.971 | 516.828 | 292.913 |
| g26739.t1 | 0 | 0.12 | 44.265 | 0.802 | 4.853 | 9.073 |
| g25529.t1 | 2.935 | 4.766 | 219.297 | 100.026 | 28.99 | 14.982 |
| g7528.t1 | 0 | 0 | 773.037 | 150.938 | 10.43 | 7.496 |
| g45520.t1 | 0.205 | 0 | 38.171 | 11.641 | 1.725 | 3.234 |
| g40833.t1 | 0 | 0.265 | 30.554 | 38.519 | 4.484 | 0.23 |
| g29822.t1 | 0.215 | 0 | 81.847 | 13.012 | 9.764 | 1.427 |
| g18457.t1 | 0 | 0.111 | 4.254 | 6.897 | 1.486 | 0.389 |
| g38283.t1 | 6.036 | 5.936 | 0 | 1.91 | 0.567 | 0 |
| g23165.t1 | 0 | 0 | 0 | 23.983 | 7.342 | 28.247 |
| g25332.t1 | 0 | 0 | 2.199 | 2.59 | 4.114 | 2.555 |
| g38962.t1 | 1.311 | 0.974 | 66.354 | 15.521 | 7.383 | 0.579 |
| g15051.t1 | 0 | 0.171 | 0 | 1.737 | 3.909 | 8.285 |
| g18556.t1 | 0 | 0.572 | 194.204 | 173.427 | 29.868 | 6.188 |
| g21966.t1 | 1.849 | 1.999 | 69.2 | 36.792 | 5.453 | 0.968 |
| g15905.t1 | 1.047 | 0.495 | 124.43 | 23.841 | 22.789 | 4.412 |
| g22686.t1 | 3.679 | 3.348 | 76.385 | 52.568 | 31.659 | 33.887 |
| g7530.t1 | 1.536 | 0 | 740.284 | 141.999 | 8.722 | 7.925 |
| g42052.t1 | 2.074 | 1.623 | 71.485 | 20.722 | 9.001 | 5.2 |
| g39238.t1 | 3.082 | 4.288 | 138.485 | 176.455 | 121.978 | 105.163 |
| g4338.t2 | 0.675 | 3.468 | 38.186 | 5.587 | 71.858 | 50.406 |
| g20957.t1 | 12.337 | 10.045 | 2.127 | 0.894 | 0 | 0.289 |
| g8409.t1 | 183.81 | 163.913 | 8.479 | 7.344 | 19.159 | 29.924 |
| g25873.t1 | 0 | 0 | 10.477 | 34.08 | 0.895 | 0 |
| g12358.t1 | 134.453 | 80.718 | 5.088 | 0.945 | 2.414 | 0.928 |
| g13994.t1 | 0 | 0 | 111.998 | 32.434 | 6.857 | 0 |
| g25221.t1 | 3.033 | 3.681 | 482.974 | 193.215 | 244.72 | 453.692 |
| g35868.t1 | 6.183 | 3.126 | 169.974 | 139.591 | 211.476 | 175.203 |
| g31298.t1 | 0.196 | 0.478 | 4.714 | 5.567 | 6.044 | 5.5 |
| g37003.t1 | 5.009 | 5.825 | 1442.542 | 71.38 | 55.195 | 17.218 |
| g43770.t1 | 3.884 | 3.758 | 143.084 | 46.93 | 56.255 | 42.321 |
| g19489.t1 | 180.161 | 254.924 | 6419.636 | 2372.177 | 772.552 | 556.271 |
| g15029.t1 | 27.129 | 53.599 | 886.114 | 484.313 | 125.813 | 86.538 |
| g3070.t1 | 0.235 | 0 | 53.549 | 1.31 | 3.975 | 7.995 |
| g20284.t1 | 0.362 | 1.785 | 376.954 | 390.087 | 179.809 | 13.165 |
| g6665.t1 | 0.489 | 0 | 52.902 | 23.394 | 9.666 | 11.818 |
| g8282.t2 | 0.421 | 1.17 | 0 | 11.306 | 35.412 | 8.384 |
| g3401.t1 | 0.372 | 0.367 | 30.482 | 3.657 | 7.941 | 18.895 |
| g46326.t1 | 0 | 0 | 41.822 | 16.608 | 0 | 0.549 |
| g9248.t1 | 4.559 | 6.218 | 408.859 | 158.759 | 47.467 | 83.574 |
| g40078.t1 | 182.362 | 433.042 | 19867.893 | 5140.82 | 2876.095 | 9312.599 |
| g16147.t1 | 1.008 | 0.965 | 60.562 | 33.308 | 25.77 | 2.346 |
| g36590.t1 | 67.427 | 50.788 | 2404.006 | 1161.11 | 301.566 | 251.41 |
| g28048.t1 | 50.384 | 57.664 | 0 | 1.29 | 0 | 0.769 |
| g34201.t1 | 0 | 0.273 | 253.487 | 20.499 | 2.538 | 0 |
| g3392.t1 | 1.262 | 2.426 | 395.12 | 107.187 | 27.388 | 9.203 |
| g37513.t1 | 135.559 | 307.31 | 17.648 | 61.07 | 4.41 | 5.5 |
| g7748.t1 | 22.022 | 30.921 | 1094.23 | 973.807 | 253.532 | 477.747 |
| g63.t1 | 0.519 | 0.624 | 203.201 | 39.901 | 23.159 | 33.258 |
| g45240.t1 | 34.164 | 37.13 | 26.487 | 11.895 | 1.905 | 2.056 |
| g40141.t1 | 0.871 | 0.88 | 37.381 | 16.659 | 8.442 | 22.119 |
| g12737.t1 | 80.723 | 276.62 | 34881.67 | 7891.727 | 2871.512 | 1021.701 |
| g20132.t1 | 20.868 | 15.836 | 3903.186 | 1461.685 | 661.619 | 835.908 |
| g17312.t1 | 5.185 | 6.714 | 3.305 | 0.894 | 0 | 0.11 |
| g3394.t1 | 1.888 | 1.734 | 59.326 | 30.037 | 13.23 | 27.768 |
| g41924.t1 | 0 | 1.008 | 103.16 | 23.861 | 18.404 | 7.915 |
| g44602.t1 | 0 | 0 | 0 | 19.097 | 5.584 | 7.646 |
| g21536.t1 | 0 | 1.17 | 14.3 | 5.831 | 10.898 | 238.395 |
| g5286.t1 | 6.075 | 3.101 | 140.857 | 107.147 | 29.343 | 73.403 |
| g26678.t1 | 0.166 | 1.845 | 1.006 | 1.676 | 48.822 | 40.564 |
| g31054.t2 | 0 | 0.034 | 9.198 | 2.346 | 10.734 | 26.77 |
| g9483.t1 | 2.162 | 1.059 | 116.324 | 164.153 | 86.205 | 65.797 |
| g25391.t1 | 0 | 0 | 16.226 | 28.574 | 12.869 | 14.842 |
| g37905.t1 | 0 | 0.735 | 14.357 | 152.035 | 70.585 | 18.086 |
| g37247.t2 | 0 | 0.017 | 0 | 0 | 43.238 | 2.685 |
| g25559.t1 | 0 | 0 | 34.075 | 63.508 | 3.383 | 2.296 |
| g8481.t1 | 1.223 | 0 | 336.556 | 21.962 | 10.807 | 0.579 |
| g32414.t1 | 0.147 | 0 | 2.688 | 7.293 | 26.789 | 10.979 |
| g21406.t1 | 0.323 | 0 | 91.346 | 10.767 | 5.396 | 4.182 |
| g43321.t1 | 6.017 | 23.583 | 19.186 | 0.447 | 0 | 0.439 |
| g26639.t1 | 0.548 | 3.348 | 91.519 | 60.745 | 18.387 | 19.833 |
| g4073.t1 | 34.956 | 27.231 | 8.494 | 20.479 | 0 | 0 |
| g38145.t1 | 0.685 | 1.256 | 28.6 | 14.384 | 32.849 | 10.84 |
| g7995.t1 | 0.382 | 0.47 | 18.482 | 15.846 | 16.252 | 14.383 |
| g8601.t1 | 0.235 | 0 | 23.958 | 16.893 | 0 | 0.26 |
| g6621.t1 | 3.796 | 8.806 | 3666.169 | 1026.445 | 162.662 | 52.452 |
| g17074.t1 | 1.624 | 1.11 | 72.376 | 46.117 | 22.28 | 14.952 |
| g10072.t1 | 0.675 | 2.204 | 9.413 | 20.712 | 40.224 | 13.724 |
| g45442.t1 | 1.771 | 4.647 | 80.855 | 101.905 | 47.558 | 19.224 |
| g8874.t1 | 0 | 0 | 25.467 | 7.039 | 0 | 0 |
| g32620.t1 | 0 | 0 | 118.049 | 2.184 | 7.095 | 7.566 |
| g2449.t1 | 0.078 | 0.094 | 0.891 | 3.403 | 2.505 | 6.478 |
| g38465.t1 | 47.508 | 55.495 | 3.722 | 2.59 | 0.871 | 12.796 |
| g45603.t1 | 48.711 | 55.819 | 0 | 12.352 | 0 | 0.938 |
| g15905.t2 | 0 | 0.393 | 111.237 | 44.126 | 8.722 | 1.377 |
| g27121.t1 | 10.889 | 31.467 | 0 | 0 | 0 | 0.329 |
| g25635.t1 | 119.485 | 217.469 | 0 | 0 | 0 | 0 |
| g10726.t1 | 31.121 | 42.597 | 5693.967 | 842.951 | 265.127 | 218.212 |
| g36617.t1 | 16.573 | 11.155 | 0 | 2.326 | 0 | 0 |
| g30297.t1 | 0.147 | 0 | 52.37 | 31.297 | 13.444 | 9.213 |
| g37577.t1 | 0.166 | 1.435 | 77.161 | 24.156 | 0.936 | 0 |
| g4537.t1 | 1.575 | 0.641 | 70.579 | 39.84 | 8.861 | 2.006 |
| g42892.t1 | 0 | 0 | 90.038 | 30.423 | 0 | 41.163 |
| g33708.t1 | 95.916 | 42.409 | 21.859 | 3.576 | 1.207 | 5.889 |
| g6959.t1 | 2.964 | 4.544 | 120.866 | 44.411 | 67.596 | 37.979 |
| g35421.t2 | 0 | 0 | 12.949 | 7.679 | 6.274 | 0.03 |
| g15050.t1 | 0 | 0.171 | 0.819 | 3.362 | 5.396 | 2.715 |
| g14135.t1 | 41.305 | 11.693 | 1224.078 | 1307.771 | 871.938 | 422.041 |
| g21640.t1 | 0.567 | 0.461 | 102.068 | 11.783 | 1.068 | 11.419 |
| g19212.t1 | 6.066 | 29.101 | 1140.492 | 634.367 | 364.948 | 453.143 |
| g26328.t2 | 107.353 | 143.174 | 3856.449 | 1347.489 | 266.425 | 136.186 |
| g13689.t1 | 1.849 | 1.614 | 97.857 | 33.064 | 1.486 | 2.026 |
| g22695.t2 | 66.869 | 92.147 | 48.174 | 62.055 | 0 | 0 |
| g7716.t1 | 0.196 | 0 | 78.11 | 8.431 | 0 | 1.068 |
| g12159.t1 | 5.616 | 10.549 | 1265.885 | 96.267 | 142.147 | 147.804 |
| g26865.t1 | 2.661 | 1.614 | 283.021 | 68.475 | 2.488 | 0.958 |
| g31542.t1 | 54.513 | 157.575 | 3077.261 | 2295.819 | 603.763 | 489.386 |
| g902.t1 | 1.282 | 0 | 364.882 | 44.604 | 0 | 0 |
| g23982.t1 | 0 | 1.811 | 251.82 | 59.739 | 4.27 | 8.065 |
| g40900.t1 | 19.743 | 37.677 | 8647.963 | 3955.006 | 2379.683 | 316.658 |
| g22737.t1 | 13.178 | 10.549 | 124.229 | 233.482 | 135.454 | 268.878 |
| g11343.t1 | 0 | 0 | 8.005 | 6.308 | 4.361 | 0.858 |
| g42096.t1 | 39.006 | 30.058 | 844.594 | 339.562 | 149.276 | 105.413 |
| g4919.t1 | 2.289 | 3.306 | 114.341 | 39.626 | 34.648 | 20.841 |
| g31522.t1 | 0 | 0.897 | 62.646 | 106.588 | 40.199 | 8.454 |
| g14378.t1 | 0 | 0 | 55.101 | 67.805 | 0 | 2.156 |
| g25851.t1 | 5.283 | 22.815 | 1303.079 | 320.332 | 123.21 | 80.869 |
| g43000.t1 | 38.057 | 101.338 | 1789.862 | 1286.226 | 627.325 | 880.355 |
| g2810.t1 | 0 | 0 | 4.441 | 2.702 | 7.104 | 5.809 |
| g46479.t1 | 5.449 | 6.517 | 153.188 | 69.41 | 24.966 | 46.353 |
| g21488.t1 | 2.769 | 0 | 582.771 | 124.771 | 15.636 | 47.771 |
| g5459.t1 | 0.157 | 0 | 0 | 2.103 | 4.426 | 3.124 |
| g27892.t1 | 1.36 | 3.289 | 1624.659 | 1447.383 | 183.341 | 60.687 |
| g43958.t1 | 13.922 | 8.362 | 0 | 9.112 | 0 | 0.339 |
| g11188.t1 | 18.96 | 6.346 | 2802.159 | 3181.495 | 322.359 | 681.477 |
| g43036.t1 | 0 | 0.265 | 0 | 5.81 | 13.501 | 27.738 |
| g39590.t1 | 0.597 | 0 | 170.405 | 49.256 | 13.411 | 23.656 |
| g21195.t2 | 8.189 | 5.185 | 0 | 4.663 | 0 | 0 |
| g42941.t1 | 2.593 | 2.246 | 100.559 | 37.29 | 15.406 | 15.741 |
| g26328.t1 | 169.977 | 160.138 | 3680.785 | 1739.201 | 736.795 | 189.027 |
| g44745.t1 | 0 | 0 | 42.382 | 12.088 | 3.071 | 0 |
| g26993.t1 | 0.538 | 0.905 | 10.017 | 5.567 | 18.371 | 27.908 |
| g41213.t1 | 0 | 0 | 517.265 | 125.482 | 0 | 8.354 |
| g16101.t1 | 3.698 | 3.673 | 0.848 | 2.062 | 0 | 0 |
| g9521.t1 | 9.891 | 9.643 | 0.949 | 2.966 | 0.879 | 0.349 |
| g38689.t1 | 0.969 | 1.734 | 35.11 | 19.767 | 24.284 | 15.84 |
| g5458.t1 | 0.254 | 1.862 | 19.905 | 27.223 | 46.909 | 45.295 |
| g18900.t1 | 18.491 | 20.081 | 0 | 8.411 | 0.756 | 0.299 |
| g5079.t1 | 0.245 | 0.154 | 37.338 | 8.827 | 0 | 0.808 |
| g46625.t1 | 0.264 | 0.325 | 47.053 | 50.871 | 11.818 | 4.591 |
| g27718.t1 | 0 | 0 | 35.254 | 23.516 | 6.685 | 17.617 |
| g2914.t1 | 0 | 0.29 | 93.085 | 30.454 | 8.902 | 0.789 |
| g7389.t1 | 0 | 0.384 | 75.983 | 58.551 | 106.062 | 124.897 |
| g32573.t1 | 5.586 | 3.323 | 3.923 | 1.138 | 0 | 0 |
| g46980.t1 | 0 | 0 | 12.13 | 5.465 | 0.542 | 0.21 |
| g46631.t1 | 2.094 | 3.34 | 5.921 | 0.477 | 0 | 0.07 |
| g21847.t1 | 19.635 | 31.356 | 0.762 | 0.427 | 0 | 1.407 |
| g25635.t2 | 125.325 | 234.262 | 2.759 | 11.783 | 0 | 2.525 |
| g38297.t1 | 99.292 | 19.594 | 0 | 3.616 | 4.246 | 0.828 |
| g44242.t1 | 172.217 | 149.427 | 5530.432 | 2343.612 | 735.177 | 1223.424 |
| g19284.t1 | 4.735 | 3.528 | 135.122 | 47.174 | 63.613 | 46.503 |
| g18278.t1 | 0 | 0 | 21.486 | 4.195 | 0.542 | 0 |
| g44436.t1 | 1.057 | 2.195 | 49.942 | 23.597 | 53.183 | 59.489 |
| g196.t1 | 3.737 | 2.494 | 0 | 1.188 | 0 | 0 |
| g5396.t1 | 0.734 | 0.299 | 140.842 | 20.936 | 2.759 | 9.133 |
| g4649.t1 | 0.489 | 3.767 | 326.179 | 61.852 | 62.627 | 37.37 |
| g29678.t1 | 2.397 | 2.477 | 408.701 | 118.981 | 60.689 | 46.693 |
| g18742.t2 | 3.229 | 3.374 | 0 | 3.891 | 0 | 0 |
| g19264.t1 | 3.816 | 9.046 | 157.657 | 81.264 | 55.598 | 45.046 |
| g28434.t1 | 0 | 0.376 | 4.268 | 1.351 | 5.116 | 5.13 |
| g16987.t1 | 0.078 | 0.196 | 95.859 | 7.761 | 0.895 | 7.396 |
| g4791.t1 | 10.175 | 12.078 | 412.769 | 193.642 | 69.887 | 102.069 |
| g27078.t1 | 0 | 0 | 5.174 | 5.516 | 2.414 | 0.12 |
| g4819.t1 | 0.411 | 1.999 | 97.67 | 7.69 | 16.228 | 33.487 |
| g32155.t2 | 16.906 | 22.336 | 0 | 0 | 23.496 | 6.009 |
| g7576.t1 | 102.657 | 82.35 | 2759.374 | 853.678 | 200.973 | 149.241 |
| g2910.t1 | 45.375 | 20.141 | 0 | 0.792 | 0 | 0 |
| g23914.t1 | 1.565 | 0.769 | 101.32 | 17.919 | 4.394 | 0.519 |
| g2176.t1 | 21.122 | 32.407 | 1309.618 | 314.42 | 90.007 | 40.514 |
| g8860.t2 | 0 | 0 | 357.251 | 37.798 | 17.533 | 9.462 |
| g476.t1 | 1.37 | 4.894 | 76.644 | 37.666 | 51.836 | 84.422 |
| g29068.t1 | 1.428 | 2.144 | 90.484 | 42.389 | 47.278 | 47.631 |
| g7695.t1 | 1.213 | 0.726 | 166.726 | 270.772 | 27.216 | 3.244 |
| g10971.t1 | 0.753 | 0 | 37.812 | 11.956 | 12.647 | 14.243 |
| g14990.t1 | 0.245 | 0.222 | 11.972 | 5.445 | 1.692 | 1.068 |
| g17169.t1 | 4.089 | 5.253 | 724.001 | 260.461 | 75.48 | 61.276 |
| g25321.t1 | 5.078 | 8.149 | 198.803 | 163.3 | 31.708 | 35.114 |
| g31923.t1 | 2.24 | 3.895 | 163.65 | 44.797 | 37.744 | 23.326 |
| g42762.t1 | 1.321 | 4.254 | 4587.665 | 3634.917 | 838.218 | 1267.561 |
| g15657.t2 | 0 | 0 | 1.567 | 0 | 16.244 | 8.614 |
| g10822.t1 | 5.802 | 9.652 | 23.512 | 1.087 | 0 | 0 |
| g20382.t1 | 6.878 | 19.577 | 1249.717 | 531.375 | 59.753 | 130.546 |
| g38551.t1 | 2.426 | 2.537 | 87.006 | 53.959 | 9.699 | 4.831 |
| g30393.t1 | 0 | 0.171 | 137.925 | 7.456 | 6.143 | 5.26 |
| g22919.t1 | 0.939 | 2.861 | 15.392 | 33.196 | 39.698 | 56.854 |
| g1897.t1 | 0.655 | 0.401 | 32.997 | 9.315 | 5.51 | 2.435 |
| g6281.t1 | 0.127 | 0.162 | 52.298 | 26.736 | 8.894 | 2.905 |
| g4716.t1 | 0.655 | 0 | 30.525 | 33.938 | 11.005 | 72.385 |
| g39632.t1 | 3.659 | 2.118 | 20.609 | 19.107 | 114.768 | 46.842 |
| g43661.t1 | 0 | 0 | 21.04 | 4.459 | 10.331 | 9.732 |
| g36301.t1 | 30.759 | 79.181 | 3406.271 | 669.514 | 304.21 | 185.713 |
| g43946.t1 | 0.704 | 3.015 | 283.97 | 53.177 | 41.932 | 31.86 |
| g13420.t1 | 12.317 | 36.848 | 2995.802 | 677.691 | 371.272 | 169.533 |
| g4028.t1 | 0 | 0 | 190.827 | 60.806 | 14.675 | 5.58 |
| g30554.t1 | 98.48 | 44.177 | 0 | 0.589 | 1.979 | 0.19 |
| g33387.t1 | 4.813 | 26.069 | 164.786 | 40.185 | 143.905 | 474.643 |
| g31543.t1 | 23.314 | 38.232 | 981.326 | 541.878 | 73.788 | 87.127 |
| g4147.t2 | 9.959 | 19.586 | 4980.069 | 715.672 | 169.207 | 78.923 |
| g14597.t1 | 0.313 | 0.196 | 1.911 | 15.186 | 12.491 | 8.544 |
| g9961.t1 | 20.496 | 16.912 | 0 | 0 | 0 | 7.576 |
| g9736.t1 | 0 | 0 | 440.635 | 193.286 | 568.549 | 3.274 |
| g8332.t1 | 0 | 0 | 82.968 | 18.731 | 0 | 0 |
| g12433.t1 | 0 | 0 | 8.853 | 30.007 | 10.545 | 1.916 |
| g38248.t1 | 103.283 | 135.675 | 3555.507 | 1772.936 | 348.852 | 238.514 |
| g35053.t1 | 0.861 | 1.042 | 40.657 | 8.949 | 16.901 | 20.442 |
| g41037.t1 | 0 | 0 | 244.864 | 42.095 | 0 | 8.145 |
| g30614.t1 | 20.848 | 26.778 | 1878.635 | 322.933 | 135.512 | 254.325 |
| g28468.t1 | 0.382 | 0.931 | 144.953 | 0.416 | 21.467 | 19.883 |
| g13374.t1 | 0.851 | 2.998 | 79.921 | 47.55 | 12.22 | 31.741 |
| g26070.t1 | 0.108 | 0.794 | 36.964 | 15.511 | 4.238 | 3.204 |
| g45432.t1 | 0.117 | 0.299 | 3.607 | 12.708 | 12.138 | 11.239 |
| g6538.t1 | 0 | 1.196 | 818.754 | 80.35 | 27.479 | 10.301 |
| g8188.t1 | 0.098 | 0 | 47.901 | 3.555 | 0 | 0.1 |
| g7729.t1 | 0 | 0 | 30.396 | 45.843 | 20.974 | 2.376 |
| g46137.t1 | 7.885 | 15.614 | 569.535 | 289.025 | 102.925 | 132.942 |
| g31099.t1 | 0 | 2.135 | 295.913 | 106.933 | 26.132 | 51.424 |
| g20484.t1 | 103.508 | 181.962 | 4617.86 | 2385.626 | 766.163 | 486.83 |
| g44142.t1 | 0 | 0 | 46.722 | 3.758 | 1.757 | 0.17 |
| g22220.t1 | 121.128 | 155.397 | 8821.069 | 2123.235 | 1082.839 | 239.662 |
| g16750.t1 | 0.176 | 0 | 54.425 | 12.758 | 9.625 | 4.701 |
| g4590.t1 | 0 | 0.419 | 209.841 | 29.123 | 8.327 | 7.286 |
| g15977.t1 | 0 | 0.307 | 119.055 | 49.561 | 40.766 | 16.29 |
| g19709.t1 | 1.761 | 2.118 | 269.095 | 60.725 | 29.753 | 35.294 |
| g15672.t1 | 440.732 | 205.596 | 23.067 | 46.067 | 12.031 | 9.832 |
| g36911.t1 | 50.414 | 51.113 | 9107.295 | 4183.916 | 2163.961 | 342.021 |
| g25927.t1 | 0.196 | 0.495 | 280.391 | 190.757 | 14.634 | 7.067 |
| g8416.t1 | 3.855 | 3.724 | 0 | 0 | 0 | 0 |
| g11272.t1 | 1.076 | 2.204 | 468.919 | 80.959 | 19.159 | 6.697 |
| g46038.t1 | 0 | 2.656 | 93.358 | 28.839 | 17.254 | 25.133 |
| g19023.t1 | 1.448 | 1.025 | 25.179 | 43.791 | 15.735 | 30.104 |
| g15481.t1 | 1.399 | 1.332 | 65.132 | 25.385 | 3.49 | 5.12 |
| g7002.t1 | 140.558 | 223.294 | 3.909 | 9.518 | 12.302 | 110.404 |
| g1993.t1 | 0.333 | 0.273 | 185.38 | 27.792 | 2.497 | 1.347 |
| g18194.t1 | 3.365 | 2.34 | 0 | 0.853 | 0 | 0.18 |
| g34052.t1 | 129.248 | 64.472 | 2.817 | 6.156 | 5.248 | 8.574 |
| g8012.t1 | 19.997 | 15.81 | 3.363 | 4.144 | 0 | 0.818 |
| g5223.t1 | 0 | 0 | 329.93 | 91.717 | 26.633 | 52.143 |
| g42915.t1 | 0.587 | 0.359 | 33.357 | 23.76 | 9.937 | 3.533 |
| g16243.t1 | 2.642 | 2.11 | 122.734 | 97.293 | 141.515 | 148.512 |
| g1667.t1 | 110.357 | 87.022 | 4319.706 | 1694.933 | 502.99 | 463.614 |
| g22603.t1 | 0 | 0.333 | 1.638 | 3.84 | 4.574 | 9.892 |
| g4835.t1 | 0 | 0 | 44.969 | 32.485 | 0 | 3.284 |
| g23576.t1 | 0 | 1.896 | 133.987 | 93.159 | 347.859 | 324.693 |
| g31332.t1 | 5.087 | 8.328 | 21.529 | 0 | 0 | 0 |
| g20350.t1 | 0 | 0.401 | 16.01 | 24.938 | 21.204 | 3.923 |
| g41056.t1 | 0 | 0.598 | 237.65 | 17.848 | 8.919 | 12.816 |
| g7005.t1 | 0 | 1.631 | 164.268 | 176.688 | 140.193 | 128.26 |
| g22841.t1 | 0.616 | 0 | 6.151 | 4.104 | 5.765 | 8.774 |
| g26550.t1 | 1.536 | 7.209 | 212.672 | 221.241 | 47.533 | 15.96 |
| g46713.t1 | 0.43 | 0.102 | 0.517 | 3.616 | 8.623 | 5.46 |
| g12235.t1 | 0.528 | 4.459 | 1246.368 | 3465.299 | 1410.191 | 917.925 |
| g6953.t1 | 0.196 | 0 | 17.404 | 11.184 | 0 | 0.429 |
| g15275.t1 | 339.699 | 183.397 | 1511.482 | 17.99 | 2.209 | 2.715 |
| g11981.t2 | 0 | 0 | 0 | 0 | 16.458 | 13.335 |
| g1956.t1 | 0 | 0 | 4.685 | 4.012 | 7.063 | 3.673 |
| g22080.t1 | 4.344 | 10.412 | 717.721 | 137.011 | 84.333 | 43.958 |
| g25742.t1 | 7.915 | 3.827 | 273.522 | 95.607 | 2.973 | 3.404 |
| g14781.t1 | 0.108 | 0 | 6.625 | 13.632 | 3.712 | 0 |
| g25628.t1 | 0.616 | 0.752 | 105.689 | 16.862 | 12.688 | 7.426 |
| g39339.t1 | 0 | 1.273 | 41.822 | 62.309 | 11.67 | 12.197 |
| g31819.t1 | 7.915 | 25.958 | 1006.922 | 404.928 | 267.575 | 208.64 |
| g9820.t1 | 10.957 | 4.638 | 184.23 | 98.868 | 39.969 | 39.386 |
| g25298.t1 | 223.394 | 164.725 | 18.065 | 35.502 | 0 | 13.884 |
| g15774.t1 | 1.918 | 1.563 | 8.609 | 19.503 | 57.306 | 22.588 |
| g20000.t1 | 30.192 | 24.856 | 0 | 6.582 | 0 | 1.847 |
| g25322.t1 | 24.008 | 44.459 | 1186.999 | 721.969 | 186.445 | 124.537 |
| g45242.t1 | 5.586 | 11.01 | 6.079 | 3.007 | 0 | 0 |
| g43386.t1 | 0.157 | 0.188 | 79.145 | 35.909 | 6.102 | 11.429 |
| g1266.t1 | 42.352 | 71.032 | 3928.911 | 1253.233 | 803.759 | 615.2 |
| g24591.t1 | 0.254 | 0.102 | 4.656 | 9.01 | 7.235 | 2.745 |
| g8360.t1 | 1.017 | 3.289 | 380.073 | 128.925 | 116.714 | 77.385 |
| g21419.t1 | 1.585 | 0.649 | 154.682 | 36.863 | 11.891 | 17.917 |
| g27691.t1 | 0 | 0.427 | 0 | 5.77 | 5.79 | 4.581 |
| g20560.t1 | 35.563 | 9.114 | 49.237 | 98.289 | 0 | 0.868 |
| g25885.t1 | 19.019 | 29.144 | 1093.698 | 389.488 | 232.828 | 213.182 |
| g26404.t1 | 8.815 | 6.218 | 1.293 | 0.65 | 0 | 0.09 |
| g26992.t1 | 7.044 | 5.663 | 136.746 | 35.675 | 136.998 | 159.133 |
| g7028.t1 | 2.984 | 3.562 | 269.21 | 101.336 | 50.342 | 45.964 |
| g14553.t1 | 0 | 0 | 39.163 | 8.573 | 3.244 | 3.893 |
| g11596.t1 | 2.182 | 1.461 | 122.36 | 29.732 | 16.72 | 9.133 |
| g19194.t1 | 17.131 | 15.289 | 844.206 | 176.069 | 122.29 | 138.272 |
| g7643.t1 | 26.63 | 28.435 | 1647.338 | 319.479 | 48.863 | 37.72 |
| g3794.t1 | 2.152 | 51.121 | 376.451 | 2.509 | 0 | 0.21 |
| g18059.t1 | 3.757 | 4.937 | 252.136 | 63.508 | 38.064 | 63.441 |
| g18610.t1 | 104.555 | 91.139 | 10.764 | 1.991 | 3.383 | 1.637 |
| g19335.t1 | 0 | 0 | 10.118 | 7.253 | 17.648 | 17.677 |
| g12611.t1 | 0 | 0 | 531.694 | 98.512 | 20.12 | 12.137 |
| g5253.t1 | 1.379 | 1.529 | 517.409 | 152.055 | 13.961 | 6.079 |
| g23574.t1 | 1.604 | 0.649 | 28.327 | 9.914 | 18.043 | 23.007 |
| g36008.t1 | 14.607 | 19.167 | 5388.023 | 2437.391 | 587.24 | 857.418 |
| g15672.t2 | 535.132 | 448.579 | 30.985 | 0 | 0 | 11.738 |
| g44412.t1 | 15.556 | 27.948 | 334.457 | 495.639 | 246.847 | 159.801 |
| g14091.t1 | 140.216 | 92.941 | 8956.163 | 1938.085 | 829.816 | 434.059 |
| g33801.t1 | 102.804 | 53.206 | 15.435 | 10.016 | 4.829 | 4.003 |
| g24192.t1 | 0 | 0 | 70.349 | 45.457 | 1.897 | 2.206 |
| g6690.t1 | 0.411 | 2.477 | 469.091 | 85.591 | 32.127 | 16.809 |
| g32596.t1 | 0.157 | 0 | 7.286 | 6.928 | 17.903 | 12.676 |
| g44263.t1 | 5.674 | 3.22 | 253.631 | 177.613 | 19.143 | 6.977 |
| g44701.t1 | 0 | 1.461 | 383.522 | 61.822 | 7.03 | 21.989 |
| g20323.t1 | 90.643 | 35.584 | 1.725 | 4.784 | 1.618 | 0.789 |
| g33611.t1 | 8.756 | 8.413 | 0 | 11.783 | 0 | 0 |
| g41645.t1 | 19.42 | 59.484 | 83.873 | 16.72 | 1.963 | 2.495 |
| g43969.t1 | 0 | 0.478 | 31.632 | 12.159 | 5.494 | 0.21 |
| g10363.t1 | 9.412 | 17.297 | 81.875 | 137.56 | 664.469 | 135.866 |
| g30255.t1 | 0 | 0.359 | 8.666 | 64.046 | 25.951 | 8.255 |
| g6619.t1 | 3.248 | 4.689 | 148.948 | 221.231 | 40.512 | 83.115 |
| g24040.t1 | 49.67 | 57.348 | 4893.638 | 1600.342 | 648.8 | 836.387 |
| g5615.t1 | 0 | 0 | 16.499 | 24.806 | 0 | 0.669 |
| g8389.t1 | 0.92 | 0.837 | 115.189 | 18.345 | 29.31 | 17.378 |
| g38328.t2 | 0 | 0 | 0 | 2.844 | 1.495 | 2.635 |
| g42890.t1 | 0.548 | 2.665 | 269.038 | 135.772 | 81.368 | 208.421 |
| g34016.t1 | 0 | 0.666 | 399.245 | 30.616 | 17.369 | 5.59 |
| g17719.t1 | 0 | 1.452 | 297.206 | 33.105 | 31.363 | 78.403 |
| g1956.t2 | 0 | 1.025 | 0 | 8.898 | 19.669 | 12.876 |
| g11481.t1 | 4.735 | 12.915 | 45.299 | 68.506 | 270.983 | 170.99 |
| g34894.t1 | 418.367 | 198.464 | 13341.705 | 3326.399 | 2127.433 | 1683.275 |
| g39330.t1 | 0 | 0.256 | 54.21 | 31.449 | 3.827 | 14.523 |
| g5819.t1 | 0.881 | 0.589 | 65.679 | 39.586 | 3.605 | 1.058 |
| g33488.t1 | 221.408 | 88.79 | 4.254 | 8.055 | 5.576 | 22.159 |
| g42803.t1 | 0.636 | 1.034 | 24.058 | 12.677 | 9.518 | 71.417 |
| g18663.t1 | 4.745 | 13.026 | 1487.094 | 240.277 | 190.978 | 122.761 |
| g42397.t1 | 0.137 | 0 | 6.611 | 1.686 | 3.096 | 4.242 |
| g29670.t1 | 1.477 | 2.041 | 96.477 | 42.399 | 19.759 | 12.187 |
| g28843.t2 | 49.435 | 27.231 | 0 | 0 | 0 | 0 |
| g37830.t1 | 96.65 | 70.195 | 3721.773 | 2755.438 | 470.485 | 659.518 |
| g2252.t1 | 1.751 | 2.451 | 113.507 | 37.381 | 19.677 | 14.762 |
| g46809.t1 | 0.92 | 2.195 | 37.424 | 5.942 | 20.851 | 31.202 |
| g14454.t1 | 0.235 | 0.145 | 32.078 | 92.174 | 25.475 | 5.879 |
| g33130.t1 | 129.923 | 113.953 | 7.488 | 2.539 | 0 | 0.689 |
| g17034.t2 | 0 | 0 | 0 | 5.465 | 6.545 | 27.648 |
| g21326.t1 | 0 | 0 | 174.285 | 26.736 | 24.038 | 6.877 |
| g27263.t1 | 25.73 | 71.664 | 1580.366 | 1889.702 | 1010.045 | 194.327 |
| g43199.t1 | 1.262 | 1.281 | 74.733 | 21.626 | 1.174 | 0.689 |
| g21533.t1 | 0 | 0.12 | 4.11 | 1.839 | 2.735 | 2.146 |
| g877.t1 | 0.313 | 0 | 3.722 | 2.072 | 8.779 | 8.853 |
| g40217.t1 | 0 | 0 | 30.468 | 28.93 | 3.605 | 0.23 |
| g6513.t1 | 1.653 | 5.757 | 432.601 | 117.183 | 64.721 | 36.342 |
| g40373.t1 | 1.282 | 1.922 | 488.938 | 74.55 | 23.11 | 4.052 |
| g10332.t1 | 0 | 0.077 | 0 | 0.203 | 1.035 | 1.637 |
| g21911.t1 | 1.242 | 1.016 | 300.497 | 153.589 | 42.762 | 34.066 |
| g39675.t1 | 8.052 | 6.978 | 444.559 | 133.821 | 46.671 | 79.112 |
| g24175.t2 | 0 | 0 | 46.42 | 22.896 | 3.071 | 3.284 |
| g45375.t1 | 0 | 0.171 | 30.64 | 12.291 | 15.932 | 4.891 |
| g32987.t1 | 3.101 | 3.357 | 482.069 | 123.785 | 109.339 | 64.43 |
| g44536.t1 | 0 | 1.384 | 192.48 | 60.257 | 9.058 | 2.475 |
| g37022.t1 | 0 | 0.991 | 367.685 | 5.384 | 36.717 | 18.136 |
| g46293.t1 | 10.048 | 8.234 | 256.304 | 86.231 | 41.185 | 42.85 |
| g43056.t1 | 0.528 | 2.614 | 172.244 | 8.726 | 49.217 | 42.071 |
| g40210.t1 | 0 | 0 | 6.209 | 5.577 | 2.07 | 0.2 |
| g41524.t1 | 0.333 | 0.401 | 1.969 | 0.549 | 8.286 | 5.759 |
| g20527.t1 | 0.519 | 1.213 | 277.388 | 40.398 | 22.157 | 13.255 |
| g44291.t1 | 0.714 | 4.22 | 440.908 | 95.343 | 45.85 | 75.18 |
| g43322.t1 | 34.976 | 106.07 | 14.832 | 0.995 | 0 | 0.988 |
| g44010.t1 | 0.792 | 0.854 | 243.657 | 37.971 | 6.709 | 4.392 |
| g21028.t1 | 0.068 | 0.393 | 123.079 | 37.625 | 37.366 | 42.201 |
| g33158.t1 | 18.432 | 12.326 | 1.466 | 1.9 | 0 | 0.269 |
| g4147.t1 | 4.461 | 1.239 | 370.3 | 91.828 | 25.59 | 12.497 |
| g19158.t1 | 1.536 | 0 | 252.568 | 88.75 | 34.894 | 33.268 |
| g22657.t1 | 0 | 0 | 103.318 | 98.665 | 15.833 | 13.016 |
| g39264.t1 | 0 | 0 | 0 | 0 | 6.406 | 5.989 |
| g37576.t1 | 3.874 | 3.28 | 179.459 | 36.863 | 3.121 | 2.326 |
| g8369.t1 | 3.18 | 2.11 | 144.45 | 54.193 | 63.243 | 26.121 |
| g30555.t1 | 8.062 | 22.396 | 0 | 3.555 | 0 | 0 |
| g12043.t1 | 1190.423 | 1037.087 | 52.931 | 52.933 | 61.962 | 25.622 |
| g16051.t1 | 1.115 | 2.289 | 227.259 | 45.457 | 168.542 | 149.061 |
| g16053.t1 | 0 | 0.29 | 373.433 | 82.036 | 112.641 | 0.519 |
| g18613.t1 | 4.637 | 9.951 | 2320.205 | 8301.288 | 3100.522 | 935.153 |
| g29581.t1 | 3.845 | 3.425 | 58.996 | 19.026 | 93.613 | 160.231 |
| g20444.t1 | 0.646 | 0.521 | 31.977 | 19.209 | 32.209 | 19.184 |
| g10789.t1 | 20.976 | 28.939 | 217.343 | 142.832 | 508.352 | 421.672 |
| g30704.t1 | 5.009 | 4.809 | 201.16 | 60.501 | 31.905 | 27.039 |
| g29097.t1 | 0.881 | 1.512 | 38.545 | 19.006 | 12.606 | 6.308 |
| g4465.t1 | 61.078 | 58.006 | 4189.973 | 547.658 | 147.625 | 182.01 |
| g28538.t1 | 0 | 0.726 | 136.157 | 24.582 | 6.677 | 0.649 |
| g24267.t1 | 2.73 | 1.051 | 66.469 | 53.472 | 10.545 | 8.993 |
| g31519.t1 | 23.451 | 37.13 | 2186.721 | 414.883 | 53.446 | 85.061 |
| g35223.t1 | 0 | 0 | 64.716 | 40.083 | 7.432 | 8.953 |
| g503.t1 | 0.929 | 0.658 | 65.592 | 17.502 | 9.723 | 25.203 |
| g30342.t1 | 0 | 0.777 | 18.899 | 13.784 | 47.328 | 32.3 |
| g15119.t1 | 1.438 | 0 | 0 | 7.334 | 26.936 | 12.926 |
| g13358.t1 | 0 | 0 | 24.777 | 54.203 | 31.847 | 7.536 |
| g33369.t1 | 5.185 | 3.374 | 0 | 0 | 0 | 0 |
| g20198.t1 | 15.624 | 3.041 | 0 | 0 | 0 | 0 |
| g42207.t1 | 4.383 | 35.618 | 7108.427 | 2788.065 | 579.356 | 433.43 |
| g43274.t1 | 0 | 0.982 | 211.105 | 6.003 | 57.413 | 19.983 |
| g29003.t1 | 3.317 | 8.089 | 116.583 | 195.166 | 5.075 | 21.989 |
| g47019.t1 | 1.154 | 1.546 | 1487.755 | 331.262 | 20.547 | 2.306 |
| g18154.t1 | 0.47 | 0.47 | 92.898 | 10.676 | 4.254 | 5.55 |
| g1866.t1 | 22.776 | 19.031 | 689.84 | 469.106 | 279.721 | 261.252 |
| g28114.t1 | 0.225 | 0.273 | 41.405 | 6.907 | 3.318 | 1.467 |
| g24175.t1 | 0 | 0 | 36.447 | 14.018 | 0 | 0 |
| g28520.t1 | 5.361 | 37.284 | 101.68 | 9.579 | 0.452 | 0.09 |
| g33102.t1 | 7.514 | 2.024 | 245.382 | 78.318 | 40.191 | 14.603 |
| g22260.t1 | 377.277 | 299.725 | 2.314 | 6.45 | 0 | 0.21 |
| g39865.t1 | 3.268 | 7.679 | 859.469 | 88.446 | 24.703 | 8.275 |
| g39214.t2 | 1.36 | 0.999 | 20.408 | 14.881 | 38.22 | 27.628 |
| g14001.t1 | 0.695 | 1.23 | 641.767 | 139.093 | 34.089 | 47.172 |
| g7553.t1 | 0.068 | 0.179 | 72.792 | 15.298 | 12.335 | 5.979 |
| g35942.t1 | 0.655 | 3.032 | 365.859 | 158.576 | 88.775 | 105.193 |
| g11275.t1 | 1.008 | 3.314 | 700.949 | 81.477 | 55.852 | 11.658 |
| g14387.t1 | 0 | 0.214 | 182.549 | 38.621 | 24.161 | 13.405 |
| g5158.t1 | 0 | 0.29 | 26.774 | 9.924 | 3.975 | 1.547 |
| g4037.t1 | 0 | 0.538 | 140.339 | 22.124 | 7.481 | 33.957 |
| g35781.t1 | 7.073 | 2.161 | 368.719 | 83.976 | 38.097 | 57.203 |
| g11705.t1 | 1.419 | 10.088 | 16.039 | 11.946 | 130.059 | 79.581 |

**Supplementary table 5 *In silico* expression matrix for *P.sumatrense* root tissue in response to drought and salinity stress**

| Gene ID | Control_Root1 | Control_Root2 | Drought_Root1 | Drought_Root2 | Salt_Root1 | Salt_Root2 |
| --- | --- | --- | --- | --- | --- | --- |
| g8661.t2 | 0 | 0 | 14.325 | 6.163 | 0 | 0 |
| g261.t1 | 18.018 | 23.035 | 23.981 | 19.59 | 0 | 0 |
| g8468.t1 | 4.163 | 5.018 | 1.328 | 2.024 | 0 | 0.363 |
| g10652.t1 | 10.195 | 14.562 | 0 | 0 | 0 | 0 |
| g31852.t1 | 6.499 | 10.526 | 1.635 | 1.429 | 0 | 0.522 |
| g30496.t1 | 13.371 | 9.553 | 2.166 | 2.818 | 0.495 | 0.681 |
| g38606.t1 | 201.79 | 169.019 | 12.578 | 23.291 | 7.038 | 13.499 |
| g32624.t1 | 6.742 | 5.571 | 0 | 0 | 1.023 | 0 |
| g39952.t1 | 5.885 | 5.75 | 12.322 | 12.335 | 0 | 0 |
| g42302.t2 | 0 | 0 | 0 | 0 | 2.867 | 11.558 |
| g25464.t1 | 29.433 | 28.302 | 0 | 0 | 30.13 | 34.957 |
| g17189.t2 | 0 | 0 | 2.442 | 6.54 | 0 | 0 |
| g35496.t1 | 1.696 | 1.759 | 1.686 | 0.496 | 0 | 0 |
| g9375.t1 | 2.008 | 1.536 | 0.296 | 0 | 0 | 0 |
| g44304.t1 | 10.913 | 9.714 | 1.717 | 9.418 | 0 | 0 |
| g24678.t2 | 0 | 0 | 1.502 | 4.545 | 0 | 2.52 |
| g15162.t1 | 9.364 | 6.75 | 3.433 | 4.178 | 0 | 0 |
| g4349.t1 | 62.63 | 72.148 | 29.069 | 30.952 | 2.102 | 1.408 |
| g22882.t1 | 14.894 | 17.035 | 1.185 | 0.595 | 0 | 3.27 |
| g21952.t1 | 4.327 | 4.518 | 2.401 | 1.826 | 0 | 0.329 |
| g45838.t2 | 10.489 | 6.419 | 4.741 | 0 | 0 | 0 |
| g35421.t2 | 28.221 | 4.044 | 0 | 0 | 4.227 | 12.364 |
| g1804.t1 | 2.302 | 4.393 | 1.093 | 0.754 | 0 | 0 |
| g42553.t1 | 4.587 | 4.535 | 0 | 0 | 0 | 0 |
| g44231.t1 | 14.279 | 21.936 | 4.159 | 2.064 | 1.709 | 0 |
| g17201.t1 | 5.167 | 7.482 | 8.665 | 9.388 | 0 | 0 |
| g12269.t1 | 16.573 | 13.383 | 8.583 | 6.986 | 0.27 | 0.193 |
| g28843.t2 | 29.883 | 18.267 | 4.843 | 0 | 0.472 | 0 |
| g32195.t1 | 6.318 | 4.446 | 2.054 | 0 | 0 | 0 |
| g14634.t1 | 3.116 | 3.357 | 0 | 0 | 0 | 0.272 |
| g34703.t2 | 3.323 | 4.768 | 1.349 | 4.079 | 0 | 0 |
| g30561.t1 | 0 | 0 | 0 | 3.384 | 7.6 | 3.713 |
| g25082.t1 | 21.861 | 9.901 | 0 | 0.724 | 1.169 | 0 |
| g19604.t1 | 0 | 0 | 0.633 | 1.28 | 13.356 | 9.162 |
| g31762.t1 | 2.198 | 4.616 | 0.204 | 0.853 | 0.326 | 0 |
| g36922.t2 | 11.121 | 10.857 | 3.75 | 6.341 | 0 | 0 |
| g13468.t1 | 4.561 | 3.277 | 2.074 | 2.104 | 0 | 0.329 |
| g42660.t1 | 20.009 | 35.766 | 0.429 | 2.62 | 0 | 0.477 |
| g7589.t1 | 4.128 | 3.205 | 2.789 | 1.538 | 0 | 0.284 |
| g13610.t1 | 13.587 | 22.276 | 24.165 | 0 | 0 | 0 |
| g6522.t1 | 4.18 | 3.902 | 0.286 | 3.533 | 0.461 | 0 |
| g39214.t1 | 4.855 | 3.911 | 3.852 | 0.715 | 0 | 0 |
| g25249.t2 | 0 | 0 | 5.671 | 0 | 114.877 | 29.575 |
| g8999.t1 | 10.299 | 14.071 | 4.107 | 1.181 | 0 | 0 |
| g36303.t1 | 0 | 0 | 4.823 | 15.074 | 2.575 | 12.114 |
| g33577.t2 | 4.838 | 1.268 | 1.635 | 1.628 | 0 | 0 |
| g30702.t1 | 42.189 | 33.811 | 0.828 | 2.501 | 1.979 | 1.828 |
| g24085.t2 | 5.002 | 9.25 | 0 | 0.258 | 3.058 | 5.336 |
| g36713.t1 | 1.402 | 2.803 | 1.236 | 2.838 | 0.124 | 0 |
| g34518.t2 | 0 | 0 | 0.868 | 0 | 0.72 | 1.135 |
| g44960.t1 | 37.983 | 24.776 | 32.737 | 27.707 | 0 | 0 |
| g3894.t1 | 48.455 | 46.435 | 15.255 | 9.517 | 0 | 0.329 |
| g25048.t1 | 32.22 | 31.15 | 9.513 | 16.007 | 0 | 1.499 |
| g28517.t1 | 5.885 | 5.75 | 12.322 | 12.335 | 0 | 0 |
| g22204.t1 | 16.374 | 17.267 | 0 | 1.201 | 1.259 | 0 |
| g36146.t1 | 14.885 | 11.973 | 1.89 | 1.528 | 0.596 | 0.829 |
| g40517.t1 | 42.587 | 21.249 | 2.135 | 0 | 0 | 0 |
| g33402.t1 | 2.068 | 0.893 | 0.685 | 0 | 0 | 0 |
| g29884.t1 | 1.627 | 1.214 | 1.073 | 0.724 | 0 | 0.079 |
| g43180.t1 | 2.466 | 2.482 | 1.717 | 1.737 | 0 | 0 |
| g18586.t1 | 2.293 | 2.027 | 0.858 | 1.389 | 0 | 0.193 |
| g35695.t2 | 0 | 0 | 3.535 | 10.926 | 0 | 0 |
| g18345.t1 | 2.406 | 3.848 | 1.543 | 1.628 | 0 | 0 |
| g45627.t1 | 5.504 | 3.946 | 0.593 | 0.506 | 0 | 0.272 |
| g15110.t2 | 5.097 | 3.955 | 0 | 3.384 | 0 | 0 |
| g23208.t1 | 2.215 | 2.062 | 2.943 | 0 | 0 | 0 |
| g20501.t1 | 0 | 0 | 1.379 | 1.28 | 0.708 | 1.998 |
| g13840.t1 | 9.139 | 9.491 | 12.68 | 3.92 | 1.124 | 0.193 |
| g46716.t1 | 3.306 | 1.812 | 1.318 | 0.675 | 0 | 0 |
| g6357.t1 | 137.792 | 139.725 | 9.523 | 6.748 | 10.062 | 3.508 |
| g32329.t1 | 5.089 | 4.446 | 1.614 | 2.302 | 0.506 | 0 |
| g3840.t1 | 17.187 | 16.338 | 8.797 | 11.393 | 0 | 0.976 |
| g19989.t1 | 8.49 | 5.955 | 1.757 | 0.893 | 0 | 0.488 |
| g34961.t1 | 0 | 0.482 | 5.528 | 4.664 | 2.428 | 6.437 |
| g34518.t1 | 1.16 | 1.321 | 0 | 1.191 | 0 | 0 |
| g26115.t1 | 6.811 | 3.509 | 2.37 | 2.402 | 0 | 0 |
| g42211.t1 | 31.415 | 35.954 | 3.004 | 2.411 | 1.99 | 0.874 |
| g1251.t1 | 0.831 | 0 | 0 | 0 | 8.308 | 17.836 |
| g36067.t1 | 7.477 | 3.893 | 0 | 7.86 | 0 | 0 |
| g8523.t1 | 47.313 | 42.703 | 3.065 | 1.538 | 2.451 | 1.124 |
| g30378.t1 | 2.717 | 1.83 | 2.738 | 0 | 0 | 0 |
| g28349.t1 | 2.596 | 5.616 | 2.36 | 2.392 | 0 | 0 |
| g45298.t1 | 8.905 | 9.633 | 0.296 | 0.228 | 0.697 | 0.579 |
| g32737.t2 | 6.993 | 5.839 | 0.552 | 1.072 | 0 | 0.114 |
| g32899.t1 | 2.363 | 1.83 | 1.165 | 1.032 | 0.225 | 0 |
| g7727.t1 | 2.155 | 2.661 | 1.88 | 0.576 | 0 | 0 |
| g15287.t1 | 0.831 | 1.562 | 0.123 | 0 | 0 | 0 |
| g153.t1 | 21.203 | 16.303 | 0.746 | 1.131 | 0 | 0 |
| g22524.t1 | 3.851 | 2.839 | 3.157 | 6.014 | 0 | 0 |
| g42920.t1 | 0 | 0 | 733.123 | 0 | 445.6 | 35.49 |
| g43210.t1 | 4.059 | 3.794 | 1.124 | 2.58 | 0 | 0.307 |
| g14303.t1 | 25.729 | 25.82 | 0.756 | 0.863 | 1.911 | 1.862 |
| g35929.t1 | 2.882 | 3.732 | 0.858 | 0.873 | 0 | 0 |
| g5009.t1 | 2.544 | 2.098 | 3.443 | 1.618 | 0 | 0 |
| g32830.t1 | 0 | 0.259 | 1.952 | 0.992 | 7.229 | 5.393 |
| g20376.t1 | 45.824 | 44.31 | 1.073 | 1.35 | 2.114 | 2.952 |
| g17941.t1 | 3.955 | 4.491 | 3.137 | 3.195 | 0 | 0 |
| g29750.t1 | 6.958 | 5.027 | 2.708 | 4.555 | 0 | 0 |
| g7004.t1 | 6.499 | 2.411 | 0 | 0 | 0 | 0 |
| g24913.t1 | 4.942 | 3.053 | 0 | 1.3 | 0 | 0.352 |
| g42497.t1 | 2.129 | 2.625 | 0 | 0 | 0 | 3.928 |
| g28126.t1 | 4.673 | 5.777 | 0.347 | 0 | 0 | 0.772 |
| g6242.t2 | 4.18 | 1.92 | 0 | 0 | 0 | 0.715 |
| g11021.t1 | 2.077 | 1.848 | 0 | 0 | 0 | 0 |
| g21385.t1 | 2.562 | 4.714 | 3.637 | 1.846 | 0.405 | 0 |
| g38545.t1 | 2.12 | 1.982 | 0.889 | 0.456 | 0 | 0.159 |
| g19034.t1 | 0 | 0 | 62.338 | 46.017 | 0 | 63.169 |
| g42302.t1 | 7.183 | 8.178 | 11.025 | 15.114 | 0 | 0 |
| g42119.t2 | 0 | 0 | 4.833 | 3.007 | 0 | 6.96 |
| g17189.t1 | 7.036 | 5.973 | 0 | 0 | 1.057 | 3.94 |
| g4988.t1 | 1.532 | 3.803 | 3.28 | 1.657 | 0 | 0 |
| g40678.t1 | 4.163 | 4.411 | 1.543 | 0.625 | 0 | 0 |
| g21428.t1 | 6.499 | 7.892 | 4.271 | 1.796 | 0 | 0.397 |
| g26965.t1 | 4.379 | 3.027 | 0.347 | 2.769 | 0 | 0 |

**Supplementary table 6. Uniprot database based annotation for differentially expressed *P.sumatrense* genes in root tissue in response to drought and salinity stress**

| g153.t1 | sp_Q6K9A2_CCR1_ORYSJ_Cinnamoyl-CoA_reductase_1_OS=Oryza_sativa_subsp._japonica_OX=39947_GN=CCR1_PE=1_SV=1 |
| --- | --- |
| g261.t1 | sp_Q5SMM8_PHT1_ORYSJ_Putrescine_hydroxycinnamoyltransferase_1_OS=Oryza_sativa_subsp._japonica_OX=39947_GN=PHT1_PE=2_SV=1 |
| g1251.t1 | sp_Q9ZUH0_FBK35_ARATH_F-box/kelch-repeat_protein_At2g24250_OS=Arabidopsis_thaliana_OX=3702_GN=At2g24250_PE=2_SV=2 |
| g1804.t1 | sp_Q6Z8L2_LAC9_ORYSJ_Putative_laccase-9_OS=Oryza_sativa_subsp._japonica_OX=39947_GN=LAC9_PE=3_SV=1 |
| g3840.t1 | sp_Q2L6L1_CNPY1_DANRE_Protein_canopy-1_OS=Danio_rerio_OX=7955_GN=cnpy1_PE=1_SV=1 |
| g3894.t1 | sp_P22988_LDHA_HORVU_L-lactate_dehydrogenase_A_OS=Hordeum_vulgare_OX=4513_PE=1_SV=1 |
| g4349.t1 | sp_P24632_HSP22_MAIZE_17.8_kDa_class_II_heat_shock_protein_OS=Zea_mays_OX=4577_PE=2_SV=1 |
| g4988.t1 | sp_Q52RG7_SGPL_ORYSJ_Sphingosine-1-phosphate_lyase_OS=Oryza_sativa_subsp._japonica_OX=39947_GN=SPL_PE=2_SV=3 |
| g4988.t1 | sp_Q52RG7_SGPL_ORYSJ_Sphingosine-1-phosphate_lyase_OS=Oryza_sativa_subsp._japonica_OX=39947_GN=SPL_PE=2_SV=3 |
| g5009.t1 | sp_Q8L5Z1_GDL17_ARATH_GDSL_esterase/lipase_At1g33811_OS=Arabidopsis_thaliana_OX=3702_GN=At1g33811_PE=2_SV=1 |
| g6242.t2 | sp_Q8S0F0_FH1_ORYSJ_Formin-like_protein_1_OS=Oryza_sativa_subsp._japonica_OX=39947_GN=FH1_PE=2_SV=1 |
| g6242.t2 | sp_Q8S0F0_FH1_ORYSJ_Formin-like_protein_1_OS=Oryza_sativa_subsp._japonica_OX=39947_GN=FH1_PE=2_SV=1 |
| g6242.t2 | sp_Q8S0F0_FH1_ORYSJ_Formin-like_protein_1_OS=Oryza_sativa_subsp._japonica_OX=39947_GN=FH1_PE=2_SV=1 |
| g6357.t1 | sp_Q8LJJ9_STAD1_ORYSJ_Stearoyl-[acyl-carrier-protein]_9-desaturase_1,_chloroplastic_OS=Oryza_sativa_subsp._japonica_OX=39947_GN=Os01g0880800_PE=2_SV=1 |
| g6522.t1 | sp_Q5YDB5_CPL2_ARATH_RNA_polymerase_II_C-terminal_domain_phosphatase-like_2_OS=Arabidopsis_thaliana_OX=3702_GN=CPL2_PE=1_SV=3 |
| g7004.t1 | sp_Q8L7I2_SBT36_ARATH_Subtilisin-like_protease_SBT3.6_OS=Arabidopsis_thaliana_OX=3702_GN=SBT3.6_PE=2_SV=1 |
| g7004.t1 | sp_Q8L7I2_SBT36_ARATH_Subtilisin-like_protease_SBT3.6_OS=Arabidopsis_thaliana_OX=3702_GN=SBT3.6_PE=2_SV=1 |
| g7589.t1 | sp_O74777_KRR1_SCHPO_KRR1_small_subunit_processome_component_homolog_OS=Schizosaccharomyces_pombe_(strain_972_/_ATCC_24843)_OX=284812_GN=mis3_PE=3_SV=1 |
| g7727.t1 | sp_Q9FR44_PEAM1_ARATH_Phosphoethanolamine_N-methyltransferase_1_OS=Arabidopsis_thaliana_OX=3702_GN=NMT1_PE=1_SV=1 |
| g8468.t1 | sp_Q9D6J3_YJU2_MOUSE_YJU2_splicing_factor_homolog_OS=Mus_musculus_OX=10090_GN=Yju2_PE=1_SV=1 |
| g8523.t1 | sp_Q9ZUN4_ACCO1_ARATH_1-aminocyclopropane-1-carboxylate_oxidase_1_OS=Arabidopsis_thaliana_OX=3702_GN=ACO1_PE=2_SV=1 |
| g8661.t2 | sp_Q6YW46_EF1G2_ORYSJ_Elongation_factor_1-gamma_2_OS=Oryza_sativa_subsp._japonica_OX=39947_GN=Os02g0220500_PE=2_SV=2 |
| g8999.t1 | sp_O23193_CBSX1_ARATH_CBS_domain-containing_protein_CBSX1,_chloroplastic_OS=Arabidopsis_thaliana_OX=3702_GN=CBSX1_PE=1_SV=2 |
| g9375.t1 | sp_Q43207_FKB70_WHEAT_70_kDa_peptidyl-prolyl_isomerase_OS=Triticum_aestivum_OX=4565_GN=FKBP70_PE=1_SV=1 |
| g10652.t1 | sp_P16347_IAAS_WHEAT_Endogenous_alpha-amylase/subtilisin_inhibitor_OS=Triticum_aestivum_OX=4565_PE=1_SV=1 |
| g11021.t1 | sp_Q7XU38_C87A3_ORYSJ_Cytochrome_P450_87A3_OS=Oryza_sativa_subsp._japonica_OX=39947_GN=CYP87A3_PE=2_SV=3 |
| g12269.t1 | sp_Q94EE9_TYDC_ORYSJ_Tyrosine_decarboxylase_OS=Oryza_sativa_subsp._japonica_OX=39947_GN=TYDC_PE=1_SV=1 |
| g13468.t1 | sp_P22988_LDHA_HORVU_L-lactate_dehydrogenase_A_OS=Hordeum_vulgare_OX=4513_PE=1_SV=1 |
| g13610.t1 | sp_Q10I20_XAT3_ORYSJ_Alpha-1,3-arabinosyltransferase_XAT3_OS=Oryza_sativa_subsp._japonica_OX=39947_GN=XAT3_PE=1_SV=1 |
| g13840.t1 | sp_Q9SS90_RGLG1_ARATH_E3_ubiquitin-protein_ligase_RGLG1_OS=Arabidopsis_thaliana_OX=3702_GN=RGLG1_PE=1_SV=1 |
| g14303.t1 | sp_Q5ZAJ0_RBOHB_ORYSJ_Respiratory_burst_oxidase_homolog_protein_B_OS=Oryza_sativa_subsp._japonica_OX=39947_GN=RBOHB_PE=1_SV=1 |
| g14634.t1 | sp_Q9S850_SUOX_ARATH_Sulfite_oxidase_OS=Arabidopsis_thaliana_OX=3702_GN=SOX_PE=1_SV=1 |
| g15110.t2 | sp_P27898_MYBP_MAIZE_Myb-related_protein_P_OS=Zea_mays_OX=4577_GN=P_PE=2_SV=1 |
| g15162.t1 | sp_A5H8G4_PER1_MAIZE_Peroxidase_1_OS=Zea_mays_OX=4577_GN=PER1_PE=1_SV=1 |
| g15162.t1 | sp_A5H8G4_PER1_MAIZE_Peroxidase_1_OS=Zea_mays_OX=4577_GN=PER1_PE=1_SV=1 |
| g15287.t1 | sp_Q9CAP8_LACS9_ARATH_Long_chain_acyl-CoA_synthetase_9,_chloroplastic_OS=Arabidopsis_thaliana_OX=3702_GN=LACS9_PE=1_SV=1 |
| g17189.t1 | sp_O31222_DRNG_AERHY_Extracellular_deoxyribonuclease_OS=Aeromonas_hydrophila_OX=644_PE=3_SV=2 |
| g17189.t2 | sp_O31222_DRNG_AERHY_Extracellular_deoxyribonuclease_OS=Aeromonas_hydrophila_OX=644_PE=3_SV=2 |
| g17201.t1 | sp_Q7XC27_CNBL1_ORYSJ_Calcineurin_B-like_protein_1_OS=Oryza_sativa_subsp._japonica_OX=39947_GN=CBL1_PE=2_SV=2 |
| g17941.t1 | sp_Q06215_PPO_VICFA_Polyphenol_oxidase_A1,_chloroplastic_OS=Vicia_faba_OX=3906_PE=1_SV=1 |
| g18345.t1 | sp_Q6ZK48_SLU7_ORYSJ_Pre-mRNA-splicing_factor_SLU7_OS=Oryza_sativa_subsp._japonica_OX=39947_GN=Os08g0127700_PE=2_SV=1 |
| g18586.t1 | sp_Q94EE9_TYDC_ORYSJ_Tyrosine_decarboxylase_OS=Oryza_sativa_subsp._japonica_OX=39947_GN=TYDC_PE=1_SV=1 |
| g19034.t1 | sp_Q9SX65_DNAT1_ARATH_1,4-dihydroxy-2-naphthoyl-CoA_thioesterase_1_OS=Arabidopsis_thaliana_OX=3702_GN=DHNAT1_PE=1_SV=1 |
| g19604.t1 | sp_Q9SGA8_U83A1_ARATH_UDP-glycosyltransferase_83A1_OS=Arabidopsis_thaliana_OX=3702_GN=UGT83A1_PE=2_SV=1 |
| g19989.t1 | sp_Q8L970_P4H7_ARATH_Probable_prolyl_4-hydroxylase_7_OS=Arabidopsis_thaliana_OX=3702_GN=P4H7_PE=2_SV=1 |
| g20376.t1 | sp_P04707_ADH2_MAIZE_Alcohol_dehydrogenase_2_OS=Zea_mays_OX=4577_GN=ADH2_PE=2_SV=1 |
| g20501.t1 | sp_Q9M175_PTR39_ARATH_Protein_NRT1/_PTR_FAMILY_2.3_OS=Arabidopsis_thaliana_OX=3702_GN=NPF2.3_PE=2_SV=1 |
| g21385.t1 | sp_Q9SH89_OFT19_ARATH_O-fucosyltransferase_19_OS=Arabidopsis_thaliana_OX=3702_GN=OFUT19_PE=2_SV=1 |
| g21385.t1 | sp_Q9SH89_OFT19_ARATH_O-fucosyltransferase_19_OS=Arabidopsis_thaliana_OX=3702_GN=OFUT19_PE=2_SV=1 |
| g21428.t1 | sp_B2C6R6_TAFCL_ARATH_Transcription_initiation_factor_TFIID_subunit_12b_OS=Arabidopsis_thaliana_OX=3702_GN=TAF12B_PE=1_SV=1 |
| g21952.t1 | sp_Q7XIM0_CDPKH_ORYSJ_Calcium-dependent_protein_kinase_17_OS=Oryza_sativa_subsp._japonica_OX=39947_GN=CPK17_PE=2_SV=1 |
| g22204.t1 | sp_Q8L970_P4H7_ARATH_Probable_prolyl_4-hydroxylase_7_OS=Arabidopsis_thaliana_OX=3702_GN=P4H7_PE=2_SV=1 |
| g22524.t1 | sp_Q8N5L8_RP25L_HUMAN_Ribonuclease_P_protein_subunit_p25-like_protein_OS=Homo_sapiens_OX=9606_GN=RPP25L_PE=1_SV=1 |
| g22882.t1 | sp_Q851K0_GL35_ORYSJ_Germin-like_protein_3-5_OS=Oryza_sativa_subsp._japonica_OX=39947_GN=Os03g0693900_PE=2_SV=1 |
| g23208.t1 | sp_Q10CH5_PIL13_ORYSJ_Transcription_factor_PHYTOCHROME_INTERACTING_FACTOR-LIKE_13_OS=Oryza_sativa_subsp._japonica_OX=39947_GN=PIL13_PE=1_SV=1 |
| g23208.t1 | sp_Q10CH5_PIL13_ORYSJ_Transcription_factor_PHYTOCHROME_INTERACTING_FACTOR-LIKE_13_OS=Oryza_sativa_subsp._japonica_OX=39947_GN=PIL13_PE=1_SV=1 |
| g24085.t2 | sp_Q9FYC2_PAO_ARATH_Pheophorbide_a_oxygenase,_chloroplastic_OS=Arabidopsis_thaliana_OX=3702_GN=PAO_PE=1_SV=1 |
| g24678.t2 | sp_Q9ZU91_E133_ARATH_Glucan_endo-1,3-beta-glucosidase_3_OS=Arabidopsis_thaliana_OX=3702_GN=At2g01630_PE=2_SV=2 |
| g24913.t1 | sp_Q9C6T1_PMK_ARATH_Phosphomevalonate_kinase,_peroxisomal_OS=Arabidopsis_thaliana_OX=3702_GN=PMK_PE=1_SV=1 |
| g25048.t1 | sp_Q84Q72_HS181_ORYSJ_18.1_kDa_class_I_heat_shock_protein_OS=Oryza_sativa_subsp._japonica_OX=39947_GN=HSP18.1_PE=2_SV=1 |
| g25082.t1 | sp_B8E548_RNFA_SHEB2_Ion-translocating_oxidoreductase_complex_subunit_A_OS=Shewanella_baltica_(strain_OS223)_OX=407976_GN=rnfA_PE=3_SV=1 |
| g25249.t2 | sp_Q10MX3_ASNS1_ORYSJ_Asparagine_synthetase_[glutamine-hydrolyzing]_1_OS=Oryza_sativa_subsp._japonica_OX=39947_GN=Os03g0291500_PE=2_SV=1 |
| g25464.t1 | sp_Q10MB4_MYB2_ORYSJ_Transcription_factor_MYB2_OS=Oryza_sativa_subsp._japonica_OX=39947_GN=MYB2_PE=2_SV=1 |
| g26115.t1 | sp_P19182_IFRD1_MOUSE_Interferon-related_developmental_regulator_1_OS=Mus_musculus_OX=10090_GN=Ifrd1_PE=1_SV=2 |
| g26965.t1 | sp_A7PZL3_PGLR_VITVI_Probable_polygalacturonase_OS=Vitis_vinifera_OX=29760_GN=GSVIVT00026920001_PE=1_SV=1 |
| g28126.t1 | sp_Q945B6_AOP1L_ARATH_Probable_2-oxoglutarate-dependent_dioxygenase_AOP1.2_OS=Arabidopsis_thaliana_OX=3702_GN=AOP1.2_PE=2_SV=1 |
| g28349.t1 | sp_Q5Z5C9_BURPB_ORYSJ_BURP_domain-containing_protein_11_OS=Oryza_sativa_subsp._japonica_OX=39947_GN=BURP11_PE=2_SV=1 |
| g28517.t1 | sp_P43309_PPO_MALDO_Polyphenol_oxidase,_chloroplastic_OS=Malus_domestica_OX=3750_PE=2_SV=1 |
| g28843.t2 | sp_P43284_TRPB2_MAIZE_Tryptophan_synthase_beta_chain_2,_chloroplastic_(Fragment)_OS=Zea_mays_OX=4577_GN=TSB2_PE=2_SV=1 |
| g29750.t1 | sp_Q2QZ14_GLO1B_ORYSJ_Very-long-chain_aldehyde_decarbonylase_GL1-11_OS=Oryza_sativa_subsp._japonica_OX=39947_GN=GL1-11_PE=2_SV=1 |
| g29884.t1 | sp_P93026_VSR1_ARATH_Vacuolar-sorting_receptor_1_OS=Arabidopsis_thaliana_OX=3702_GN=VSR1_PE=1_SV=2 |
| g30378.t1 | sp_P73627_Y1770_SYNY3_Uncharacterized_protein_sll1770_OS=Synechocystis_sp._(strain_PCC_6803_/_Kazusa)_OX=1111708_GN=sll1770_PE=3_SV=1 |
| g30496.t1 | sp_Q00799_RBP2_PLAVB_Reticulocyte-binding_protein_2_OS=Plasmodium_vivax_(strain_Belem)_OX=31273_GN=RBP-2_PE=3_SV=2 |
| g30561.t1 | sp_Q9SII6_PIX13_ARATH_Probable_serine/threonine-protein_kinase_PIX13_OS=Arabidopsis_thaliana_OX=3702_GN=PIX13_PE=1_SV=2 |
| g30702.t1 | sp_Q9ZUN4_ACCO1_ARATH_1-aminocyclopropane-1-carboxylate_oxidase_1_OS=Arabidopsis_thaliana_OX=3702_GN=ACO1_PE=2_SV=1 |
| g31762.t1 | sp_Q80Y55_BSDC1_MOUSE_BSD_domain-containing_protein_1_OS=Mus_musculus_OX=10090_GN=Bsdc1_PE=1_SV=1 |
| g31852.t1 | sp_Q5F3L3_F234B_CHICK_Protein_FAM234B_OS=Gallus_gallus_OX=9031_GN=FAM234B_PE=2_SV=1 |
| g32195.t1 | sp_F4IFC5_SYTM2_ARATH_Threonine--tRNA_ligase,_chloroplastic/mitochondrial_2_OS=Arabidopsis_thaliana_OX=3702_GN=EMB2761_PE=2_SV=1 |
| g32329.t1 | sp_Q9FK30_OFT36_ARATH_O-fucosyltransferase_36_OS=Arabidopsis_thaliana_OX=3702_GN=OFUT36_PE=2_SV=1 |
| g32624.t1 | sp_Q3ITB6_COFE_NATPD_Coenzyme_F420:L-glutamate_ligase_OS=Natronomonas_pharaonis_(strain_ATCC_35678_/_DSM_2160_/_CIP_103997_/_NBRC_14720_/_NCIMB_2260_/_Gabara)_OX=348780_GN=cofE_PE=3_SV=1 |
| g32737.t2 | sp_Q9MA92_FPP3_ARATH_Filament-like_plant_protein_3_OS=Arabidopsis_thaliana_OX=3702_GN=FPP3_PE=3_SV=2 |
| g32737.t2 | sp_Q9MA92_FPP3_ARATH_Filament-like_plant_protein_3_OS=Arabidopsis_thaliana_OX=3702_GN=FPP3_PE=3_SV=2 |
| g32830.t1 | sp_Q9FGY4_FB341_ARATH_F-box_protein_At5g49610_OS=Arabidopsis_thaliana_OX=3702_GN=At5g49610_PE=1_SV=1 |
| g32899.t1 | sp_Q9LMX7_C78A5_ARATH_Cytochrome_P450_78A5_OS=Arabidopsis_thaliana_OX=3702_GN=CYP78A5_PE=2_SV=1 |
| g33402.t1 | sp_O53732_UFAA1_MYCTU_Tuberculostearic_acid_methyltransferase_UfaA1_OS=Mycobacterium_tuberculosis_(strain_ATCC_25618_/_H37Rv)_OX=83332_GN=ufaA1_PE=1_SV=3 |
| g33577.t2 | sp_Q9LT38_UNCL_ARATH_Serine/threonine-protein_kinase_UCNL_OS=Arabidopsis_thaliana_OX=3702_GN=UCNL_PE=2_SV=1 |
| g34518.t1 | sp_A3BXL8_AB53G_ORYSJ_ABC_transporter_G_family_member_53_OS=Oryza_sativa_subsp._japonica_OX=39947_GN=ABCG53_PE=2_SV=1 |
| g34518.t2 | sp_A3BXL8_AB53G_ORYSJ_ABC_transporter_G_family_member_53_OS=Oryza_sativa_subsp._japonica_OX=39947_GN=ABCG53_PE=2_SV=1 |
| g34703.t2 | sp_Q94C11_SUGP1_ARATH_SURP_and_G-patch_domain-containing_protein_1-like_protein_OS=Arabidopsis_thaliana_OX=3702_GN=At3g52120_PE=2_SV=1 |
| g34961.t1 | sp_Q8LGZ9_G2OX5_ORYSJ_Gibberellin_2-beta-dioxygenase_5_OS=Oryza_sativa_subsp._japonica_OX=39947_GN=GA2OX5_PE=1_SV=1 |
| g35421.t2 | sp_Q9LH39_PHSD_ARATH_Probable_polyamine_transporter_At3g19553_OS=Arabidopsis_thaliana_OX=3702_GN=At3g19553_PE=3_SV=1 |
| g35496.t1 | sp_Q5XVA8_Y3905_ARATH_Uncharacterized_protein_At3g49055_OS=Arabidopsis_thaliana_OX=3702_GN=At3g49055_PE=2_SV=1 |
| g35695.t2 | sp_A2ZF66_GT6_ORYSI_Probable_glycosyltransferase_6_OS=Oryza_sativa_subsp._indica_OX=39946_GN=GT6_PE=3_SV=1 |
| g35929.t1 | sp_Q9SLY8_CALR_ORYSJ_Calreticulin_OS=Oryza_sativa_subsp._japonica_OX=39947_GN=Os07g0246200_PE=1_SV=2 |
| g36067.t1 | sp_O80674_BH106_ARATH_Transcription_factor_bHLH106_OS=Arabidopsis_thaliana_OX=3702_GN=BHLH106_PE=2_SV=1 |
| g36146.t1 | sp_Q8L970_P4H7_ARATH_Probable_prolyl_4-hydroxylase_7_OS=Arabidopsis_thaliana_OX=3702_GN=P4H7_PE=2_SV=1 |
| g36303.t1 | sp_O80612_APY6_ARATH_Probable_apyrase_6_OS=Arabidopsis_thaliana_OX=3702_GN=APY6_PE=2_SV=2 |
| g36713.t1 | sp_Q6Z8L2_LAC9_ORYSJ_Putative_laccase-9_OS=Oryza_sativa_subsp._japonica_OX=39947_GN=LAC9_PE=3_SV=1 |
| g36922.t2 | sp_Q9FJ45_GDL83_ARATH_GDSL_esterase/lipase_At5g45910_OS=Arabidopsis_thaliana_OX=3702_GN=At5g45910_PE=2_SV=1 |
| g38545.t1 | sp_Q8R149_BUD13_MOUSE_BUD13_homolog_OS=Mus_musculus_OX=10090_GN=Bud13_PE=1_SV=1 |
| g38606.t1 | sp_Q02921_NO93_SOYBN_Early_nodulin-93_OS=Glycine_max_OX=3847_PE=2_SV=1 |
| g39214.t1 | sp_Q653P0_KOR1_ORYSJ_Potassium_channel_KOR1_OS=Oryza_sativa_subsp._japonica_OX=39947_GN=Os06g0250600_PE=2_SV=1 |
| g39952.t1 | sp_P43309_PPO_MALDO_Polyphenol_oxidase,_chloroplastic_OS=Malus_domestica_OX=3750_PE=2_SV=1 |
| g40517.t1 | sp_A9WMW3_FTHS_RENSM_Formate--tetrahydrofolate_ligase_OS=Renibacterium_salmoninarum_(strain_ATCC_33209_/_DSM_20767_/_JCM_11484_/_NBRC_15589_/_NCIMB_2235)_OX=288705_GN=fhs_PE=3_SV=1 |
| g40678.t1 | sp_Q9LI61_AVT6A_ARATH_Amino_acid_transporter_AVT6A_OS=Arabidopsis_thaliana_OX=3702_GN=AVT6A_PE=2_SV=1 |
| g40678.t1 | sp_Q9LI61_AVT6A_ARATH_Amino_acid_transporter_AVT6A_OS=Arabidopsis_thaliana_OX=3702_GN=AVT6A_PE=2_SV=1 |
| g42119.t2 | sp_C0LGD6_Y1570_ARATH_Probable_LRR_receptor-like_serine/threonine-protein_kinase_At1g05700_OS=Arabidopsis_thaliana_OX=3702_GN=At1g05700_PE=1_SV=1 |
| g42211.t1 | sp_Q0DHF6_PDC1_ORYSJ_Pyruvate_decarboxylase_1_OS=Oryza_sativa_subsp._japonica_OX=39947_GN=PDC1_PE=2_SV=1 |
| g42302.t2 | sp_Q9SKD9_WRK46_ARATH_Probable_WRKY_transcription_factor_46_OS=Arabidopsis_thaliana_OX=3702_GN=WRKY46_PE=1_SV=1 |
| g42302.t1 | sp_Q9SKD9_WRK46_ARATH_Probable_WRKY_transcription_factor_46_OS=Arabidopsis_thaliana_OX=3702_GN=WRKY46_PE=1_SV=1 |
| g42497.t1 | sp_Q94F40_GDL9_ARATH_GDSL_esterase/lipase_At1g28600_OS=Arabidopsis_thaliana_OX=3702_GN=At1g28600_PE=2_SV=1 |
| g42553.t1 | sp_P93732_PIP_ARATH_Proline_iminopeptidase_OS=Arabidopsis_thaliana_OX=3702_GN=PIP_PE=2_SV=3 |
| g42660.t1 | sp_Q65X92_C3H37_ORYSJ_Zinc_finger_CCCH_domain-containing_protein_37_OS=Oryza_sativa_subsp._japonica_OX=39947_GN=Os05g0525900_PE=2_SV=1 |
| g42920.t1 | sp_O65101_PSAH_MAIZE_Photosystem_I_reaction_center_subunit_VI,_chloroplastic_OS=Zea_mays_OX=4577_GN=PSAH_PE=1_SV=1 |
| g43180.t1 | sp_Q6L5C4_P2C52_ORYSJ_Probable_protein_phosphatase_2C_52_OS=Oryza_sativa_subsp._japonica_OX=39947_GN=Os05g0587100_PE=2_SV=1 |
| g43180.t1 | sp_Q6L5C4_P2C52_ORYSJ_Probable_protein_phosphatase_2C_52_OS=Oryza_sativa_subsp._japonica_OX=39947_GN=Os05g0587100_PE=2_SV=1 |
| g43210.t1 | sp_Q9FY51_HPAT3_ARATH_Hydroxyproline_O-arabinosyltransferase_3_OS=Arabidopsis_thaliana_OX=3702_GN=HPAT3_PE=1_SV=1 |
| g44231.t1 | sp_Q2RBF0_CIPKF_ORYSJ_CBL-interacting_protein_kinase_15_OS=Oryza_sativa_subsp._japonica_OX=39947_GN=CIPK15_PE=2_SV=1 |
| g44304.t1 | sp_B9DHD7_VAP22_ARATH_Vesicle-associated_protein_2-2_OS=Arabidopsis_thaliana_OX=3702_GN=PVA22_PE=1_SV=1 |
| g45298.t1 | sp_Q5ZAJ0_RBOHB_ORYSJ_Respiratory_burst_oxidase_homolog_protein_B_OS=Oryza_sativa_subsp._japonica_OX=39947_GN=RBOHB_PE=1_SV=1 |
| g45627.t1 | sp_Q9S850_SUOX_ARATH_Sulfite_oxidase_OS=Arabidopsis_thaliana_OX=3702_GN=SOX_PE=1_SV=1 |
| g45838.t2 | sp_Q7ZVZ7_AB17C_DANRE_Alpha/beta_hydrolase_domain-containing_protein_17C_OS=Danio_rerio_OX=7955_GN=abhd17c_PE=2_SV=1 |
| g46716.t1 | sp_Q38931_FKB62_ARATH_Peptidyl-prolyl_cis-trans_isomerase_FKBP62_OS=Arabidopsis_thaliana_OX=3702_GN=FKBP62_PE=1_SV=2 |

**Supplementary table 7. Uniprot database based annotation for differentially expressed *P.sumatrense* genes in leaf tissue in response to drought and salinity stress**

| g63.t1 | sp_Q9FFA5_RMR14_ARATH_Remorin_1.4_OS=Arabidopsis_thaliana_OX=3702_GN=REM1.4_PE=2_SV=1 |
| --- | --- |
| g196.t1 | sp_Q84K16_AP1G1_ARATH_AP-1_complex_subunit_gamma-1_OS=Arabidopsis_thaliana_OX=3702_GN=GAMMA-ADR_PE=1_SV=1 |
| g196.t1 | sp_Q84K16_AP1G1_ARATH_AP-1_complex_subunit_gamma-1_OS=Arabidopsis_thaliana_OX=3702_GN=GAMMA-ADR_PE=1_SV=1 |
| g476.t1 | sp_P17847_NIR_MAIZE_Ferredoxin--nitrite_reductase,_chloroplastic_(Fragment)_OS=Zea_mays_OX=4577_GN=NIR_PE=2_SV=1 |
| g503.t1 | sp_Q9M160_Y4095_ARATH_Uncharacterized_protein_At4g00950_OS=Arabidopsis_thaliana_OX=3702_GN=At4g00950_PE=2_SV=1 |
| g877.t1 | sp_P35134_UBC11_ARATH_Ubiquitin-conjugating_enzyme_E2_11_OS=Arabidopsis_thaliana_OX=3702_GN=UBC11_PE=1_SV=2 |
| g1266.t1 | sp_Q6ESR4_DHN1_ORYSJ_Dehydrin_DHN1_OS=Oryza_sativa_subsp._japonica_OX=39947_GN=DHN1_PE=2_SV=1 |
| g1266.t1 | sp_Q6ESR4_DHN1_ORYSJ_Dehydrin_DHN1_OS=Oryza_sativa_subsp._japonica_OX=39947_GN=DHN1_PE=2_SV=1 |
| g1667.t1 | sp_O24575_DCAM_MAIZE_S-adenosylmethionine_decarboxylase_proenzyme_OS=Zea_mays_OX=4577_GN=SAMDC_PE=2_SV=1 |
| g1866.t1 | sp_O04331_PHB3_ARATH_Prohibitin-3,_mitochondrial_OS=Arabidopsis_thaliana_OX=3702_GN=PHB3_PE=1_SV=1 |
| g1897.t1 | sp_Q9SZ67_WRK19_ARATH_Probable_WRKY_transcription_factor_19_OS=Arabidopsis_thaliana_OX=3702_GN=WRKY19_PE=3_SV=1 |
| g1897.t1 | sp_Q9SZ67_WRK19_ARATH_Probable_WRKY_transcription_factor_19_OS=Arabidopsis_thaliana_OX=3702_GN=WRKY19_PE=3_SV=1 |
| g1897.t1 | sp_Q9SZ67_WRK19_ARATH_Probable_WRKY_transcription_factor_19_OS=Arabidopsis_thaliana_OX=3702_GN=WRKY19_PE=3_SV=1 |
| g1956.t1 | sp_A2X674_HOX7_ORYSI_Homeobox-leucine_zipper_protein_HOX7_OS=Oryza_sativa_subsp._indica_OX=39946_GN=HOX7_PE=1_SV=2 |
| g1956.t2 | sp_A2X674_HOX7_ORYSI_Homeobox-leucine_zipper_protein_HOX7_OS=Oryza_sativa_subsp._indica_OX=39946_GN=HOX7_PE=1_SV=2 |
| g1993.t1 | sp_O81815_MO1_ARATH_Monooxygenase_1_OS=Arabidopsis_thaliana_OX=3702_GN=MO1_PE=2_SV=1 |
| g2176.t1 | sp_P49174_INVA_MAIZE_Beta-fructofuranosidase,_cell_wall_isozyme_OS=Zea_mays_OX=4577_PE=2_SV=1 |
| g2252.t1 | sp_Q9LYU3_EF113_ARATH_Ethylene-responsive_transcription_factor_ERF113_OS=Arabidopsis_thaliana_OX=3702_GN=ERF113_PE=2_SV=1 |
| g2449.t1 | sp_F4KAF2_MORC4_ARATH_Protein_MICRORCHIDIA_4_OS=Arabidopsis_thaliana_OX=3702_GN=MORC4_PE=1_SV=2 |
| g2810.t1 | sp_Q9LV40_ROGF8_ARATH_Rho_guanine_nucleotide_exchange_factor_8_OS=Arabidopsis_thaliana_OX=3702_GN=ROPGEF8_PE=1_SV=1 |
| g2910.t1 | sp_Q9C3Z5_RLA2_PODAS_60S_acidic_ribosomal_protein_P2_OS=Podospora_anserina_OX=5145_PE=3_SV=1 |
| g2914.t1 | sp_Q41060_SBP65_PEA_Seed_biotin-containing_protein_SBP65_OS=Pisum_sativum_OX=3888_GN=SBP65_PE=1_SV=1 |
| g3070.t1 | sp_Q67YC0_PPSP1_ARATH_Inorganic_pyrophosphatase_1_OS=Arabidopsis_thaliana_OX=3702_GN=PS2_PE=1_SV=1 |
| g3392.t1 | sp_Q5UNY4_YL728_MIMIV_Uncharacterized_protein_L728_OS=Acanthamoeba_polyphaga_mimivirus_OX=212035_GN=MIMI_L728_PE=4_SV=1 |
| g3394.t1 | sp_Q6ETK9_ADPO2_ORYSJ_Heptahelical_transmembrane_protein_ADIPOR2_OS=Oryza_sativa_subsp._japonica_OX=39947_GN=ADIPOR2_PE=2_SV=1 |
| g3401.t1 | sp_Q6ETL8_GLO12_ORYSJ_Very-long-chain_aldehyde_decarbonylase_GL1-2_OS=Oryza_sativa_subsp._japonica_OX=39947_GN=GL1-2_PE=2_SV=1 |
| g3794.t1 | sp_Q29VN2_TPS2_MAIZE_Terpene_synthase_2,_chloroplastic_OS=Zea_mays_OX=4577_GN=TPS2_PE=1_SV=1 |
| g4028.t1 | sp_P46518_LEA14_GOSHI_Late_embryogenesis_abundant_protein_Lea14-A_OS=Gossypium_hirsutum_OX=3635_GN=LEA14-A_PE=2_SV=1 |
| g4037.t1 | sp_Q9CA93_BAC2_ARATH_Mitochondrial_arginine_transporter_BAC2_OS=Arabidopsis_thaliana_OX=3702_GN=BAC2_PE=1_SV=1 |
| g4073.t1 | sp_Q6I581_GH35_ORYSJ_Jasmonic_acid-amido_synthetase_JAR1_OS=Oryza_sativa_subsp._japonica_OX=39947_GN=GH3.5_PE=2_SV=1 |
| g4147.t1 | sp_Q1T7C2_C7111_SOLLC_Cytochrome_P450_710A11_OS=Solanum_lycopersicum_OX=4081_GN=CYP710A11_PE=1_SV=1 |
| g4147.t2 | sp_Q1T7C2_C7111_SOLLC_Cytochrome_P450_710A11_OS=Solanum_lycopersicum_OX=4081_GN=CYP710A11_PE=1_SV=1 |
| g4338.t2 | sp_Q4KLH6_CE162_RAT_Centrosomal_protein_of_162_kDa_OS=Rattus_norvegicus_OX=10116_GN=Cep162_PE=1_SV=2 |
| g4465.t1 | sp_Q5VQG4_RFS_ORYSJ_Galactinol--sucrose_galactosyltransferase_OS=Oryza_sativa_subsp._japonica_OX=39947_GN=RFS_PE=1_SV=1 |
| g4537.t1 | sp_O81865_P2A01_ARATH_Protein_PHLOEM_PROTEIN_2-LIKE_A1_OS=Arabidopsis_thaliana_OX=3702_GN=PP2A1_PE=2_SV=1 |
| g4590.t1 | sp_Q9GZZ6_ACH10_HUMAN_Neuronal_acetylcholine_receptor_subunit_alpha-10_OS=Homo_sapiens_OX=9606_GN=CHRNA10_PE=1_SV=1 |
| g4649.t1 | sp_Q10PI6_LCAT1_ORYSJ_Lecithin-cholesterol_acyltransferase-like_1_OS=Oryza_sativa_subsp._japonica_OX=39947_GN=Os03g0232800_PE=2_SV=1 |
| g4716.t1 | sp_P27777_HS16A_ORYSJ_16.9_kDa_class_I_heat_shock_protein_1_OS=Oryza_sativa_subsp._japonica_OX=39947_GN=HSP16.9A_PE=1_SV=1 |
| g4791.t1 | sp_Q5ZEG0_MHZ4_ORYSJ_Protein_MAO_HUZI_4,_chloroplastic_OS=Oryza_sativa_subsp._japonica_OX=39947_GN=MHZ4_PE=2_SV=1 |
| g4819.t1 | sp_K4BNG7_NAP2_SOLLC_NAC_domain-containing_protein_2_OS=Solanum_lycopersicum_OX=4081_GN=NAP2_PE=2_SV=1 |
| g4835.t1 | sp_Q489A2_RL24_COLP3_50S_ribosomal_protein_L24_OS=Colwellia_psychrerythraea_(strain_34H_/_ATCC_BAA-681)_OX=167879_GN=rplX_PE=3_SV=1 |
| g4919.t1 | sp_Q5Z8T8_XYXT1_ORYSJ_Beta-1,2-xylosyltransferase_XYXT1_OS=Oryza_sativa_subsp._japonica_OX=39947_GN=XYXT1_PE=1_SV=1 |
| g5079.t1 | sp_O06837_STYD_PSEFL_Phenylacetaldehyde_dehydrogenase_OS=Pseudomonas_fluorescens_OX=294_GN=styD_PE=1_SV=1 |
| g5158.t1 | sp_Q9SGN7_CCR12_ARATH_Serine/threonine-protein_kinase-like_protein_At1g28390_OS=Arabidopsis_thaliana_OX=3702_GN=At1g28390_PE=2_SV=1 |
| g5158.t1 | sp_Q9SGN7_CCR12_ARATH_Serine/threonine-protein_kinase-like_protein_At1g28390_OS=Arabidopsis_thaliana_OX=3702_GN=At1g28390_PE=2_SV=1 |
| g5223.t1 | sp_P20075_LEAD8_DAUCA_Embryonic_protein_DC-8_OS=Daucus_carota_OX=4039_PE=3_SV=1 |
| g5253.t1 | sp_Q9M060_IF62_ARATH_Eukaryotic_translation_initiation_factor_6-2_OS=Arabidopsis_thaliana_OX=3702_GN=EIF6-2_PE=2_SV=1 |
| g5396.t1 | sp_Q84JC2_DOGL4_ARATH_Protein_DOG1-like_4_OS=Arabidopsis_thaliana_OX=3702_GN=DOGL4_PE=2_SV=1 |
| g5458.t1 | sp_Q9SJA7_SOX_ARATH_Probable_sarcosine_oxidase_OS=Arabidopsis_thaliana_OX=3702_GN=At2g24580_PE=2_SV=1 |
| g5459.t1 | sp_Q9SJA7_SOX_ARATH_Probable_sarcosine_oxidase_OS=Arabidopsis_thaliana_OX=3702_GN=At2g24580_PE=2_SV=1 |
| g5615.t1 | sp_P46420_GSTF4_MAIZE_Glutathione_S-transferase_4_OS=Zea_mays_OX=4577_GN=GST4_PE=1_SV=2 |
| g5819.t1 | sp_Q6YRM6_Y8219_ORYSJ_Glycosyltransferase_family_92_protein_Os08g0121900_OS=Oryza_sativa_subsp._japonica_OX=39947_GN=Os08g0121900_PE=2_SV=1 |
| g5830.t1 | sp_Q42993_CHI1_ORYSJ_Chitinase_1_OS=Oryza_sativa_subsp._japonica_OX=39947_GN=Cht1_PE=2_SV=1 |
| g6281.t1 | sp_P48979_PGLR_PRUPE_Polygalacturonase_OS=Prunus_persica_OX=3760_PE=2_SV=1 |
| g6513.t1 | sp_Q9ZUX1_C94C1_ARATH_Cytochrome_P450_94C1_OS=Arabidopsis_thaliana_OX=3702_GN=CYP94C1_PE=1_SV=1 |
| g6538.t1 | sp_Q6WNQ8_C81E8_MEDTR_Cytochrome_P450_81E8_OS=Medicago_truncatula_OX=3880_GN=CYP81E8_PE=2_SV=1 |
| g6619.t1 | sp_Q941T1_P5CS2_ORYSJ_Delta-1-pyrroline-5-carboxylate_synthase_2_OS=Oryza_sativa_subsp._japonica_OX=39947_GN=P5CS2_PE=2_SV=1 |
| g6619.t1 | sp_Q941T1_P5CS2_ORYSJ_Delta-1-pyrroline-5-carboxylate_synthase_2_OS=Oryza_sativa_subsp._japonica_OX=39947_GN=P5CS2_PE=2_SV=1 |
| g6621.t1 | sp_P81713_IBB3_WHEAT_Bowman-Birk_type_trypsin_inhibitor_OS=Triticum_aestivum_OX=4565_PE=1_SV=1 |
| g6665.t1 | sp_P48495_TPIS_PETHY_Triosephosphate_isomerase,_cytosolic_OS=Petunia_hybrida_OX=4102_GN=TPIP1_PE=2_SV=1 |
| g6690.t1 | sp_Q9LX85_ZAT8_ARATH_Zinc_finger_protein_ZAT8_OS=Arabidopsis_thaliana_OX=3702_GN=ZAT8_PE=2_SV=1 |
| g6953.t1 | sp_Q93ZT5_EDL3_ARATH_EID1-like_F-box_protein_3_OS=Arabidopsis_thaliana_OX=3702_GN=EDL3_PE=2_SV=1 |
| g6959.t1 | sp_Q8S2G5_GPDH2_ORYSJ_Probable_glycerol-3-phosphate_dehydrogenase_[NAD(+)]_2,_cytosolic_OS=Oryza_sativa_subsp._japonica_OX=39947_GN=Os01g0801600_PE=2_SV=1 |
| g6959.t1 | sp_Q8S2G5_GPDH2_ORYSJ_Probable_glycerol-3-phosphate_dehydrogenase_[NAD(+)]_2,_cytosolic_OS=Oryza_sativa_subsp._japonica_OX=39947_GN=Os01g0801600_PE=2_SV=1 |
| g7002.t1 | sp_Q9SZY3_SBT38_ARATH_Subtilisin-like_protease_SBT3.8_OS=Arabidopsis_thaliana_OX=3702_GN=SBT3.8_PE=3_SV=1 |
| g7005.t1 | sp_Q0JIL1_NRX2_ORYSJ_Probable_nucleoredoxin_2_OS=Oryza_sativa_subsp._japonica_OX=39947_GN=Os01g0794400_PE=2_SV=1 |
| g7028.t1 | sp_Q9C7F5_NTF2B_ARATH_Nuclear_transport_factor_2B_OS=Arabidopsis_thaliana_OX=3702_GN=NTF2B_PE=1_SV=1 |
| g7389.t1 | sp_P51615_MAOX_VITVI_NADP-dependent_malic_enzyme_OS=Vitis_vinifera_OX=29760_PE=2_SV=1 |
| g7528.t1 | sp_O14354_MG101_SCHPO_Mitochondrial_genome_maintenance_protein_mgm101_OS=Schizosaccharomyces_pombe_(strain_972_/_ATCC_24843)_OX=284812_GN=mgm101_PE=3_SV=2 |
| g7530.t1 | sp_A1S8H7_Y2480_SHEAM_UPF0235_protein_Sama_2480_OS=Shewanella_amazonensis_(strain_ATCC_BAA-1098_/_SB2B)_OX=326297_GN=Sama_2480_PE=3_SV=1 |
| g7553.t1 | sp_Q9SV13_SUT31_ARATH_Sulfate_transporter_3.1_OS=Arabidopsis_thaliana_OX=3702_GN=SULTR3;1_PE=2_SV=1 |
| g7576.t1 | sp_Q55EX9_Y8948_DICDI_Putative_methyltransferase_DDB_G0268948_OS=Dictyostelium_discoideum_OX=44689_GN=DDB_G0268948_PE=1_SV=2 |
| g7643.t1 | sp_Q5N8F2_ILL2_ORYSJ_IAA-amino_acid_hydrolase_ILR1-like_2_OS=Oryza_sativa_subsp._japonica_OX=39947_GN=ILL2_PE=2_SV=1 |
| g7695.t1 | sp_Q6L4D2_PM19L_ORYSJ_Membrane_protein_PM19L_OS=Oryza_sativa_subsp._japonica_OX=39947_GN=PM19L_PE=2_SV=1 |
| g7716.t1 | sp_Q9SYI3_AB5B_ARATH_ABC_transporter_B_family_member_5_OS=Arabidopsis_thaliana_OX=3702_GN=ABCB5_PE=3_SV=1 |
| g7716.t1 | sp_Q9SYI3_AB5B_ARATH_ABC_transporter_B_family_member_5_OS=Arabidopsis_thaliana_OX=3702_GN=ABCB5_PE=3_SV=1 |
| g7729.t1 | sp_Q74KA5_SECA1_LACJO_Protein_translocase_subunit_SecA_1_OS=Lactobacillus_johnsonii_(strain_CNCM_I-12250_/_La1_/_NCC_533)_OX=257314_GN=secA1_PE=3_SV=1 |
| g7748.t1 | sp_Q06398_GSTU6_ORYSJ_Probable_glutathione_S-transferase_GSTU6_OS=Oryza_sativa_subsp._japonica_OX=39947_GN=GSTU6_PE=2_SV=2 |
| g7995.t1 | sp_P42776_GBF3_ARATH_G-box-binding_factor_3_OS=Arabidopsis_thaliana_OX=3702_GN=GBF3_PE=1_SV=2 |
| g8012.t1 | sp_Q8H0Y8_WRK41_ARATH_Probable_WRKY_transcription_factor_41_OS=Arabidopsis_thaliana_OX=3702_GN=WRKY41_PE=2_SV=2 |
| g8153.t1 | sp_Q8LBB2_KING1_ARATH_SNF1-related_protein_kinase_regulatory_subunit_gamma-1_OS=Arabidopsis_thaliana_OX=3702_GN=KING1_PE=1_SV=2 |
| g8188.t1 | sp_Q9LUC5_C7A15_ARATH_Cytochrome_P450_72A15_OS=Arabidopsis_thaliana_OX=3702_GN=CYP72A15_PE=2_SV=1 |
| g8224.t1 | sp_Q9FLV9_SLAH3_ARATH_S-type_anion_channel_SLAH3_OS=Arabidopsis_thaliana_OX=3702_GN=SLAH3_PE=1_SV=1 |
| g8282.t2 | sp_Q6P3W7_SCYL2_HUMAN_SCY1-like_protein_2_OS=Homo_sapiens_OX=9606_GN=SCYL2_PE=1_SV=1 |
| g8332.t1 | sp_B9G300_AB52G_ORYSJ_ABC_transporter_G_family_member_52_OS=Oryza_sativa_subsp._japonica_OX=39947_GN=ABCG52_PE=2_SV=2 |
| g8360.t1 | sp_A2WSD3_SWT6B_ORYSI_Bidirectional_sugar_transporter_SWEET6b_OS=Oryza_sativa_subsp._indica_OX=39946_GN=SWEET6B_PE=3_SV=1 |
| g8369.t1 | sp_Q5ZD81_CML12_ORYSJ_Probable_calcium-binding_protein_CML12_OS=Oryza_sativa_subsp._japonica_OX=39947_GN=CML12_PE=2_SV=1 |
| g8369.t1 | sp_Q5ZD81_CML12_ORYSJ_Probable_calcium-binding_protein_CML12_OS=Oryza_sativa_subsp._japonica_OX=39947_GN=CML12_PE=2_SV=1 |
| g8389.t1 | sp_Q9SU92_PPSP3_ARATH_Thiamine_phosphate_phosphatase-like_protein_OS=Arabidopsis_thaliana_OX=3702_GN=At4g29530_PE=1_SV=1 |
| g8409.t1 | sp_Q9LXF8_AVT1J_ARATH_Amino_acid_transporter_AVT1J_OS=Arabidopsis_thaliana_OX=3702_GN=AVT1J_PE=2_SV=1 |
| g8416.t1 | sp_Q9FJU3_FBD28_ARATH_Putative_FBD-associated_F-box_protein_At5g56690_OS=Arabidopsis_thaliana_OX=3702_GN=At5g56690_PE=4_SV=1 |
| g8481.t1 | sp_Q9C9F0_WRKY9_ARATH_Probable_WRKY_transcription_factor_9_OS=Arabidopsis_thaliana_OX=3702_GN=WRKY9_PE=2_SV=1 |
| g8601.t1 | sp_Q9M9Q6_SCP50_ARATH_Serine_carboxypeptidase-like_50_OS=Arabidopsis_thaliana_OX=3702_GN=SCPL50_PE=2_SV=1 |
| g8860.t2 | sp_Q652Q8_URH2_ORYSJ_Probable_uridine_nucleosidase_2_OS=Oryza_sativa_subsp._japonica_OX=39947_GN=URH2_PE=2_SV=1 |
| g8874.t1 | sp_O22527_CLH1_ARATH_Chlorophyllase-1_OS=Arabidopsis_thaliana_OX=3702_GN=CLH1_PE=1_SV=1 |
| g9248.t1 | sp_Q0J2L7_P2C68_ORYSJ_Probable_protein_phosphatase_2C_68_OS=Oryza_sativa_subsp._japonica_OX=39947_GN=Os09g0325700_PE=2_SV=2 |
| g9483.t1 | sp_Q4UL64_PRIA_RICFE_Primosomal_protein_N'_OS=Rickettsia_felis_(strain_ATCC_VR-1525_/_URRWXCal2)_OX=315456_GN=priA_PE=3_SV=1 |
| g9521.t1 | sp_Q9FE20_PBS1_ARATH_Serine/threonine-protein_kinase_PBS1_OS=Arabidopsis_thaliana_OX=3702_GN=PBS1_PE=1_SV=1 |
| g9521.t1 | sp_Q9FE20_PBS1_ARATH_Serine/threonine-protein_kinase_PBS1_OS=Arabidopsis_thaliana_OX=3702_GN=PBS1_PE=1_SV=1 |
| g9736.t1 | sp_Q8W453_DIRL1_ARATH_Putative_lipid-transfer_protein_DIR1_OS=Arabidopsis_thaliana_OX=3702_GN=DIR1_PE=1_SV=1 |
| g9820.t1 | sp_Q69T31_CINV1_ORYSJ_Cytosolic_invertase_1_OS=Oryza_sativa_subsp._japonica_OX=39947_GN=CINV1_PE=1_SV=1 |
| g9961.t1 | sp_Q7XV13_LOX5_ORYSJ_Putative_lipoxygenase_5_OS=Oryza_sativa_subsp._japonica_OX=39947_GN=Os04g0447100_PE=3_SV=2 |
| g10072.t1 | sp_F4J1Q9_AVT1I_ARATH_Amino_acid_transporter_AVT1I_OS=Arabidopsis_thaliana_OX=3702_GN=AVT1I_PE=3_SV=1 |
| g10222.t1 | sp_Q9FGS8_C2GR2_ARATH_C2_and_GRAM_domain-containing_protein_At5g50170_OS=Arabidopsis_thaliana_OX=3702_GN=At5g50170_PE=2_SV=1 |
| g10222.t1 | sp_Q9FGS8_C2GR2_ARATH_C2_and_GRAM_domain-containing_protein_At5g50170_OS=Arabidopsis_thaliana_OX=3702_GN=At5g50170_PE=2_SV=1 |
| g10332.t1 | sp_G5E8K5_ANK3_MOUSE_Ankyrin-3_OS=Mus_musculus_OX=10090_GN=Ank3_PE=1_SV=1 |
| g10332.t1 | sp_G5E8K5_ANK3_MOUSE_Ankyrin-3_OS=Mus_musculus_OX=10090_GN=Ank3_PE=1_SV=1 |
| g10332.t1 | sp_G5E8K5_ANK3_MOUSE_Ankyrin-3_OS=Mus_musculus_OX=10090_GN=Ank3_PE=1_SV=1 |
| g10332.t1 | sp_G5E8K5_ANK3_MOUSE_Ankyrin-3_OS=Mus_musculus_OX=10090_GN=Ank3_PE=1_SV=1 |
| g10332.t1 | sp_G5E8K5_ANK3_MOUSE_Ankyrin-3_OS=Mus_musculus_OX=10090_GN=Ank3_PE=1_SV=1 |
| g10332.t1 | sp_G5E8K5_ANK3_MOUSE_Ankyrin-3_OS=Mus_musculus_OX=10090_GN=Ank3_PE=1_SV=1 |
| g10332.t1 | sp_G5E8K5_ANK3_MOUSE_Ankyrin-3_OS=Mus_musculus_OX=10090_GN=Ank3_PE=1_SV=1 |
| g10363.t1 | sp_P29022_CHIA_MAIZE_Endochitinase_A_OS=Zea_mays_OX=4577_GN=CHIA_PE=1_SV=1 |
| g10726.t1 | sp_P49175_INV1_MAIZE_Beta-fructofuranosidase_1_OS=Zea_mays_OX=4577_GN=IVR1_PE=3_SV=1 |
| g10789.t1 | sp_Q33E23_DHE2_ORYSJ_Glutamate_dehydrogenase_2,_mitochondrial_OS=Oryza_sativa_subsp._japonica_OX=39947_GN=GDH2_PE=2_SV=1 |
| g10822.t1 | sp_Q8L7W2_NUDT8_ARATH_Nudix_hydrolase_8_OS=Arabidopsis_thaliana_OX=3702_GN=NUDT8_PE=2_SV=2 |
| g10971.t1 | sp_Q9Y2K9_STB5L_HUMAN_Syntaxin-binding_protein_5-like_OS=Homo_sapiens_OX=9606_GN=STXBP5L_PE=1_SV=2 |
| g11188.t1 | sp_Q96273_LEA18_ARATH_Late_embryogenesis_abundant_protein_18_OS=Arabidopsis_thaliana_OX=3702_GN=LEA18_PE=2_SV=1 |
| g11272.t1 | sp_O82807_AOX1A_ORYSJ_Ubiquinol_oxidase_1a,_mitochondrial_OS=Oryza_sativa_subsp._japonica_OX=39947_GN=AOX1A_PE=2_SV=1 |
| g11275.t1 | sp_O82766_AOX1B_ORYSJ_Ubiquinol_oxidase_1b,_mitochondrial_OS=Oryza_sativa_subsp._japonica_OX=39947_GN=AOX1B_PE=2_SV=1 |
| g11343.t1 | sp_A8SE82_RPOC1_CERDE_DNA-directed_RNA_polymerase_subunit_beta'_OS=Ceratophyllum_demersum_OX=4428_GN=rpoC1_PE=3_SV=1 |
| g11481.t1 | sp_Q7FAS1_GLO3_ORYSJ_Peroxisomal_(S)-2-hydroxy-acid_oxidase_GLO3_OS=Oryza_sativa_subsp._japonica_OX=39947_GN=GLO3_PE=2_SV=1 |
| g11596.t1 | sp_Q7X7H9_SK3_ORYSJ_Shikimate_kinase_3,_chloroplastic_OS=Oryza_sativa_subsp._japonica_OX=39947_GN=SK3_PE=1_SV=2 |
| g11705.t1 | sp_Q9C9G4_ENDO2_ARATH_Endonuclease_2_OS=Arabidopsis_thaliana_OX=3702_GN=ENDO2_PE=1_SV=1 |
| g11981.t2 | sp_Q9M022_AIRP2_ARATH_E3_ubiquitin-protein_ligase_AIRP2_OS=Arabidopsis_thaliana_OX=3702_GN=AIRP2_PE=1_SV=1 |
| g12043.t1 | sp_Q96520_PER12_ARATH_Peroxidase_12_OS=Arabidopsis_thaliana_OX=3702_GN=PER12_PE=1_SV=1 |
| g12159.t1 | sp_Q7WZE5_KGUA_SHEVD_Guanylate_kinase_OS=Shewanella_violacea_(strain_JCM_10179_/_CIP_106290_/_LMG_19151_/_DSS12)_OX=637905_GN=gmk_PE=3_SV=1 |
| g12235.t1 | sp_P12950_DHN1_MAIZE_Dehydrin_DHN1_OS=Zea_mays_OX=4577_GN=DHN1_PE=1_SV=2 |
| g12304.t1 | sp_Q2R4Z4_DHR21_ORYSJ_Water_stress-inducible_protein_Rab21_OS=Oryza_sativa_subsp._japonica_OX=39947_GN=RAB21_PE=2_SV=1 |
| g12358.t1 | sp_C5Y376_CSPLB_SORBI_CASP-like_protein_1U2_OS=Sorghum_bicolor_OX=4558_GN=Sb05g019440_PE=2_SV=1 |
| g12433.t1 | sp_Q9ZSA7_DLO2_ARATH_Protein_DMR6-LIKE_OXYGENASE_2_OS=Arabidopsis_thaliana_OX=3702_GN=DLO2_PE=2_SV=1 |
| g12737.t1 | sp_Q9SVG5_BBE18_ARATH_Berberine_bridge_enzyme-like_18_OS=Arabidopsis_thaliana_OX=3702_GN=At4g20820_PE=3_SV=1 |
| g13374.t1 | sp_Q5SMM8_PHT1_ORYSJ_Putrescine_hydroxycinnamoyltransferase_1_OS=Oryza_sativa_subsp._japonica_OX=39947_GN=PHT1_PE=2_SV=1 |
| g13420.t1 | sp_P42813_RNS1_ARATH_Ribonuclease_1_OS=Arabidopsis_thaliana_OX=3702_GN=RNS1_PE=1_SV=1 |
| g13689.t1 | sp_F4K956_A70_ARATH_Pathogen-associated_molecular_patterns-induced_protein_A70_OS=Arabidopsis_thaliana_OX=3702_GN=A70_PE=1_SV=1 |
| g13689.t1 | sp_F4K956_A70_ARATH_Pathogen-associated_molecular_patterns-induced_protein_A70_OS=Arabidopsis_thaliana_OX=3702_GN=A70_PE=1_SV=1 |
| g13994.t1 | sp_O22527_CLH1_ARATH_Chlorophyllase-1_OS=Arabidopsis_thaliana_OX=3702_GN=CLH1_PE=1_SV=1 |
| g14001.t1 | sp_Q2QSL4_CDKF2_ORYSJ_Putative_cyclin-dependent_kinase_F-2_OS=Oryza_sativa_subsp._japonica_OX=39947_GN=CDKF-2_PE=3_SV=1 |
| g14091.t1 | sp_P58293_Y3446_CLOAB_Uncharacterized_isomerase_CA_C3446_OS=Clostridium_acetobutylicum_(strain_ATCC_824_/_DSM_792_/_JCM_1419_/_LMG_5710_/_VKM_B-1787)_OX=272562_GN=CA_C3446_PE=3_SV=1 |
| g14135.t1 | sp_Q94A08_RFS2_ARATH_Probable_galactinol--sucrose_galactosyltransferase_2_OS=Arabidopsis_thaliana_OX=3702_GN=RFS2_PE=2_SV=2 |
| g14378.t1 | sp_Q01417_PM1_SOYBN_18_kDa_seed_maturation_protein_OS=Glycine_max_OX=3847_GN=GMPM1_PE=2_SV=1 |
| g14387.t1 | sp_Q8VZ80_PLT5_ARATH_Polyol_transporter_5_OS=Arabidopsis_thaliana_OX=3702_GN=PLT5_PE=1_SV=2 |
| g14387.t1 | sp_Q8VZ80_PLT5_ARATH_Polyol_transporter_5_OS=Arabidopsis_thaliana_OX=3702_GN=PLT5_PE=1_SV=2 |
| g14454.t1 | sp_Q8S9J6_ASPA_ARATH_Aspartyl_protease_family_protein_At5g10770_OS=Arabidopsis_thaliana_OX=3702_GN=At5g10770_PE=2_SV=1 |
| g14553.t1 | sp_Q9XIH6_PLT2_ARATH_Putative_polyol_transporter_2_OS=Arabidopsis_thaliana_OX=3702_GN=PLT2_PE=3_SV=1 |
| g14597.t1 | sp_P0DKC6_HSD1B_ARATH_11-beta-hydroxysteroid_dehydrogenase_1B_OS=Arabidopsis_thaliana_OX=3702_GN=HSD1_PE=1_SV=1 |
| g14781.t1 | sp_P23673_CTFB_CLOAB_Butyrate--acetoacetate_CoA-transferase_subunit_B_OS=Clostridium_acetobutylicum_(strain_ATCC_824_/_DSM_792_/_JCM_1419_/_LMG_5710_/_VKM_B-1787)_OX=272562_GN=ctfB_PE=1_SV=1 |
| g14990.t1 | sp_P31414_AVP1_ARATH_Pyrophosphate-energized_vacuolar_membrane_proton_pump_1_OS=Arabidopsis_thaliana_OX=3702_GN=AVP1_PE=1_SV=1 |
| g15029.t1 | sp_P93438_METK2_ORYSJ_S-adenosylmethionine_synthase_2_OS=Oryza_sativa_subsp._japonica_OX=39947_GN=SAM2_PE=2_SV=1 |
| g15050.t1 | sp_Q9SJA7_SOX_ARATH_Probable_sarcosine_oxidase_OS=Arabidopsis_thaliana_OX=3702_GN=At2g24580_PE=2_SV=1 |
| g15051.t1 | sp_Q9SJA7_SOX_ARATH_Probable_sarcosine_oxidase_OS=Arabidopsis_thaliana_OX=3702_GN=At2g24580_PE=2_SV=1 |
| g15119.t1 | sp_Q0WMZ5_OP162_ARATH_Outer_envelope_pore_protein_16-2,_chloroplastic_OS=Arabidopsis_thaliana_OX=3702_GN=OEP162_PE=2_SV=1 |
| g15275.t1 | sp_Q9ZP27_AVCO2_AVESA_Avenacosidase_2_OS=Avena_sativa_OX=4498_GN=P60B_PE=1_SV=1 |
| g15481.t1 | sp_Q9SLN8_DBR_TOBAC_2-alkenal_reductase_(NADP(+)-dependent)_OS=Nicotiana_tabacum_OX=4097_GN=DBR_PE=1_SV=1 |
| g15657.t2 | sp_Q9V6X7_OFUT1_DROME_GDP-fucose_protein_O-fucosyltransferase_1_OS=Drosophila_melanogaster_OX=7227_GN=O-fut1_PE=1_SV=1 |
| g15672.t1 | sp_Q99090_CPRF2_PETCR_Light-inducible_protein_CPRF2_OS=Petroselinum_crispum_OX=4043_GN=CPRF2_PE=2_SV=2 |
| g15672.t2 | sp_Q99090_CPRF2_PETCR_Light-inducible_protein_CPRF2_OS=Petroselinum_crispum_OX=4043_GN=CPRF2_PE=2_SV=2 |
| g15774.t1 | sp_P0DKC6_HSD1B_ARATH_11-beta-hydroxysteroid_dehydrogenase_1B_OS=Arabidopsis_thaliana_OX=3702_GN=HSD1_PE=1_SV=1 |
| g15905.t2 | sp_Q42602_C89A2_ARATH_Cytochrome_P450_89A2_OS=Arabidopsis_thaliana_OX=3702_GN=CYP89A2_PE=2_SV=2 |
| g15905.t1 | sp_Q42602_C89A2_ARATH_Cytochrome_P450_89A2_OS=Arabidopsis_thaliana_OX=3702_GN=CYP89A2_PE=2_SV=2 |
| g15977.t1 | sp_Q04980_LTI65_ARATH_Low-temperature-induced_65_kDa_protein_OS=Arabidopsis_thaliana_OX=3702_GN=LTI65_PE=2_SV=2 |
| g16051.t1 | sp_Q9FT97_AGAL1_ARATH_Alpha-galactosidase_1_OS=Arabidopsis_thaliana_OX=3702_GN=AGAL1_PE=2_SV=1 |
| g16053.t1 | sp_Q9ATL7_TIP31_MAIZE_Aquaporin_TIP3-1_OS=Zea_mays_OX=4577_GN=TIP3-1_PE=2_SV=1 |
| g16058.t1 | sp_C0HJG8_BSP_BOSSE_Basic_secretory_protease_(Fragments)_OS=Boswellia_serrata_OX=613112_PE=1_SV=1 |
| g16101.t1 | sp_P96198_DHAS_AZOVI_Aspartate-semialdehyde_dehydrogenase_OS=Azotobacter_vinelandii_OX=354_GN=asd_PE=3_SV=1 |
| g16147.t1 | sp_Q9AV57_FLOT1_ORYSJ_Flotillin-like_protein_1_OS=Oryza_sativa_subsp._japonica_OX=39947_GN=FLOT1_PE=3_SV=1 |
| g16243.t1 | sp_B8BHF1_GLO15_ORYSI_Very-long-chain_aldehyde_decarbonylase_GL1-5_OS=Oryza_sativa_subsp._indica_OX=39946_GN=GL1-5_PE=2_SV=1 |
| g16545.t1 | sp_Q0V842_NEN2_ARATH_Protein_NEN2_OS=Arabidopsis_thaliana_OX=3702_GN=NEN2_PE=2_SV=1 |
| g16545.t1 | sp_Q0V842_NEN2_ARATH_Protein_NEN2_OS=Arabidopsis_thaliana_OX=3702_GN=NEN2_PE=2_SV=1 |
| g16750.t1 | sp_Q9ZPE7_EXO_ARATH_Protein_EXORDIUM_OS=Arabidopsis_thaliana_OX=3702_GN=EXO_PE=2_SV=1 |
| g16987.t1 | sp_P46032_PTR2_ARATH_Protein_NRT1/_PTR_FAMILY_8.3_OS=Arabidopsis_thaliana_OX=3702_GN=NPF8.3_PE=1_SV=1 |
| g17034.t2 | sp_Q9C7Z9_SCP18_ARATH_Serine_carboxypeptidase-like_18_OS=Arabidopsis_thaliana_OX=3702_GN=SCPL18_PE=2_SV=2 |
| g17074.t1 | sp_Q38909_XTH28_ARATH_Probable_xyloglucan_endotransglucosylase/hydrolase_protein_28_OS=Arabidopsis_thaliana_OX=3702_GN=XTH28_PE=2_SV=1 |
| g17169.t1 | sp_Q9ZVJ5_SGGP_ARATH_Haloacid_dehalogenase-like_hydrolase_domain-containing_protein_Sgpp_OS=Arabidopsis_thaliana_OX=3702_GN=SGPP_PE=1_SV=2 |
| g17312.t1 | sp_Q05085_PTR7_ARATH_Protein_NRT1/_PTR_FAMILY_6.3_OS=Arabidopsis_thaliana_OX=3702_GN=NPF6.3_PE=1_SV=1 |
| g17719.t1 | sp_Q7FAX1_PXG_ORYSJ_Peroxygenase_OS=Oryza_sativa_subsp._japonica_OX=39947_GN=PXG_PE=2_SV=1 |
| g18059.t1 | sp_Q9LRQ8_PMAT2_ARATH_Phenolic_glucoside_malonyltransferase_2_OS=Arabidopsis_thaliana_OX=3702_GN=PMAT2_PE=1_SV=1 |
| g18154.t1 | sp_P30986_RETO_ESCCA_Reticuline_oxidase_OS=Eschscholzia_californica_OX=3467_GN=BBE1_PE=1_SV=1 |
| g18194.t1 | sp_Q9SU40_SKU5_ARATH_Monocopper_oxidase-like_protein_SKU5_OS=Arabidopsis_thaliana_OX=3702_GN=SKU5_PE=1_SV=1 |
| g18278.t1 | sp_Q6ZJK7_TDC1_ORYSJ_Tryptophan_decarboxylase_1_OS=Oryza_sativa_subsp._japonica_OX=39947_GN=TDC1_PE=1_SV=1 |
| g18457.t1 | sp_Q43246_C88A1_MAIZE_Cytochrome_P450_88A1_OS=Zea_mays_OX=4577_GN=CYP88A1_PE=2_SV=1 |
| g18556.t1 | sp_Q2R4Z5_DH16B_ORYSJ_Dehydrin_Rab16B_OS=Oryza_sativa_subsp._japonica_OX=39947_GN=RAB16B_PE=2_SV=1 |
| g18610.t1 | sp_Q9M897_DMP5_ARATH_Protein_DMP5_OS=Arabidopsis_thaliana_OX=3702_GN=DMP5_PE=2_SV=1 |
| g18613.t1 | sp_P12950_DHN1_MAIZE_Dehydrin_DHN1_OS=Zea_mays_OX=4577_GN=DHN1_PE=1_SV=2 |
| g18663.t1 | sp_B8EF98_Y2728_SHEB2_UPF0319_protein_Sbal223_2728_OS=Shewanella_baltica_(strain_OS223)_OX=407976_GN=Sbal223_2728_PE=3_SV=1 |
| g18742.t2 | sp_Q56X72_RTNLS_ARATH_Reticulon-like_protein_B21_OS=Arabidopsis_thaliana_OX=3702_GN=RTNLB21_PE=2_SV=2 |
| g18900.t1 | sp_Q9SY29_PUP4_ARATH_Probable_purine_permease_4_OS=Arabidopsis_thaliana_OX=3702_GN=PUP4_PE=2_SV=1 |
| g18900.t1 | sp_Q9SY29_PUP4_ARATH_Probable_purine_permease_4_OS=Arabidopsis_thaliana_OX=3702_GN=PUP4_PE=2_SV=1 |
| g19023.t1 | sp_P22180_PMA1_SOLLC_Plasma_membrane_ATPase_1_OS=Solanum_lycopersicum_OX=4081_GN=LHA1_PE=2_SV=1 |
| g19158.t1 | sp_Q9NP78_ABCB9_HUMAN_ATP-binding_cassette_sub-family_B_member_9_OS=Homo_sapiens_OX=9606_GN=ABCB9_PE=1_SV=1 |
| g19194.t1 | sp_Q8L7U7_ACDH_ARATH_ACD11_homolog_protein_OS=Arabidopsis_thaliana_OX=3702_GN=At4g39670_PE=2_SV=1 |
| g19212.t1 | sp_P38560_GLNA2_MAIZE_Glutamine_synthetase_root_isozyme_2_OS=Zea_mays_OX=4577_GN=GLN2_PE=2_SV=1 |
| g19264.t1 | sp_O07427_MGLL_MYCTU_Monoacylglycerol_lipase_OS=Mycobacterium_tuberculosis_(strain_ATCC_25618_/_H37Rv)_OX=83332_GN=Rv0183_PE=1_SV=2 |
| g19284.t1 | sp_Q9M069_E137_ARATH_Glucan_endo-1,3-beta-glucosidase_7_OS=Arabidopsis_thaliana_OX=3702_GN=At4g34480_PE=1_SV=2 |
| g19335.t1 | sp_Q851F9_EGY3_ORYSJ_Probable_zinc_metalloprotease_EGY3,_chloroplastic_OS=Oryza_sativa_subsp._japonica_OX=39947_GN=EGY3_PE=2_SV=1 |
| g19489.t1 | sp_P24805_TSJT1_TOBAC_Stem-specific_protein_TSJT1_OS=Nicotiana_tabacum_OX=4097_GN=TSJT1_PE=2_SV=1 |
| g19709.t1 | sp_Q10EJ2_CSPLU_ORYSJ_CASP-like_protein_5C1_OS=Oryza_sativa_subsp._japonica_OX=39947_GN=Os03g0767900_PE=2_SV=1 |
| g20000.t1 | sp_Q10BU2_GL37_ORYSJ_Germin-like_protein_3-7_OS=Oryza_sativa_subsp._japonica_OX=39947_GN=GER7_PE=2_SV=1 |
| g20132.t1 | sp_Q84TB6_ADF3_ORYSJ_Actin-depolymerizing_factor_3_OS=Oryza_sativa_subsp._japonica_OX=39947_GN=ADF3_PE=1_SV=1 |
| g20198.t1 | sp_Q9LT10_CXE18_ARATH_Probable_carboxylesterase_18_OS=Arabidopsis_thaliana_OX=3702_GN=CXE18_PE=1_SV=1 |
| g20284.t1 | sp_Q7GCM7_XIP1_ORYSJ_Xylanase_inhibitor_protein_1_OS=Oryza_sativa_subsp._japonica_OX=39947_GN=RIXI_PE=1_SV=1 |
| g20323.t1 | sp_B9DGI8_BZP63_ARATH_Basic_leucine_zipper_63_OS=Arabidopsis_thaliana_OX=3702_GN=BZIP63_PE=1_SV=1 |
| g20350.t1 | sp_Q84MA8_AZG2_ARATH_Adenine/guanine_permease_AZG2_OS=Arabidopsis_thaliana_OX=3702_GN=AZG2_PE=2_SV=1 |
| g20444.t1 | sp_Q09893_YAI5_SCHPO_Uncharacterized_protein_C24B11.05_OS=Schizosaccharomyces_pombe_(strain_972_/_ATCC_24843)_OX=284812_GN=SPAC24B11.05_PE=3_SV=1 |
| g20484.t1 | sp_P37220_ASR3_SOLLC_Abscisic_stress-ripening_protein_3_OS=Solanum_lycopersicum_OX=4081_GN=ASR3_PE=3_SV=2 |
| g20527.t1 | sp_Q5Z8T8_XYXT1_ORYSJ_Beta-1,2-xylosyltransferase_XYXT1_OS=Oryza_sativa_subsp._japonica_OX=39947_GN=XYXT1_PE=1_SV=1 |
| g20560.t1 | sp_A2WZ60_IRO2_ORYSI_Protein_IRON-RELATED_TRANSCRIPTION_FACTOR_2_OS=Oryza_sativa_subsp._indica_OX=39946_GN=IRO2_PE=3_SV=1 |
| g20957.t1 | sp_Q9M390_PTR1_ARATH_Protein_NRT1/_PTR_FAMILY_8.1_OS=Arabidopsis_thaliana_OX=3702_GN=NPF8.1_PE=1_SV=1 |
| g21028.t1 | sp_P29250_LOX2_ORYSJ_Linoleate_9S-lipoxygenase_2_OS=Oryza_sativa_subsp._japonica_OX=39947_GN=LOX1.1_PE=2_SV=2 |
| g21195.t2 | sp_Q0WP01_PTR9_ARATH_Protein_NRT1/_PTR_FAMILY_5.10_OS=Arabidopsis_thaliana_OX=3702_GN=NPF5.10_PE=2_SV=1 |
| g21326.t1 | sp_Q6WNQ8_C81E8_MEDTR_Cytochrome_P450_81E8_OS=Medicago_truncatula_OX=3880_GN=CYP81E8_PE=2_SV=1 |
| g21406.t1 | sp_Q40193_RB11C_LOTJA_Ras-related_protein_Rab11C_OS=Lotus_japonicus_OX=34305_GN=RAB11C_PE=2_SV=1 |
| g21419.t1 | sp_Q0PGJ6_AKRC9_ARATH_NADPH-dependent_aldo-keto_reductase,_chloroplastic_OS=Arabidopsis_thaliana_OX=3702_GN=AKR4C9_PE=1_SV=1 |
| g21419.t1 | sp_Q0PGJ6_AKRC9_ARATH_NADPH-dependent_aldo-keto_reductase,_chloroplastic_OS=Arabidopsis_thaliana_OX=3702_GN=AKR4C9_PE=1_SV=1 |
| g21488.t1 | sp_Q9LX85_ZAT8_ARATH_Zinc_finger_protein_ZAT8_OS=Arabidopsis_thaliana_OX=3702_GN=ZAT8_PE=2_SV=1 |
| g21488.t1 | sp_Q9LX85_ZAT8_ARATH_Zinc_finger_protein_ZAT8_OS=Arabidopsis_thaliana_OX=3702_GN=ZAT8_PE=2_SV=1 |
| g21533.t1 | sp_Q9ZQC5_ICR2_ARATH_Interactor_of_constitutive_active_ROPs_2,_chloroplastic_OS=Arabidopsis_thaliana_OX=3702_GN=ICR2_PE=1_SV=1 |
| g21536.t1 | sp_Q3AUR5_CLPS_SYNS9_ATP-dependent_Clp_protease_adapter_protein_ClpS_OS=Synechococcus_sp._(strain_CC9902)_OX=316279_GN=clpS_PE=3_SV=1 |
| g21640.t1 | sp_Q42797_TCMO_SOYBN_Trans-cinnamate_4-monooxygenase_OS=Glycine_max_OX=3847_GN=CYP73A11_PE=2_SV=1 |
| g21640.t1 | sp_Q42797_TCMO_SOYBN_Trans-cinnamate_4-monooxygenase_OS=Glycine_max_OX=3847_GN=CYP73A11_PE=2_SV=1 |
| g21847.t1 | sp_Q9LXA5_LRK91_ARATH_L-type_lectin-domain_containing_receptor_kinase_IX.1_OS=Arabidopsis_thaliana_OX=3702_GN=LECRK91_PE=1_SV=1 |
| g21911.t1 | sp_Q69NX5_NCED4_ORYSJ_9-cis-epoxycarotenoid_dioxygenase_NCED4,_chloroplastic_OS=Oryza_sativa_subsp._japonica_OX=39947_GN=NCED4_PE=2_SV=1 |
| g21966.t1 | sp_B8XCH5_QKY_ARATH_Protein_QUIRKY_OS=Arabidopsis_thaliana_OX=3702_GN=QKY_PE=2_SV=1 |
| g21966.t1 | sp_B8XCH5_QKY_ARATH_Protein_QUIRKY_OS=Arabidopsis_thaliana_OX=3702_GN=QKY_PE=2_SV=1 |
| g21966.t1 | sp_B8XCH5_QKY_ARATH_Protein_QUIRKY_OS=Arabidopsis_thaliana_OX=3702_GN=QKY_PE=2_SV=1 |
| g22080.t1 | sp_Q8L5C6_XIP1_WHEAT_Xylanase_inhibitor_protein_1_OS=Triticum_aestivum_OX=4565_GN=XIPI_PE=1_SV=2 |
| g22220.t1 | sp_P31110_TLP_ORYSJ_Thaumatin-like_protein_OS=Oryza_sativa_subsp._japonica_OX=39947_GN=Os12g0628600_PE=1_SV=1 |
| g22260.t1 | sp_P82953_LECH_HORVV_Horcolin_OS=Hordeum_vulgare_subsp._vulgare_OX=112509_PE=1_SV=2 |
| g22603.t1 | sp_Q851F9_EGY3_ORYSJ_Probable_zinc_metalloprotease_EGY3,_chloroplastic_OS=Oryza_sativa_subsp._japonica_OX=39947_GN=EGY3_PE=2_SV=1 |
| g22657.t1 | sp_P10765_PRL_LOXAF_Prolactin_OS=Loxodonta_africana_OX=9785_GN=PRL_PE=1_SV=1 |
| g22686.t1 | sp_O07427_MGLL_MYCTU_Monoacylglycerol_lipase_OS=Mycobacterium_tuberculosis_(strain_ATCC_25618_/_H37Rv)_OX=83332_GN=Rv0183_PE=1_SV=2 |
| g22695.t2 | sp_Q9LVJ0_UVB31_ARATH_UV-B-induced_protein_At3g17800,_chloroplastic_OS=Arabidopsis_thaliana_OX=3702_GN=At3g17800_PE=2_SV=1 |
| g22737.t1 | sp_P38560_GLNA2_MAIZE_Glutamine_synthetase_root_isozyme_2_OS=Zea_mays_OX=4577_GN=GLN2_PE=2_SV=1 |
| g22841.t1 | sp_Q6GMH0_PRP18_DANRE_Pre-mRNA-splicing_factor_18_OS=Danio_rerio_OX=7955_GN=prpf18_PE=2_SV=1 |
| g22919.t1 | sp_P22180_PMA1_SOLLC_Plasma_membrane_ATPase_1_OS=Solanum_lycopersicum_OX=4081_GN=LHA1_PE=2_SV=1 |
| g23165.t1 | sp_Q6Q1S1_NS3_CVHNL_Non-structural_protein_3_OS=Human_coronavirus_NL63_OX=277944_GN=3_PE=4_SV=1 |
| g23574.t1 | sp_Q9FXA3_BH095_ARATH_Transcription_factor_bHLH95_OS=Arabidopsis_thaliana_OX=3702_GN=BHLH95_PE=2_SV=2 |
| g23576.t1 | sp_Q9AV39_GLO1A_ORYSJ_Very-long-chain_aldehyde_decarbonylase_GL1-10_OS=Oryza_sativa_subsp._japonica_OX=39947_GN=GL1-10_PE=2_SV=1 |
| g23914.t1 | sp_Q69IL4_RF2A_ORYSJ_Transcription_factor_RF2a_OS=Oryza_sativa_subsp._japonica_OX=39947_GN=RF2a_PE=1_SV=1 |
| g24040.t1 | sp_Q10RZ1_BAMY2_ORYSJ_Beta-amylase_2,_chloroplastic_OS=Oryza_sativa_subsp._japonica_OX=39947_GN=BAMY2_PE=1_SV=1 |
| g24175.t2 | sp_P09444_LEA34_GOSHI_Late_embryogenesis_abundant_protein_D-34_OS=Gossypium_hirsutum_OX=3635_PE=3_SV=1 |
| g24175.t1 | sp_P09444_LEA34_GOSHI_Late_embryogenesis_abundant_protein_D-34_OS=Gossypium_hirsutum_OX=3635_PE=3_SV=1 |
| g24192.t1 | sp_Q8NF91_SYNE1_HUMAN_Nesprin-1_OS=Homo_sapiens_OX=9606_GN=SYNE1_PE=1_SV=4 |
| g24267.t1 | sp_P13940_LEA29_GOSHI_Late_embryogenesis_abundant_protein_D-29_OS=Gossypium_hirsutum_OX=3635_PE=3_SV=1 |
| g24591.t1 | sp_Q3ZBR5_TTC1_BOVIN_Tetratricopeptide_repeat_protein_1_OS=Bos_taurus_OX=9913_GN=TTC1_PE=2_SV=1 |
| g25221.t1 | sp_Q9ZNQ7_RCI2A_ARATH_Hydrophobic_protein_RCI2A_OS=Arabidopsis_thaliana_OX=3702_GN=RCI2A_PE=2_SV=1 |
| g25298.t1 | sp_Q9FLB1_PYL5_ARATH_Abscisic_acid_receptor_PYL5_OS=Arabidopsis_thaliana_OX=3702_GN=PYL5_PE=1_SV=1 |
| g25321.t1 | sp_Q05736_PR1_ASPOF_Pathogenesis-related_protein_1_OS=Asparagus_officinalis_OX=4686_GN=PR1_PE=2_SV=1 |
| g25322.t1 | sp_Q05736_PR1_ASPOF_Pathogenesis-related_protein_1_OS=Asparagus_officinalis_OX=4686_GN=PR1_PE=2_SV=1 |
| g25332.t1 | sp_Q740E5_PYRG_MYCPA_CTP_synthase_OS=Mycobacterium_paratuberculosis_(strain_ATCC_BAA-968_/_K-10)_OX=262316_GN=pyrG_PE=3_SV=1 |
| g25391.t1 | sp_Q0WMZ5_OP162_ARATH_Outer_envelope_pore_protein_16-2,_chloroplastic_OS=Arabidopsis_thaliana_OX=3702_GN=OEP162_PE=2_SV=1 |
| g25494.t1 | sp_Q2GWF3_SET1_CHAGB_Histone-lysine_N-methyltransferase,_H3_lysine-4_specific_OS=Chaetomium_globosum_(strain_ATCC_6205_/_CBS_148.51_/_DSM_1962_/_NBRC_6347_/_NRRL_1970)_OX=306901_GN=SET1_PE=3_SV=1 |
| g25529.t1 | sp_Q10M12_40C1_ORYSJ_Ricin_B-like_lectin_R40C1_OS=Oryza_sativa_subsp._japonica_OX=39947_GN=R40C1_PE=1_SV=1 |
| g25529.t1 | sp_Q10M12_40C1_ORYSJ_Ricin_B-like_lectin_R40C1_OS=Oryza_sativa_subsp._japonica_OX=39947_GN=R40C1_PE=1_SV=1 |
| g25559.t1 | sp_Q0ITS8_RL101_ORYSJ_60S_ribosomal_protein_L10-1_OS=Oryza_sativa_subsp._japonica_OX=39947_GN=SC34_PE=2_SV=1 |
| g25628.t1 | sp_Q10LN5_SWT16_ORYSJ_Bidirectional_sugar_transporter_SWEET16_OS=Oryza_sativa_subsp._japonica_OX=39947_GN=SWEET16_PE=3_SV=1 |
| g25635.t2 | sp_Q56S59_PHYLL_TOBAC_Phylloplanin_OS=Nicotiana_tabacum_OX=4097_PE=1_SV=1 |
| g25635.t1 | sp_Q56S59_PHYLL_TOBAC_Phylloplanin_OS=Nicotiana_tabacum_OX=4097_PE=1_SV=1 |
| g25851.t1 | sp_Q9Y7K5_YGI3_SCHPO_Uncharacterized_WD_repeat-containing_protein_C2A9.03_OS=Schizosaccharomyces_pombe_(strain_972_/_ATCC_24843)_OX=284812_GN=SPBC2A9.03_PE=4_SV=2 |
| g25873.t1 | sp_A4JPX5_MHPC_BURVG_2-hydroxy-6-oxononadienedioate/2-hydroxy-6-oxononatrienedioate_hydrolase_OS=Burkholderia_vietnamiensis_(strain_G4_/_LMG_22486)_OX=269482_GN=mhpC_PE=3_SV=1 |
| g25885.t1 | sp_Q75LR7_SAPK1_ORYSJ_Serine/threonine-protein_kinase_SAPK1_OS=Oryza_sativa_subsp._japonica_OX=39947_GN=SAPK1_PE=1_SV=1 |
| g25927.t1 | sp_Q9M088_E135_ARATH_Glucan_endo-1,3-beta-glucosidase_5_OS=Arabidopsis_thaliana_OX=3702_GN=At4g31140_PE=2_SV=1 |
| g26070.t1 | sp_Q9FLD5_ASD_ARATH_AAA-ATPase_ASD,_mitochondrial_OS=Arabidopsis_thaliana_OX=3702_GN=AATP1_PE=1_SV=1 |
| g26328.t1 | sp_Q7XCK6_CHI8_ORYSJ_Chitinase_8_OS=Oryza_sativa_subsp._japonica_OX=39947_GN=Cht8_PE=2_SV=1 |
| g26328.t2 | sp_Q7XCK6_CHI8_ORYSJ_Chitinase_8_OS=Oryza_sativa_subsp._japonica_OX=39947_GN=Cht8_PE=2_SV=1 |
| g26404.t1 | sp_Q05085_PTR7_ARATH_Protein_NRT1/_PTR_FAMILY_6.3_OS=Arabidopsis_thaliana_OX=3702_GN=NPF6.3_PE=1_SV=1 |
| g26404.t1 | sp_Q05085_PTR7_ARATH_Protein_NRT1/_PTR_FAMILY_6.3_OS=Arabidopsis_thaliana_OX=3702_GN=NPF6.3_PE=1_SV=1 |
| g26550.t1 | sp_Q9ZVJ5_SGGP_ARATH_Haloacid_dehalogenase-like_hydrolase_domain-containing_protein_Sgpp_OS=Arabidopsis_thaliana_OX=3702_GN=SGPP_PE=1_SV=2 |
| g26639.t1 | sp_Q38909_XTH28_ARATH_Probable_xyloglucan_endotransglucosylase/hydrolase_protein_28_OS=Arabidopsis_thaliana_OX=3702_GN=XTH28_PE=2_SV=1 |
| g26678.t1 | sp_Q8VZU3_SCP19_ARATH_Serine_carboxypeptidase-like_19_OS=Arabidopsis_thaliana_OX=3702_GN=SCPL19_PE=1_SV=1 |
| g26739.t1 | sp_P46032_PTR2_ARATH_Protein_NRT1/_PTR_FAMILY_8.3_OS=Arabidopsis_thaliana_OX=3702_GN=NPF8.3_PE=1_SV=1 |
| g26865.t1 | sp_Q39613_CYPH_CATRO_Peptidyl-prolyl_cis-trans_isomerase_OS=Catharanthus_roseus_OX=4058_GN=PCKR1_PE=1_SV=1 |
| g26992.t1 | sp_Q9ST02_NAATA_HORVU_Nicotianamine_aminotransferase_A_OS=Hordeum_vulgare_OX=4513_GN=naat-A_PE=1_SV=2 |
| g26993.t1 | sp_Q8C9S4_CC186_MOUSE_Coiled-coil_domain-containing_protein_186_OS=Mus_musculus_OX=10090_GN=Ccdc186_PE=1_SV=2 |
| g27078.t1 | sp_Q8VZ80_PLT5_ARATH_Polyol_transporter_5_OS=Arabidopsis_thaliana_OX=3702_GN=PLT5_PE=1_SV=2 |
| g27121.t1 | sp_P93844_PLDA2_ORYSJ_Phospholipase_D_alpha_2_OS=Oryza_sativa_subsp._japonica_OX=39947_GN=PLD2_PE=2_SV=2 |
| g27263.t1 | sp_P28814_BARW_HORVU_Barwin_OS=Hordeum_vulgare_OX=4513_PE=1_SV=1 |
| g27691.t1 | sp_Q84TH5_AB25G_ARATH_ABC_transporter_G_family_member_25_OS=Arabidopsis_thaliana_OX=3702_GN=ABCG25_PE=2_SV=1 |
| g27691.t1 | sp_Q84TH5_AB25G_ARATH_ABC_transporter_G_family_member_25_OS=Arabidopsis_thaliana_OX=3702_GN=ABCG25_PE=2_SV=1 |
| g27892.t1 | sp_Q9LJ97_LEA31_ARATH_Late_embryogenesis_abundant_protein_31_OS=Arabidopsis_thaliana_OX=3702_GN=RAB28_PE=1_SV=1 |
| g28048.t1 | sp_Q7Y0F2_NRX12_ORYSJ_Probable_nucleoredoxin_1-2_OS=Oryza_sativa_subsp._japonica_OX=39947_GN=Os03g0405900_PE=2_SV=1 |
| g28048.t1 | sp_Q7Y0F2_NRX12_ORYSJ_Probable_nucleoredoxin_1-2_OS=Oryza_sativa_subsp._japonica_OX=39947_GN=Os03g0405900_PE=2_SV=1 |
| g28114.t1 | sp_O80742_PUB19_ARATH_U-box_domain-containing_protein_19_OS=Arabidopsis_thaliana_OX=3702_GN=PUB19_PE=2_SV=1 |
| g28434.t1 | sp_Q8S397_NHX4_ARATH_Sodium/hydrogen_exchanger_4_OS=Arabidopsis_thaliana_OX=3702_GN=NHX4_PE=2_SV=2 |
| g28520.t1 | sp_Q9FRX6_AS1_ANTMA_Aureusidin_synthase_OS=Antirrhinum_majus_OX=4151_GN=AS1_PE=1_SV=1 |
| g28538.t1 | sp_Q2QXJ2_GL122_ORYSJ_Germin-like_protein_12-2_OS=Oryza_sativa_subsp._japonica_OX=39947_GN=Os12g0154800_PE=2_SV=1 |
| g28843.t2 | sp_P43284_TRPB2_MAIZE_Tryptophan_synthase_beta_chain_2,_chloroplastic_(Fragment)_OS=Zea_mays_OX=4577_GN=TSB2_PE=2_SV=1 |
| g29003.t1 | sp_Q0J8G8_MAD26_ORYSJ_MADS-box_transcription_factor_26_OS=Oryza_sativa_subsp._japonica_OX=39947_GN=MADS26_PE=2_SV=1 |
| g29068.t1 | sp_Q9FLI3_P2C75_ARATH_Probable_protein_phosphatase_2C_75_OS=Arabidopsis_thaliana_OX=3702_GN=AHG1_PE=1_SV=1 |
| g29097.t1 | sp_Q8NBL1_PGLT1_HUMAN_Protein_O-glucosyltransferase_1_OS=Homo_sapiens_OX=9606_GN=POGLUT1_PE=1_SV=1 |
| g29581.t1 | sp_Q9ST02_NAATA_HORVU_Nicotianamine_aminotransferase_A_OS=Hordeum_vulgare_OX=4513_GN=naat-A_PE=1_SV=2 |
| g29670.t1 | sp_Q8GT41_PLA1_PLAAC_Putative_invertase_inhibitor_OS=Platanus_acerifolia_OX=140101_PE=1_SV=1 |
| g29678.t1 | sp_Q9SYJ2_GPAT3_ARATH_Probable_glycerol-3-phosphate_acyltransferase_3_OS=Arabidopsis_thaliana_OX=3702_GN=GPAT3_PE=2_SV=1 |
| g29822.t1 | sp_Q8S3M3_NOIL_ELAOL_NOI-like_protein_OS=Elaeis_oleifera_OX=80265_PE=2_SV=1 |
| g30255.t1 | sp_P25776_ORYA_ORYSJ_Oryzain_alpha_chain_OS=Oryza_sativa_subsp._japonica_OX=39947_GN=Os04g0650000_PE=1_SV=2 |
| g30297.t1 | sp_Q8H1G5_REIL1_ARATH_Cytoplasmic_60S_subunit_biogenesis_factor_REI1_homolog_1_OS=Arabidopsis_thaliana_OX=3702_GN=REIL1_PE=1_SV=1 |
| g30342.t1 | sp_Q5Z6F5_P2C59_ORYSJ_Probable_protein_phosphatase_2C_59_OS=Oryza_sativa_subsp._japonica_OX=39947_GN=Os06g0698300_PE=2_SV=1 |
| g30393.t1 | sp_P32110_GSTX6_SOYBN_Probable_glutathione_S-transferase_OS=Glycine_max_OX=3847_GN=HSP26-A_PE=2_SV=1 |
| g30554.t1 | sp_P25765_CHI12_ORYSJ_Chitinase_12_OS=Oryza_sativa_subsp._japonica_OX=39947_GN=Cht12_PE=2_SV=2 |
| g30555.t1 | sp_Q10S65_NAC22_ORYSJ_NAC_domain-containing_protein_22_OS=Oryza_sativa_subsp._japonica_OX=39947_GN=NAC022_PE=2_SV=1 |
| g30555.t1 | sp_Q10S65_NAC22_ORYSJ_NAC_domain-containing_protein_22_OS=Oryza_sativa_subsp._japonica_OX=39947_GN=NAC022_PE=2_SV=1 |
| g30614.t1 | sp_Q54F46_WARA_DICDI_Homeobox_protein_Wariai_OS=Dictyostelium_discoideum_OX=44689_GN=warA_PE=2_SV=1 |
| g30704.t1 | sp_Q9LI74_CHUP1_ARATH_Protein_CHUP1,_chloroplastic_OS=Arabidopsis_thaliana_OX=3702_GN=CHUP1_PE=1_SV=1 |
| g31054.t1 | sp_Q9LUC5_C7A15_ARATH_Cytochrome_P450_72A15_OS=Arabidopsis_thaliana_OX=3702_GN=CYP72A15_PE=2_SV=1 |
| g31054.t2 | sp_Q9LUC5_C7A15_ARATH_Cytochrome_P450_72A15_OS=Arabidopsis_thaliana_OX=3702_GN=CYP72A15_PE=2_SV=1 |
| g31099.t1 | sp_Q9AT54_SCGT_TOBAC_Scopoletin_glucosyltransferase_OS=Nicotiana_tabacum_OX=4097_GN=TOGT1_PE=1_SV=1 |
| g31099.t1 | sp_Q9AT54_SCGT_TOBAC_Scopoletin_glucosyltransferase_OS=Nicotiana_tabacum_OX=4097_GN=TOGT1_PE=1_SV=1 |
| g31298.t1 | sp_Q9ST02_NAATA_HORVU_Nicotianamine_aminotransferase_A_OS=Hordeum_vulgare_OX=4513_GN=naat-A_PE=1_SV=2 |
| g31332.t1 | sp_O22808_LYK5_ARATH_Protein_LYK5_OS=Arabidopsis_thaliana_OX=3702_GN=LYK5_PE=2_SV=1 |
| g31332.t1 | sp_O22808_LYK5_ARATH_Protein_LYK5_OS=Arabidopsis_thaliana_OX=3702_GN=LYK5_PE=2_SV=1 |
| g31519.t1 | sp_C7G304_GOLS2_SOLLC_Galactinol_synthase_2_OS=Solanum_lycopersicum_OX=4081_GN=GOLS2_PE=2_SV=1 |
| g31522.t1 | sp_Q10MB4_MYB2_ORYSJ_Transcription_factor_MYB2_OS=Oryza_sativa_subsp._japonica_OX=39947_GN=MYB2_PE=2_SV=1 |
| g31542.t1 | sp_Q6Z4N4_40G3_ORYSJ_Ricin_B-like_lectin_R40G3_OS=Oryza_sativa_subsp._japonica_OX=39947_GN=R40G3_PE=2_SV=1 |
| g31543.t1 | sp_Q6Z4N6_40G2_ORYSJ_Ricin_B-like_lectin_R40G2_OS=Oryza_sativa_subsp._japonica_OX=39947_GN=R40G2_PE=2_SV=1 |
| g31819.t1 | sp_Q12PE6_TOLB_SHEDO_Tol-Pal_system_protein_TolB_OS=Shewanella_denitrificans_(strain_OS217_/_ATCC_BAA-1090_/_DSM_15013)_OX=318161_GN=tolB_PE=3_SV=1 |
| g32155.t2 | sp_O94260_G3BP_SCHPO_Putative_G3BP-like_protein_OS=Schizosaccharomyces_pombe_(strain_972_/_ATCC_24843)_OX=284812_GN=nxt3_PE=1_SV=1 |
| g32155.t2 | sp_O94260_G3BP_SCHPO_Putative_G3BP-like_protein_OS=Schizosaccharomyces_pombe_(strain_972_/_ATCC_24843)_OX=284812_GN=nxt3_PE=1_SV=1 |
| g32414.t1 | sp_P28522_RIPX_MAIZE_Ribosome-inactivating_protein_OS=Zea_mays_OX=4577_PE=1_SV=1 |
| g32573.t1 | sp_Q9FJU9_E1313_ARATH_Glucan_endo-1,3-beta-glucosidase_13_OS=Arabidopsis_thaliana_OX=3702_GN=At5g56590_PE=2_SV=1 |
| g32596.t1 | sp_Q8LPS2_ACD6_ARATH_Protein_ACCELERATED_CELL_DEATH_6_OS=Arabidopsis_thaliana_OX=3702_GN=ACD6_PE=1_SV=1 |
| g32620.t1 | sp_Q9FGG3_WTR45_ARATH_WAT1-related_protein_At5g64700_OS=Arabidopsis_thaliana_OX=3702_GN=At5g64700_PE=2_SV=1 |
| g32987.t1 | sp_Q9FKY4_PILS7_ARATH_Protein_PIN-LIKES_7_OS=Arabidopsis_thaliana_OX=3702_GN=PILS7_PE=2_SV=1 |
| g33102.t1 | sp_Q7XXN5_PHT3_ORYSJ_Putrescine_hydroxycinnamoyltransferase_3_OS=Oryza_sativa_subsp._japonica_OX=39947_GN=PHT3_PE=1_SV=1 |
| g33130.t1 | sp_Q651D5_PIP27_ORYSJ_Probable_aquaporin_PIP2-7_OS=Oryza_sativa_subsp._japonica_OX=39947_GN=PIP2-7_PE=2_SV=2 |
| g33158.t1 | sp_P42815_RNS3_ARATH_Ribonuclease_3_OS=Arabidopsis_thaliana_OX=3702_GN=RNS3_PE=2_SV=1 |
| g33369.t1 | sp_Q0J0G2_BGL32_ORYSJ_Beta-glucosidase_32_OS=Oryza_sativa_subsp._japonica_OX=39947_GN=BGLU32_PE=2_SV=2 |
| g33387.t1 | sp_Q9ZVN2_Y1457_ARATH_Acyltransferase-like_protein_At1g54570,_chloroplastic_OS=Arabidopsis_thaliana_OX=3702_GN=At1g54570_PE=1_SV=1 |
| g33488.t1 | sp_Q9SJA7_SOX_ARATH_Probable_sarcosine_oxidase_OS=Arabidopsis_thaliana_OX=3702_GN=At2g24580_PE=2_SV=1 |
| g33611.t1 | sp_Q8S9J6_ASPA_ARATH_Aspartyl_protease_family_protein_At5g10770_OS=Arabidopsis_thaliana_OX=3702_GN=At5g10770_PE=2_SV=1 |
| g33708.t1 | sp_Q9SJZ2_PER17_ARATH_Peroxidase_17_OS=Arabidopsis_thaliana_OX=3702_GN=PER17_PE=2_SV=1 |
| g33801.t1 | sp_Q9SJZ2_PER17_ARATH_Peroxidase_17_OS=Arabidopsis_thaliana_OX=3702_GN=PER17_PE=2_SV=1 |
| g33874.t1 | sp_Q9FMP8_RGAP6_ARATH_Rho_GTPase-activating_protein_6_OS=Arabidopsis_thaliana_OX=3702_GN=ROPGAP6_PE=2_SV=1 |
| g34016.t1 | sp_Q8VCZ2_TM45B_MOUSE_Transmembrane_protein_45B_OS=Mus_musculus_OX=10090_GN=Tmem45b_PE=1_SV=1 |
| g34052.t1 | sp_Q50EK4_C75A1_PINTA_Cytochrome_P450_750A1_OS=Pinus_taeda_OX=3352_GN=CYP750A1_PE=2_SV=1 |
| g34201.t1 | sp_Q6EPZ0_WRK62_ORYSJ_WRKY_transcription_factor_WRKY62_OS=Oryza_sativa_subsp._japonica_OX=39947_GN=WRKY62_PE=1_SV=1 |
| g34894.t1 | sp_P51140_DSH_DROME_Segment_polarity_protein_dishevelled_OS=Drosophila_melanogaster_OX=7227_GN=dsh_PE=1_SV=2 |
| g35053.t1 | sp_Q82PI3_KDPA_STRAW_Potassium-transporting_ATPase_potassium-binding_subunit_OS=Streptomyces_avermitilis_(strain_ATCC_31267_/_DSM_46492_/_JCM_5070_/_NBRC_14893_/_NCIMB_12804_/_NRRL_8165_/_MA-4680)_OX=227882_GN=kdpA_PE=3_SV=1 |
| g35421.t2 | sp_Q9LH39_PHSD_ARATH_Probable_polyamine_transporter_At3g19553_OS=Arabidopsis_thaliana_OX=3702_GN=At3g19553_PE=3_SV=1 |
| g35781.t1 | sp_Q8L8B8_LOG3_ARATH_Cytokinin_riboside_5'-monophosphate_phosphoribohydrolase_LOG3_OS=Arabidopsis_thaliana_OX=3702_GN=LOG3_PE=1_SV=1 |
| g35868.t1 | sp_Q9T0G0_HSD5_ARATH_11-beta-hydroxysteroid_dehydrogenase-like_5_OS=Arabidopsis_thaliana_OX=3702_GN=HSD5_PE=2_SV=1 |
| g35942.t1 | sp_Q9LUN4_CHX19_ARATH_Cation/H(+)_antiporter_19_OS=Arabidopsis_thaliana_OX=3702_GN=CHX19_PE=2_SV=1 |
| g36008.t1 | sp_Q84TB6_ADF3_ORYSJ_Actin-depolymerizing_factor_3_OS=Oryza_sativa_subsp._japonica_OX=39947_GN=ADF3_PE=1_SV=1 |
| g36301.t1 | sp_P42813_RNS1_ARATH_Ribonuclease_1_OS=Arabidopsis_thaliana_OX=3702_GN=RNS1_PE=1_SV=1 |
| g36590.t1 | sp_O24575_DCAM_MAIZE_S-adenosylmethionine_decarboxylase_proenzyme_OS=Zea_mays_OX=4577_GN=SAMDC_PE=2_SV=1 |
| g36911.t1 | sp_P13917_7SB1_SOYBN_Basic_7S_globulin_OS=Glycine_max_OX=3847_GN=BG_PE=1_SV=2 |
| g37003.t1 | sp_Q9FI25_BBE27_ARATH_Berberine_bridge_enzyme-like_27_OS=Arabidopsis_thaliana_OX=3702_GN=At5g44410_PE=2_SV=1 |
| g37022.t1 | sp_Q5RA52_PDP1_PONAB_[Pyruvate_dehydrogenase_[acetyl-transferring]]-phosphatase_1,_mitochondrial_OS=Pongo_abelii_OX=9601_GN=PDP1_PE=2_SV=1 |
| g37247.t2 | sp_P29518_BT1_MAIZE_Adenine_nucleotide_transporter_BT1,_chloroplastic/amyloplastic/mitochondrial_OS=Zea_mays_OX=4577_GN=BT1_PE=1_SV=1 |
| g37513.t1 | sp_C5X2M4_THI42_SORBI_Thiamine_thiazole_synthase_2,_chloroplastic_OS=Sorghum_bicolor_OX=4558_GN=THI1-2_PE=3_SV=1 |
| g37576.t1 | sp_Q84N34_HHP2_ARATH_Heptahelical_transmembrane_protein_2_OS=Arabidopsis_thaliana_OX=3702_GN=HHP2_PE=2_SV=1 |
| g37577.t1 | sp_Q67WR5_FCL2_ORYSJ_Putative_GDP-L-fucose_synthase_2_OS=Oryza_sativa_subsp._japonica_OX=39947_GN=Os06g0652300_PE=3_SV=1 |
| g37830.t1 | sp_Q9T076_ENL2_ARATH_Early_nodulin-like_protein_2_OS=Arabidopsis_thaliana_OX=3702_GN=At4g27520_PE=1_SV=1 |
| g37905.t1 | sp_P25776_ORYA_ORYSJ_Oryzain_alpha_chain_OS=Oryza_sativa_subsp._japonica_OX=39947_GN=Os04g0650000_PE=1_SV=2 |
| g37968.t1 | sp_Q0JGI1_SPL2_ORYSJ_Squamosa_promoter-binding-like_protein_2_OS=Oryza_sativa_subsp._japonica_OX=39947_GN=SPL2_PE=2_SV=2 |
| g37968.t1 | sp_Q0JGI1_SPL2_ORYSJ_Squamosa_promoter-binding-like_protein_2_OS=Oryza_sativa_subsp._japonica_OX=39947_GN=SPL2_PE=2_SV=2 |
| g38145.t1 | sp_Q03662_GSTX1_TOBAC_Probable_glutathione_S-transferase_OS=Nicotiana_tabacum_OX=4097_PE=2_SV=1 |
| g38248.t1 | sp_P37220_ASR3_SOLLC_Abscisic_stress-ripening_protein_3_OS=Solanum_lycopersicum_OX=4081_GN=ASR3_PE=3_SV=2 |
| g38283.t1 | sp_Q9STK5_PP269_ARATH_Pentatricopeptide_repeat-containing_protein_At3g48250,_chloroplastic_OS=Arabidopsis_thaliana_OX=3702_GN=At3g48250_PE=2_SV=1 |
| g38283.t1 | sp_Q9STK5_PP269_ARATH_Pentatricopeptide_repeat-containing_protein_At3g48250,_chloroplastic_OS=Arabidopsis_thaliana_OX=3702_GN=At3g48250_PE=2_SV=1 |
| g38297.t1 | sp_Q94CS9_TIP12_ORYSJ_Probable_aquaporin_TIP1-2_OS=Oryza_sativa_subsp._japonica_OX=39947_GN=TIP1-2_PE=2_SV=1 |
| g38328.t2 | sp_A2Y8B9_ANM7_ORYSI_Protein_arginine_N-methyltransferase_7_OS=Oryza_sativa_subsp._indica_OX=39946_GN=PRMT7_PE=3_SV=3 |
| g38465.t1 | sp_A2RVM0_TIC32_ARATH_Short-chain_dehydrogenase_TIC_32,_chloroplastic_OS=Arabidopsis_thaliana_OX=3702_GN=TIC32_PE=2_SV=1 |
| g38551.t1 | sp_P26301_ENO1_MAIZE_Enolase_1_OS=Zea_mays_OX=4577_GN=ENO1_PE=2_SV=1 |
| g38689.t1 | sp_O24645_HCBT1_DIACA_Anthranilate_N-benzoyltransferase_protein_1_OS=Dianthus_caryophyllus_OX=3570_GN=HCBT1_PE=1_SV=1 |
| g38962.t1 | sp_Q5SMM6_HCT4_ORYSJ_Hydroxycinnamoyltransferase_4_OS=Oryza_sativa_subsp._japonica_OX=39947_GN=HCT4_PE=1_SV=1 |
| g39214.t2 | sp_Q653P0_KOR1_ORYSJ_Potassium_channel_KOR1_OS=Oryza_sativa_subsp._japonica_OX=39947_GN=Os06g0250600_PE=2_SV=1 |
| g39238.t1 | sp_Q654V6_ODPA2_ORYSJ_Pyruvate_dehydrogenase_E1_component_subunit_alpha-2,_mitochondrial_OS=Oryza_sativa_subsp._japonica_OX=39947_GN=Os06g0246500_PE=2_SV=1 |
| g39264.t1 | sp_Q9SV61_XTH1_ARATH_Putative_xyloglucan_endotransglucosylase/hydrolase_protein_1_OS=Arabidopsis_thaliana_OX=3702_GN=XTH1_PE=3_SV=3 |
| g39330.t1 | sp_Q67WJ2_FTSH6_ORYSJ_ATP-dependent_zinc_metalloprotease_FTSH_6,_chloroplastic_OS=Oryza_sativa_subsp._japonica_OX=39947_GN=FTSH6_PE=3_SV=1 |
| g39330.t1 | sp_Q67WJ2_FTSH6_ORYSJ_ATP-dependent_zinc_metalloprotease_FTSH_6,_chloroplastic_OS=Oryza_sativa_subsp._japonica_OX=39947_GN=FTSH6_PE=3_SV=1 |
| g39339.t1 | sp_Q9CA57_GSTUA_ARATH_Glutathione_S-transferase_U10_OS=Arabidopsis_thaliana_OX=3702_GN=GSTU10_PE=2_SV=1 |
| g39590.t1 | sp_Q9SRT0_PUB9_ARATH_U-box_domain-containing_protein_9_OS=Arabidopsis_thaliana_OX=3702_GN=PUB9_PE=1_SV=1 |
| g39632.t1 | sp_Q9MAT6_HMG15_ARATH_High_mobility_group_B_protein_15_OS=Arabidopsis_thaliana_OX=3702_GN=HMGB15_PE=2_SV=1 |
| g39675.t1 | sp_P24805_TSJT1_TOBAC_Stem-specific_protein_TSJT1_OS=Nicotiana_tabacum_OX=4097_GN=TSJT1_PE=2_SV=1 |
| g39865.t1 | sp_Q84QK0_OPR1_ORYSJ_12-oxophytodienoate_reductase_1_OS=Oryza_sativa_subsp._japonica_OX=39947_GN=OPR1_PE=2_SV=1 |
| g40078.t1 | sp_P82977_ICIW_WHEAT_Subtilisin-chymotrypsin_inhibitor_WSCI_OS=Triticum_aestivum_OX=4565_PE=1_SV=2 |
| g40141.t1 | sp_Q653P0_KOR1_ORYSJ_Potassium_channel_KOR1_OS=Oryza_sativa_subsp._japonica_OX=39947_GN=Os06g0250600_PE=2_SV=1 |
| g40141.t1 | sp_Q653P0_KOR1_ORYSJ_Potassium_channel_KOR1_OS=Oryza_sativa_subsp._japonica_OX=39947_GN=Os06g0250600_PE=2_SV=1 |
| g40210.t1 | sp_Q01MW8_PHT14_ORYSI_Probable_inorganic_phosphate_transporter_1-4_OS=Oryza_sativa_subsp._indica_OX=39946_GN=PHT1-4_PE=2_SV=2 |
| g40217.t1 | sp_Q8H6H2_PHT14_ORYSJ_Probable_inorganic_phosphate_transporter_1-4_OS=Oryza_sativa_subsp._japonica_OX=39947_GN=PHT1-4_PE=2_SV=1 |
| g40373.t1 | sp_Q84QK0_OPR1_ORYSJ_12-oxophytodienoate_reductase_1_OS=Oryza_sativa_subsp._japonica_OX=39947_GN=OPR1_PE=2_SV=1 |
| g40833.t1 | sp_P09444_LEA34_GOSHI_Late_embryogenesis_abundant_protein_D-34_OS=Gossypium_hirsutum_OX=3635_PE=3_SV=1 |
| g40900.t1 | sp_P13917_7SB1_SOYBN_Basic_7S_globulin_OS=Glycine_max_OX=3847_GN=BG_PE=1_SV=2 |
| g41037.t1 | sp_Q9FI25_BBE27_ARATH_Berberine_bridge_enzyme-like_27_OS=Arabidopsis_thaliana_OX=3702_GN=At5g44410_PE=2_SV=1 |
| g41056.t1 | sp_P0CB22_ATX2_ARATH_Histone-lysine_N-methyltransferase_ATX2_OS=Arabidopsis_thaliana_OX=3702_GN=ATX2_PE=2_SV=1 |
| g41213.t1 | sp_Q9LV58_MBF1C_ARATH_Multiprotein-bridging_factor_1c_OS=Arabidopsis_thaliana_OX=3702_GN=MBF1C_PE=1_SV=1 |
| g41524.t1 | sp_Q9C942_CSE_ARATH_Caffeoylshikimate_esterase_OS=Arabidopsis_thaliana_OX=3702_GN=CSE_PE=1_SV=1 |
| g41645.t1 | sp_Q9ZVJ6_ANXD4_ARATH_Annexin_D4_OS=Arabidopsis_thaliana_OX=3702_GN=ANN4_PE=1_SV=1 |
| g41924.t1 | sp_Q8W4A7_GUX3_ARATH_Putative_UDP-glucuronate:xylan_alpha-glucuronosyltransferase_3_OS=Arabidopsis_thaliana_OX=3702_GN=GUX3_PE=2_SV=1 |
| g42052.t1 | sp_Q9ZUX1_C94C1_ARATH_Cytochrome_P450_94C1_OS=Arabidopsis_thaliana_OX=3702_GN=CYP94C1_PE=1_SV=1 |
| g42096.t1 | sp_Q8LGG8_USPAL_ARATH_Universal_stress_protein_A-like_protein_OS=Arabidopsis_thaliana_OX=3702_GN=At3g01520_PE=1_SV=2 |
| g42207.t1 | sp_O26373_Y273_METTH_UPF0098_protein_MTH_273_OS=Methanothermobacter_thermautotrophicus_(strain_ATCC_29096_/_DSM_1053_/_JCM_10044_/_NBRC_100330_/_Delta_H)_OX=187420_GN=MTH_273_PE=3_SV=1 |
| g42231.t1 | sp_Q9FLB1_PYL5_ARATH_Abscisic_acid_receptor_PYL5_OS=Arabidopsis_thaliana_OX=3702_GN=PYL5_PE=1_SV=1 |
| g42397.t1 | sp_Q9FL41_WTR38_ARATH_WAT1-related_protein_At5g07050_OS=Arabidopsis_thaliana_OX=3702_GN=At5g07050_PE=2_SV=1 |
| g42762.t1 | sp_Q42376_LEA3_MAIZE_Late_embryogenesis_abundant_protein,_group_3_OS=Zea_mays_OX=4577_GN=MGL3_PE=2_SV=1 |
| g42803.t1 | sp_Q944H0_PEAM2_ARATH_Phosphomethylethanolamine_N-methyltransferase_OS=Arabidopsis_thaliana_OX=3702_GN=NMT2_PE=1_SV=2 |
| g42803.t1 | sp_Q944H0_PEAM2_ARATH_Phosphomethylethanolamine_N-methyltransferase_OS=Arabidopsis_thaliana_OX=3702_GN=NMT2_PE=1_SV=2 |
| g42890.t1 | sp_Q9LTW4_NANA_ARATH_Aspartic_proteinase_NANA,_chloroplast_OS=Arabidopsis_thaliana_OX=3702_GN=NANA_PE=1_SV=1 |
| g42892.t1 | sp_Q9LTW4_NANA_ARATH_Aspartic_proteinase_NANA,_chloroplast_OS=Arabidopsis_thaliana_OX=3702_GN=NANA_PE=1_SV=1 |
| g42915.t1 | sp_P04518_RDRP_CARMV_RNA-directed_RNA_polymerase_OS=Carnation_mottle_virus_OX=11986_GN=ORF1_PE=4_SV=2 |
| g42941.t1 | sp_Q9D832_DNJB4_MOUSE_DnaJ_homolog_subfamily_B_member_4_OS=Mus_musculus_OX=10090_GN=Dnajb4_PE=1_SV=1 |
| g43000.t1 | sp_O49627_ISU1_ARATH_Iron-sulfur_cluster_assembly_protein_1_OS=Arabidopsis_thaliana_OX=3702_GN=ISU1_PE=2_SV=1 |
| g43036.t1 | sp_Q65XK7_P2C51_ORYSJ_Protein_phosphatase_2C_51_OS=Oryza_sativa_subsp._japonica_OX=39947_GN=PP2C51_PE=1_SV=1 |
| g43056.t1 | sp_Q6F357_PLA7_ORYSJ_Phospholipase_A1-II_7_OS=Oryza_sativa_subsp._japonica_OX=39947_GN=Os05g0574100_PE=3_SV=1 |
| g43199.t1 | sp_Q8VZG2_HSR4_ARATH_Protein_HYPER-SENSITIVITY-RELATED_4_OS=Arabidopsis_thaliana_OX=3702_GN=HSR4_PE=2_SV=1 |
| g43274.t1 | sp_F4JJ23_NEN4_ARATH_Protein_NEN4_OS=Arabidopsis_thaliana_OX=3702_GN=NEN4_PE=2_SV=1 |
| g43321.t1 | sp_P15490_VSPA_SOYBN_Stem_28_kDa_glycoprotein_OS=Glycine_max_OX=3847_GN=VSPA_PE=2_SV=1 |
| g43322.t1 | sp_P15490_VSPA_SOYBN_Stem_28_kDa_glycoprotein_OS=Glycine_max_OX=3847_GN=VSPA_PE=2_SV=1 |
| g43386.t1 | sp_Q6AT12_G3OX1_ORYSJ_Gibberellin_3-beta-dioxygenase_1_OS=Oryza_sativa_subsp._japonica_OX=39947_GN=GA3OX1_PE=1_SV=1 |
| g43402.t1 | sp_Q9SUV1_BRT1_ARATH_Adenine_nucleotide_transporter_BT1,_chloroplastic/mitochondrial_OS=Arabidopsis_thaliana_OX=3702_GN=BT1_PE=1_SV=1 |
| g43661.t1 | sp_Q6DR20_FB110_ARATH_F-box_protein_At2g17036_OS=Arabidopsis_thaliana_OX=3702_GN=At2g17036_PE=2_SV=1 |
| g43770.t1 | sp_A2AG06_MEIOC_MOUSE_Meiosis-specific_coiled-coil_domain-containing_protein_MEIOC_OS=Mus_musculus_OX=10090_GN=Meioc_PE=1_SV=1 |
| g43946.t1 | sp_Q9FLV9_SLAH3_ARATH_S-type_anion_channel_SLAH3_OS=Arabidopsis_thaliana_OX=3702_GN=SLAH3_PE=1_SV=1 |
| g43958.t1 | sp_Q9FYE2_PKS4_ARATH_Protein_PHYTOCHROME_KINASE_SUBSTRATE_4_OS=Arabidopsis_thaliana_OX=3702_GN=PKS4_PE=1_SV=1 |
| g43969.t1 | sp_Q07469_BSPA_POPDE_Bark_storage_protein_A_OS=Populus_deltoides_OX=3696_GN=BSPA_PE=2_SV=1 |
| g44010.t1 | sp_Q1T7C2_C7111_SOLLC_Cytochrome_P450_710A11_OS=Solanum_lycopersicum_OX=4081_GN=CYP710A11_PE=1_SV=1 |
| g44142.t1 | sp_P99177_CSD_STAAN_Probable_cysteine_desulfurase_OS=Staphylococcus_aureus_(strain_N315)_OX=158879_GN=csd_PE=1_SV=1 |
| g44242.t1 | sp_P19656_NLTP_MAIZE_Non-specific_lipid-transfer_protein_OS=Zea_mays_OX=4577_PE=1_SV=1 |
| g44263.t1 | sp_Q9ZVI6_LOR8_ARATH_Protein_LURP-one-related_8_OS=Arabidopsis_thaliana_OX=3702_GN=At2g38640_PE=2_SV=1 |
| g44291.t1 | sp_Q8W488_DTX21_ARATH_Protein_DETOXIFICATION_21_OS=Arabidopsis_thaliana_OX=3702_GN=DTX21_PE=1_SV=1 |
| g44291.t1 | sp_Q8W488_DTX21_ARATH_Protein_DETOXIFICATION_21_OS=Arabidopsis_thaliana_OX=3702_GN=DTX21_PE=1_SV=1 |
| g44412.t1 | sp_Q73YJ7_LIPB_MYCPA_Octanoyltransferase_OS=Mycobacterium_paratuberculosis_(strain_ATCC_BAA-968_/_K-10)_OX=262316_GN=lipB_PE=3_SV=1 |
| g44436.t1 | sp_Q8S340_PPA22_ARATH_Purple_acid_phosphatase_22_OS=Arabidopsis_thaliana_OX=3702_GN=PAP22_PE=2_SV=1 |
| g44536.t1 | sp_Q70II3_EF110_ARATH_Ethylene-responsive_transcription_factor_ERF110_OS=Arabidopsis_thaliana_OX=3702_GN=ERF110_PE=2_SV=2 |
| g44602.t1 | sp_Q9C942_CSE_ARATH_Caffeoylshikimate_esterase_OS=Arabidopsis_thaliana_OX=3702_GN=CSE_PE=1_SV=1 |
| g44701.t1 | sp_P83827_TRPD_THETH_Anthranilate_phosphoribosyltransferase_OS=Thermus_thermophilus_OX=274_GN=trpD_PE=1_SV=1 |
| g44745.t1 | sp_Q41359_PR1_SAMNI_Pathogenesis-related_protein_PR-1_type_OS=Sambucus_nigra_OX=4202_PE=2_SV=1 |
| g45240.t1 | sp_C5D3D3_TRPA_GEOSW_Tryptophan_synthase_alpha_chain_OS=Geobacillus_sp._(strain_WCH70)_OX=471223_GN=trpA_PE=3_SV=1 |
| g45242.t1 | sp_Q7T2V2_ST1S3_DANRE_Cytosolic_sulfotransferase_3_OS=Danio_rerio_OX=7955_GN=sult1st3_PE=1_SV=1 |
| g45375.t1 | sp_Q8VZ80_PLT5_ARATH_Polyol_transporter_5_OS=Arabidopsis_thaliana_OX=3702_GN=PLT5_PE=1_SV=2 |
| g45375.t1 | sp_Q8VZ80_PLT5_ARATH_Polyol_transporter_5_OS=Arabidopsis_thaliana_OX=3702_GN=PLT5_PE=1_SV=2 |
| g45432.t1 | sp_Q9ZQ89_UPS2_ARATH_Ureide_permease_2_OS=Arabidopsis_thaliana_OX=3702_GN=UPS2_PE=1_SV=2 |
| g45442.t1 | sp_Q8S9J6_ASPA_ARATH_Aspartyl_protease_family_protein_At5g10770_OS=Arabidopsis_thaliana_OX=3702_GN=At5g10770_PE=2_SV=1 |
| g45520.t1 | sp_Q8VZ80_PLT5_ARATH_Polyol_transporter_5_OS=Arabidopsis_thaliana_OX=3702_GN=PLT5_PE=1_SV=2 |
| g45603.t1 | sp_O80763_NRX1_ARATH_Probable_nucleoredoxin_1_OS=Arabidopsis_thaliana_OX=3702_GN=At1g60420_PE=1_SV=1 |
| g46038.t1 | sp_Q75BE7_ASA1_ASHGO_ASTRA-associated_protein_1_OS=Ashbya_gossypii_(strain_ATCC_10895_/_CBS_109.51_/_FGSC_9923_/_NRRL_Y-1056)_OX=284811_GN=ASA1_PE=3_SV=1 |
| g46137.t1 | sp_Q10PE7_DMAS1_ORYSJ_Deoxymugineic_acid_synthase_1_OS=Oryza_sativa_subsp._japonica_OX=39947_GN=DMAS1_PE=1_SV=1 |
| g46293.t1 | sp_Q69T31_CINV1_ORYSJ_Cytosolic_invertase_1_OS=Oryza_sativa_subsp._japonica_OX=39947_GN=CINV1_PE=1_SV=1 |
| g46326.t1 | sp_Q7G1L2_RAP26_ARATH_Ethylene-responsive_transcription_factor_RAP2-6_OS=Arabidopsis_thaliana_OX=3702_GN=RAP2-6_PE=2_SV=2 |
| g46479.t1 | sp_Q3UDW8_HGNAT_MOUSE_Heparan-alpha-glucosaminide_N-acetyltransferase_OS=Mus_musculus_OX=10090_GN=Hgsnat_PE=1_SV=2 |
| g46625.t1 | sp_Q5FH27_ASSY_EHRRG_Argininosuccinate_synthase_OS=Ehrlichia_ruminantium_(strain_Gardel)_OX=302409_GN=argG_PE=3_SV=1 |
| g46631.t1 | sp_O65351_SBT17_ARATH_Subtilisin-like_protease_SBT1.7_OS=Arabidopsis_thaliana_OX=3702_GN=SBT1.7_PE=1_SV=1 |
| g46713.t1 | sp_Q84WT5_XYN5L_ARATH_Endo-1,4-beta-xylanase_5-like_OS=Arabidopsis_thaliana_OX=3702_GN=At4g33820_PE=2_SV=1 |
| g46809.t1 | sp_Q91VC0_DCTP1_RAT_dCTP_pyrophosphatase_1_OS=Rattus_norvegicus_OX=10116_GN=Dctpp1_PE=2_SV=1 |
| g46980.t1 | sp_Q01MW8_PHT14_ORYSI_Probable_inorganic_phosphate_transporter_1-4_OS=Oryza_sativa_subsp._indica_OX=39946_GN=PHT1-4_PE=2_SV=2 |
| g47019.t1 | sp_M0Y4P1_UGT13_HORVV_UDP-glucosyltransferase_UGT13248_OS=Hordeum_vulgare_subsp._vulgare_OX=112509_PE=2_SV=1 |
